# Shared evolutionary processes shape landscapes of genomic variation in the great apes

**DOI:** 10.1101/2023.02.07.527547

**Authors:** Murillo F. Rodrigues, Andrew D. Kern, Peter L. Ralph

## Abstract

For at least the past five decades population genetics, as a field, has worked to describe the precise balance of forces that shape patterns of variation in genomes. The problem is challenging because modelling the interactions between evolutionary processes is difficult, and different processes can impact genetic variation in similar ways. In this paper, we describe how diversity and divergence between closely related species change with time, using correlations between landscapes of genetic variation as a tool to understand the interplay between evolutionary processes. We find strong correlations between landscapes of diversity and divergence in a well sampled set of great ape genomes, and explore how various processes such as incomplete lineage sorting, mutation rate variation, GC-biased gene conversion and selection contribute to these correlations. Through highly realistic, chromosome-scale, forward-in-time simulations we show that the landscapes of diversity and divergence in the great apes are too well correlated to be explained via strictly neutral processes alone. Our best fitting simulation includes both deleterious and beneficial mutations in functional portions of the genome, in which 9% of fixations within those regions is driven by positive selection. This study provides a framework for modelling genetic variation in closely related species, an approach which can shed light on the complex balance of forces that have shaped genetic variation.

## 1 Introduction

Genetic variation is determined by the combined action of mutation, demographic processes, recombination and natural selection. However, there is still no consensus on the relative contributions of these processes and their interactions in shaping patterns of genetic variation. Two major open questions are: how does the influence of selection compare to other processes? And, to what degree is genetic variation influenced by beneficial versus deleterious mutations?

Genetic variation can be measured within a species or between species with two related metrics: within-species genetic diversity and between-species genetic divergence. Both can be estimated with genetic data by computing the per site average number of differences between pairs of samples within a species or between two species. and these are estimates of the mean time to coalescence. (Note that we do not discuss *relative divergence*, which is often measured using *F_ST_*.) Evolutionary processes impact diversity and divergence in different ways, so the relationship between these carries information regarding these processes.

Natural selection directly impacts genetic diversity because it can reduce the frequencies of alleles that are deleterious (negative selection) or increase those of beneficial alleles (positive selection). Selection can also directly affect between-species genetic divergence. Deleterious alleles are more likely to be lost from the population, thus reducing divergence at the affected sites. On the other hand, beneficial alleles have a higher probability of fixation. This leads to an increase in the rate of substitution at sites under positive selection that in turn increases divergence between species. Thus, contrasting patterns of diversity and divergence at the same time can help disentangle between modes of selection (Hudson et al., 1987). Indeed, perhaps the most widely used test for detecting adaptive evolution, the McDonald-Kreitman test, compares diversity and divergence contrasted between neutral (e.g., synonymous) and functional (e.g., non-synonymous) site classes (McDonald & Kreitman, 1991). This test and its extensions have been applied to a myriad of taxa, and it has become clear that a substantial proportion of amino acid substitutions are driven by positive selection in a number of taxa (Galtier, 2016; Ingvarsson, 2010; Slotte, 2014; N. Smith & Eyre-Walker, 2002).

Selection also disturbs genetic variation at nearby locations on the genome, and this indirect effect of selection on diversity is called “linked selection”. Linked selection can be caused by at least two familiar mechanisms: genetic hitchhiking and background selection. Under genetic hitchhiking, as a beneficial mutation quickly increases in frequency in a population, its nearby genetic background is carried along, causing local reductions in levels of genetic diversity. The size of the region affected by the sweep depends on the strength of selection, which determines how fast fixation happens, and the crossover rate, because recombination allows linked sites to escape from the haplotype carrying the beneficial mutation (Kaplan et al., 1989; Maynard Smith & Haigh, 1974). Under background selection, neutral variation linked to deleterious mutations is removed from the population unless, as before, focal lineages escape via recombination (Charlesworth et al., 1993). Both of these processes leave similar footprints on patterns of within-species genetic diversity, and so attempts to determine the contributions of positive and negative selection in shaping levels of genetic variation genome-wide have proven to be difficult (Andolfatto, 2001; Y. Kim & Stephan, 2000), although the processes seem separable more locally (Schrider, 2020; Schrider & Kern, 2017). Importantly, linked selection does not affect between-species genetic divergence as strongly, as a beneficial or deleterious mutation does not affect the substitution rate of linked, neutral mutations (Birky & Walsh, 1988) (although it does affect divergence through ancestral levels of polymorphism (Begun et al., 2007; Phung et al., 2016)).

The effects of linked selection in shaping genetic variation are pervasive across genomes (Begun & Aquadro, 1992; Cai et al., 2009; Corbett-Detig et al., 2015; Lohmueller et al., 2011; Murphy et al., 2022). For example, dips in nucleotide diversity surrounding functional substitutions have been uncovered in many taxa, such as fruit flies (Kern et al., 2002; Sattath et al., 2011), rodents (Halligan et al., 2013), *Capsella* (Williamson et al., 2014) and maize (Beissinger et al., 2016). In *Drosophila melanogaster*, levels of synonymous diversity (which is putatively neutral) and amino acid divergence are negatively correlated (Andolfatto, 2007; Macpherson et al., 2007); positive selection can cause such a pattern if beneficial amino acid mutations are fixing and as they do reducing levels of linked neutral variation via selective sweeps. In contrast in humans, levels of synonymous diversity are roughly the same near amino acid substitutions and synonymous substitutions, suggesting recent, recent fixations at amino acids sites may not be the result of strongly beneficial alleles (Hernandez et al., 2011; Lohmueller et al., 2011). However, in the human genome, amino acid substitutions tend to be located in regions of lower constraint than synonymous substitutions, implying that the signal of positive selection may be confounded by the effects of background selection (Enard et al., 2014).

Two major challenges remain in the way of a fuller characterization of the effects of selection on genetic variation: (i) it is hard to model interactions between evolutionary processes (e.g., sweeps within highly constrained regions), and (ii) model identifiability is challenging for some summaries of the data (e.g., sweeps and background selection may impact diversity in similar ways). Recent computational advances have made it possible for us to move from simpler backwards-in-time coalescent models (Hudson, 1983) to more complex and computationally demanding forward-in-time simulations, and these have provided a route to studying these hard to model interactions between evolutionary processes across multiple sites (Haller & Messer, 2019; Haller et al., 2019; Kelleher et al., 2016). Simulation-based inference can then allow us to better describe the roles of different modes of selection and other processes in shaping genomic variation. However, the problem of identifying features of the data that are informative of the strength and mode of selection still remains.

One promising approach might be to compare patterns of genetic variation in multiple species jointly as each species can be thought of as semi-independent realizations of the same evolutionary processes (c.f. Won & Hey, 2005). In speciation genomics studies, it is common to visualize large scale patterns of genetic variation along chromosomes (so-called landscapes of diversity and divergence), which may contain substantial information to help us disentangle evolutionary processes. Earlier empirical surveys have focused on the identification of regions of accentuated relative divergence between populations (Cruickshank & Hahn, 2014; Harr, 2006; Turner et al., 2005), although patches of increased divergence can be the result of myriad forces besides reproductive isolation and adaptation. Recent comparative studies have found that landscapes of diversity are highly correlated between related groups of species, such as *Ficedula* flycatchers (Burri et al., 2015; Ellegren et al., 2012), warblers (Irwin et al., 2016), stonechats (van Doren et al., 2017), hummingbirds (Battey, 2020), monkeyflowers (Stankowski et al., 2019) and *Populus* (Wang et al., 2020). Burri (2017) proposed that we could capitalize on correlated genomic landscapes to study the interplay between different forms of selection and other evolutionary processes. Neutral processes, such as retained ancestral diversity (i.e., incomplete lineage sorting) or migration, could potentially produce significant correlations in levels of diversity across species, however strong correlations have been observed among taxa with long divergence times and without evidence of gene flow. For example, Stankowski et al. (2019) found that landscapes of diversity and divergence are highly correlated across a radiation of monkeyflowers which spans one million year (or about 10*N_e_*generations, where *N_e_*is the effective population size), far longer than the time scale on which we expect to see effects of ancestral variation and incomplete lineage sorting (since the coalescent timescale spans just a few multiples of *N_e_*). However, a shared process that independently occurs in the branches of a group of species could maintain correlations over long timescales. For example, if two species’ physical arrangement of functional elements and local recombination rates are similar, the direct and indirect effects of selection could make it so that peaks and valleys on the landscape of diversity are similar, maintaining correlation between their landscapes over evolutionary time (Burri, 2017; Delmore et al., 2018). Further, if mutational processes are heterogeneous across the genome in a manner that is shared among species, then correlated landscapes of diversity could be created through mutational variation as well.

Here, we aim to (i) describe whether and in what ways landscapes of within-species diversity and between-species divergence are correlated, and (ii) to tease apart the relative roles of positive and negative selection and other processes (e.g., ancestral variation, mutation rate variation) in shaping patterns of genetic variation. To understand processes driving these correlations, we employ highly realistic, chromosome-scale, forward-in-time simulations, since analytical predictions are not available. We use the great apes as a system to investigate correlated patterns of genetic variation because there is high quality population genomic data for all species (Prado-Martinez et al., 2013), the clade is about 12 million years old or 60*N_e_* generations (but there have not been many chromosomal arrangements (Jauch et al., 1992)), and lastly the landscapes of gene density, recombination rate and mutation rate are roughly conserved (Kronenberg et al., 2018; Stevison et al., 2016). Our study demonstrates that correlated landscapes can be useful in distinguishing between modes of selection and the balance of direct and linked selection shaping genomic variation.

## 2 Methods

### 2.1 Genomic data

We retrieved SNP calls for ten great ape populations made on high coverage (25) shortread sequencing data from the Great Ape Genome Project (Prado-Martinez et al., 2013), mapped onto the human reference genome (NCBI36/hg18). We analyzed 86 individuals divided into the following populations: human (*n* = 9 samples), bonobo (*n* = 13), Nigeria-Cameroon chimpanzee (*n* = 10), eastern chimpanzee (*n* = 6), central chimpanzee (*n* = 4), western chimpanzee (*n* = 4), eastern lowland gorilla (*n* = 3), western gorilla (*n* = 27), Sumatran orangutan (*n* = 5), Bornean orangutan (*n* = 5) (we excluded two samples from the original dataset: the Cross River gorilla and the chimpanzee hybrid). Prado-Martinez et al. (2013) applied several quality filters to the SNP calls (see Section 2.1 of their Supplementary Information) and, for each species, identified the genomic regions in which it would be unreliable to call SNPs (uncallable regions). For our downstream analyses, we only considered sites which were callable in all populations.

We calculated nucleotide diversity and divergence (*d_XY_*) in non-overlapping 1Mb windows using `scikit-allel` (Miles et al., 2020). Windows in which there were less than 40% callable sites were not used in any of the analyses. For example, this yielded 129 (out of 132) 1Mb windows in chromosome 12 in which 75% of the sites were callable on average.

To tease apart the effects of GC-biased gene conversion (gBGC), we decomposed diversity and divergence by allelic states. gBGC is expected to affect weak bases (A or T) which are disfavored when in heterozygotes which also carry a strong base (G or C). Thus, one way understand the effects of gBGC is by comparing sites which were weak to those that were strong in the ancestor (ancestrally strong alleles are not affected by gBGC, but ancestrally weak alleles can be). We assumed that the state in the ancestor of the great apes to be the state seen in rhesus macaques (genome version RheMac2) — sites without enough information in RheMac2 were excluded. Then, we computed divergence only considering sites which were ancestrally weak or ancestrally strong (Figure S4). This approach has two major drawbacks: (i) many of the sites cannot be used because they are missing in RheMac2 and (ii) sites can be mispolarized. Thus, we came up with a second approach to tease apart the effects of gBGC on correlations between genomic landscapes. When comparing two landscapes of divergence (which encompass four species), we can classify each site by the change in state that happened without needing to polarize mutations by looking at the ancestor. For example, if we have allelic states for four species and we see A-A-T-T as the configuration of alleles at a particular site, we know that there must have been one mutation which changed the state from a weak base to another weak base (W-W). On the other hand, if we see A-G-A-A there must have been one mutation from weak to strong (W-S) (or vice-versa). Sites with multiple mutations (e.g., A-G-G-C) were removed from the analyses. Sites that did not change from W to S (or vice-versa) are not expected to be affected by gBGC, and we refer to these as W-W or S-S mutations (Figure 6A). Sites where there may have been a weak to strong change (W-S mutations) may be affected by gBGC (Figure 6B). We only considered windows with at least 5% of callable sites in these analyses.

### 2.2 Simulations

We implemented forward-in-time Wright-Fisher simulations of the entire evolutionary history of the great apes using `SLiM` (Haller & Messer, 2019; Haller et al., 2019). Each branch in the great apes’ tree was simulated as a single population with constant size (Figure 1). Population splits occurred in a single generation, and there was no contact between populations post-split. Population sizes and split times were taken from the estimates in Prado-Martinez et al. (2013). Across all our simulations, we simulated crossover events occurred with the sex-averaged rates from the deCODE genetic map (in assembly NCBI36/hg18 coordinates) (Kong et al., 2002). We then computed diversity and divergence in the same windows used for the real data using tskit (Kelleher et al., 2018; Ralph et al., 2020).

**Figure 1:**
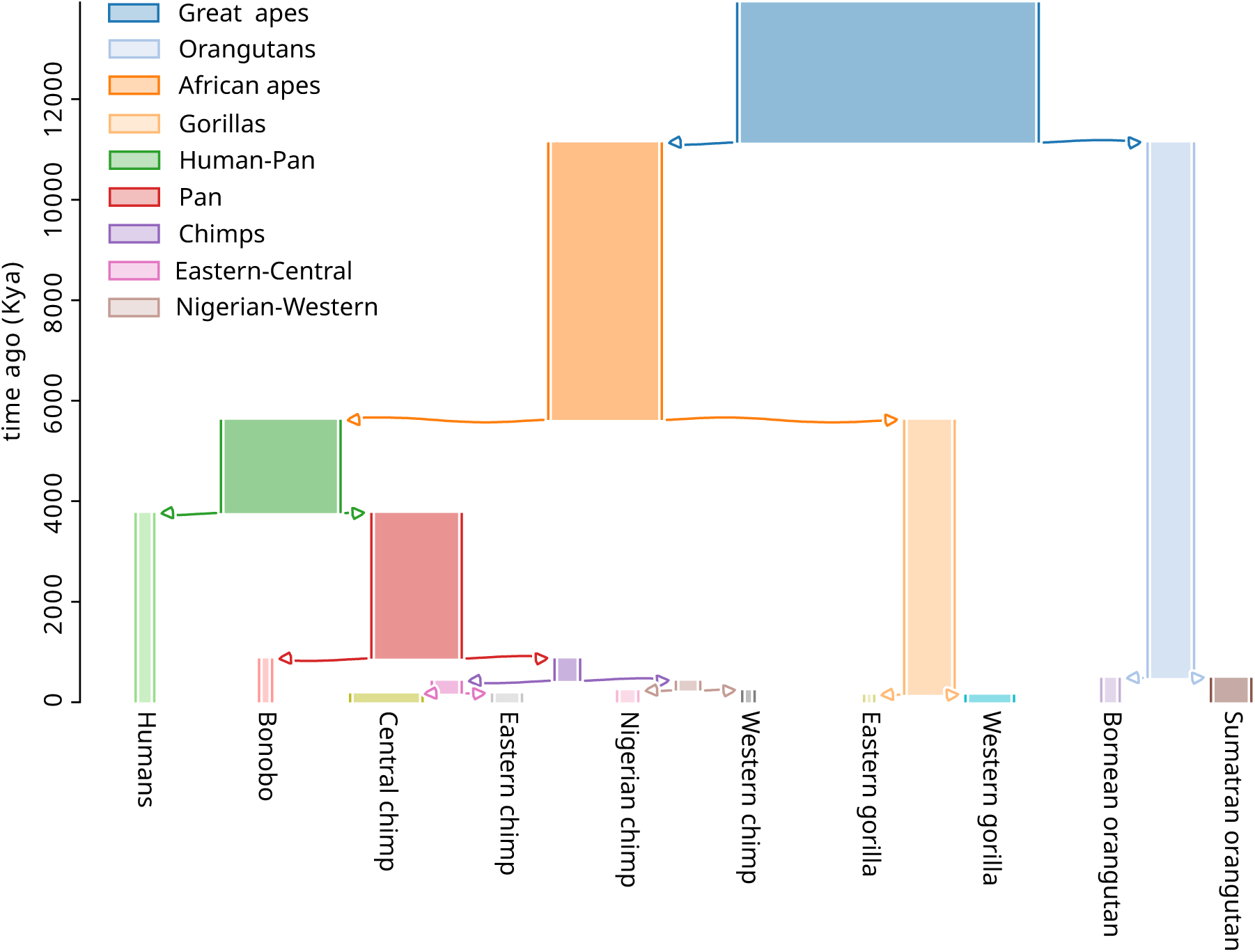
Range of parameters explored in the simulations. Arrows indicate population splits. Branch widths are proportional to population size. For example, the population size was 125, 089 for the great apes branch and 7, 672 for the humans branch. Figure was produced using *demesdraw* (Gower et al., 2022).

To improve run time, we simulated sister branches in parallel and recorded the final genealogies as tree sequences (Kelleher et al., 2016). Further, neutral mutations were not simulated with `SLiM` and were added after the fact with `msprime`. The resulting tree sequences were later joined and recapitated (i.e., we simulated genetic variation in the ancestor of all great apes using the coalescent) using `msprime, tskit` and `pyslim` (Kelleher et al., 2016, 2018; Rodrigues & Ralph, 2021). Despite our efforts to improve run time, our simulations of the entire history of the great apes were still incredibly costly (taking over a month to complete in many instances).

In our neutral simulations, we assumed that neutral mutations occurred at a rate of 2 10*^−^*^8^ new mutations per generation per site (Scally & Durbin, 2012), uniformly across the entire chromosome. To understand the effects of natural selection on landscapes, we simulated beneficial and deleterious mutations only within exons, assuming that the locations of exons were shared across all great apes (Kronenberg et al., 2018) and using exon annotations from the human reference genome NCBI36/hg18. We varied the proportions of neutral, beneficial and deleterious mutations within exons, but the distribution of fitness effects (DFE) for both deleterious and beneficial mutations were shared across all apes. The DFE for deleterious mutations was gamma-distributed with a fixed shape *α* and scale as estimated in Castellano et al. (2019), and the DFE for beneficial mutations followed an exponential distribution (Orr, 2003).

By default, we added neutral mutations to the simulated genealogies with `msprime` so that the total mutation rate (of neutral plus non-neutral mutations, if any) was constant along the genome. In addition, to simulate local variation in mutation rates along the chromosome, we selected three simulated genealogies (the fully neutral simulation, one with deleterious mutations and one with both beneficial and deleterious mutations) to add neutral mutations in a way that resulted in varying levels of neural mutation rate variation along the chromosome. To do this, we built mutation rate maps by sampling mutation rates for each 1Mb window independently from a normal distribution with mean 2 10*^−^*^8^ and standard deviation chosen from *σ/*2 10*^−^*^8^ = 0.005, 0.007, 0.011, 0.016, 0.023, 0.033, 0.048, 0.070.0.103, 0.150. In simulations with non-neutral mutations, we subtracted the non-neutral mutation rate from the respective window mutation rate for the intervals that intersected with exons. In total, we explored 56 different parameter combinations with all the different simulations (see Table 1 and Section 4.1 for the parameter space). The code used to produce the simulations can be found at https://github.com/kr-colab/greatapes sims.

**Table 1:**
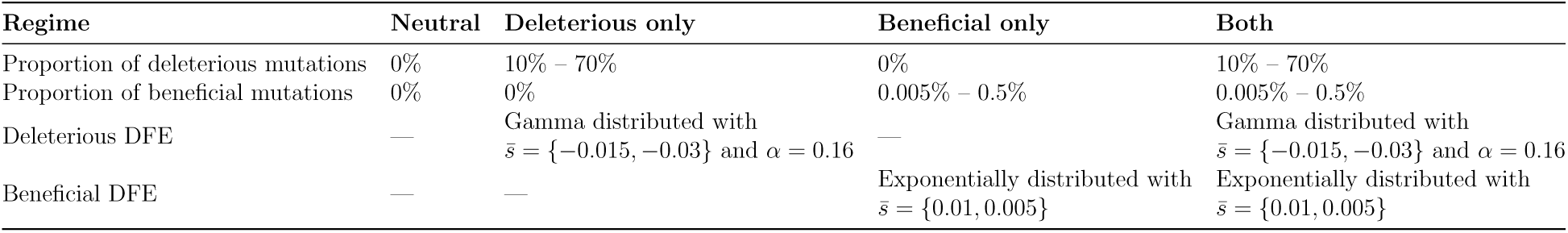
Range of parameters explored in the simulations. Non-neutral mutations were only allowed within exons. “DFE” refers to the distribution of fitness effects. Gamma distribution was parameterized with shape *α* and mean *s̄*=*α/β*, where *β* is the rate parameter.

### 2.3 Visualizing correlated landscapes of diversity and divergence

To compare landscapes of diversity and divergence along chromosomes, we computed the Spearman correlation between the landscapes across windows within a chromosome. Because of computational constraints, we focus on chromosome 12. Chromosome 12 is one of the smallest chromosomes in the great apes, there are no major inversions, and it has good variation in exon density and recombination rate. The choice was made blindly before looking at the data, but we found it behaves similarly to other chromosomes (see Figure S10 through Figure S31).

We expected landscapes of two closely related species to be more correlated than the landscapes of two distantly related species. Thus, the correlation between any two landscapes of diversity and divergence is expected to depend on distances between them in the phylogenetic tree. We decided to plot our correlations against distance (in generations) between the most common recent ancestor (MRCA) of each landscape. These distances were computed using the demographic model estimated in Prado-Martinez et al. (2013). In comparing two landscapes of diversity, this amounts to the total distance between the two tips in the species tree. For instance, the phylogenetic distance *dT* between diversity in humans and diversity in bonobos is the sum of the lengths of the human, pan and bonobo branches in the species tree used for simulation (shown in Figure 1). In comparing a landscape of diversity to a landscape of divergence, this amounts to the distance between the species of the landscape of diversity and the MRCA of the two species involved in the divergence. For example, *dT* for the landscapes of diversity in humans and divergence between Sumatran orangutans and eastern gorillas would be the distance between the humans tip and the great apes internal node. *dT* for the landscapes of divergence between the orangutans and divergence between the gorillas would be the distance between the orangutan and gorilla internal nodes. Some divergences may share branches in the tree, but these are excluded from our main figures; see subSection 4.1 and Figure S2.

## 3 Results

First, we will provide a qualitative view of the landscapes of diversity and divergence in the great apes. Then, we explore the correlations between landscapes in the real data and how they vary depending on phylogenetic distance. To understand the processes that can drive these correlations, we use forward-in-time simulations of the great apes history under different models (e.g., with and without natural selection). Lastly, we describe how genomic features are related to patterns of diversity and divergence in the real great apes data, and we speculate which processes can explain what we see in the data and simulations.

### 3.1 Landscapes of within-species diversity and between-species divergence

There is considerable variation in levels of genetic diversity across the great apes (Figure 2). Species may differ in overall levels of diversity due to population size history: species with greater historical population sizes (e.g., central chimps and western gorillas) harbor the most amount of genetic variation (Prado-Martinez et al., 2013). Levels of diversity vary along the chromosome, but do not appear to be strongly structured. Instead, diversity seems to haphazardly fluctuate up and down along the chromosome, and this variation might be attributed to neutral genealogical and mutational processes alone. A notable feature is the large dip in diversity around the 50Mb mark, which is so extensive that it almost erases the differences between-species. This dip coincides with three of the windows with the highest exon density, possibly pointing to the role of selection in shaping genetic variation in those windows.

**Figure 2:**
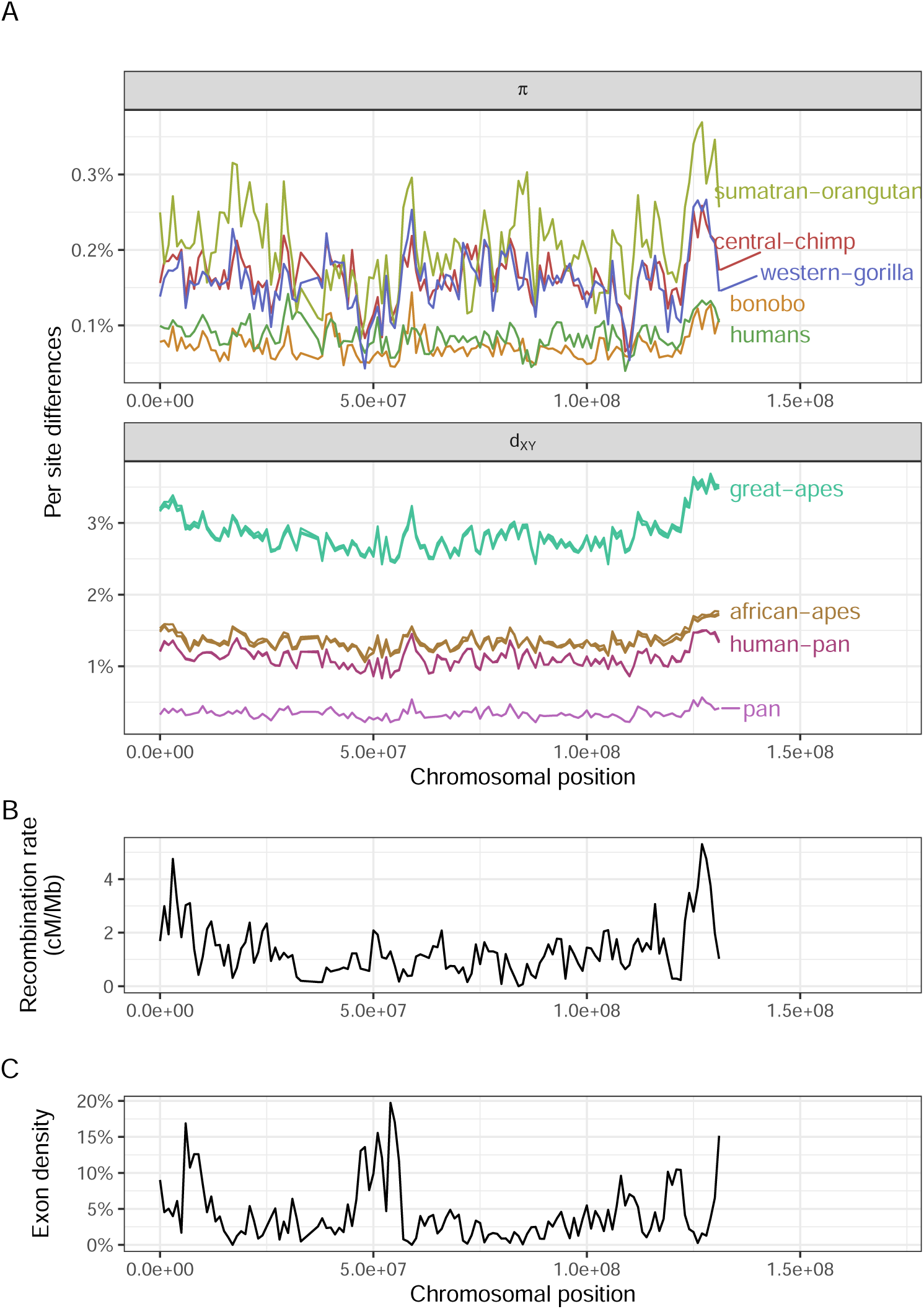
A) Landscapes of nucleotide diversity (*π*) and divergence (*d_XY_*) in 1Mb windows along chromosome 12. Lines are colored by species on the top plot and by the most common recent ancestor (MRCA) on the bottom. Genomic windows with less than 40% of callable sites were masked. Only a subset of the species are displayed for clarity. B) Recombination rate estimates from humans (deCODE map; Kong et al., 2010). C) Exon density along chromosome 12, computed as the percentage of callable nucleotides in a window that fall within an exon. for samples within species happen before the species split). As a result, samples from one population may coalesce first with a sample from another population (e.g., samples *v*_2_ and *w*_1_), a pattern called incomplete lineage sorting (ILS) (see the branch marked with *\** in the gene tree). This sharing of ancestral variation causes *π_V_* and *π_W_* to be correlated with each *<u>−T</u>* other. The probability two samples from *V* coalesce before the split with *W* is 1*−e* ^2^*^N^e*, where *T* is the split time and *N_e_* is the effective population size. Therefore, split time (*T*) should be a good predictor of the correlation between two landscapes of diversity (and/or divergence). Thus, we decided to visualize correlations between landscapes of diversity and divergence by computing the phylogenetic distance *dT*, which is simply the distance in generation time between two statistics. For example, we define *dT* (*π_W_, d_XY_*) = 2*T_V W XY_ − T_XY_*. Divergences may share branches by definition (irrespective of split times), as you can see with *d_V X_* and *d_XY_* (see subsection 2.3 for more details). In such cases, our chosen metric *dT* would not be a good proxy for expected correlations, so we omit such cases from our main figures. See subsection 2.3 and Figure S2 for more on the correlations between landscapes that share branches.

Levels of between-species genetic divergence also vary along the genome, by an even greater amount in absolute terms. Interestingly, diversity (*π*) varies (along the chromosome) by about 0.2%, whereas divergence (*d_XY_*) varies by more than 0.5%. Because *d_XY_* = *π*^anc^+*rT* (where *π*^anc^ is diversity in the ancestor, *r* is the substitution rate and *T* is the split time between the two species), this excess in variance may be due to the substitution process. Landscapes of divergence which share their most common recent ancestor (e.g., human-Bornean orangutan and bonobo-Bornean orangutan divergences — both colored in red in Figure 2A) overlap almost perfectly with each other. Curiously, divergence seems to accumulate faster in the ends of the chromosome, leading to a “smiley” pattern in the landscape of divergence — which is not apparent in the landscape of diversity. That is, with deeper split times, divergence in the ends of the chromosome seem to increase faster than in other regions of the genome (see how the divergences whose MRCA is the great apes look more like a convex parabola than a horizontal line in Figure 2A; see also Figure S1).

In comparing landscapes across species side by side, a remarkable pattern emerges: levels of genetic diversity and divergence along chromosomes have similar peaks and troughs. To get a sense of how strong this observation is, we can compare it to one of the most well studied properties of genomic variation: the correlation between exon density and genetic diversity. We found that the correlation between human diversity and exon density is 0.2 (at the 1Mb scale), but the correlation between levels of diversity in humans and western gorillas is 0.48. Below, we dissect this observation of strong correlation between landscapes across the great apes and discuss the processes that may cause it.

### 3.2 Remarkable correlations between landscapes of diversity and divergence

The landscapes of diversity and divergence are highly correlated across the great apes. To interpret this signal, we first need to understand what processes can cause such correlations, and so first we describe the toy example depicted in Figure 3. Both genetic diversity (*π*) and divergence (*d_XY_*) are estimates of the mean time to the most recent common ancestor (multiplied by twice the effective mutation rate). Populations *V* and *W* split recently, and so ancestral variation contributes significantly to within-species diversity (i.e., the coalescences

**Figure 3:**
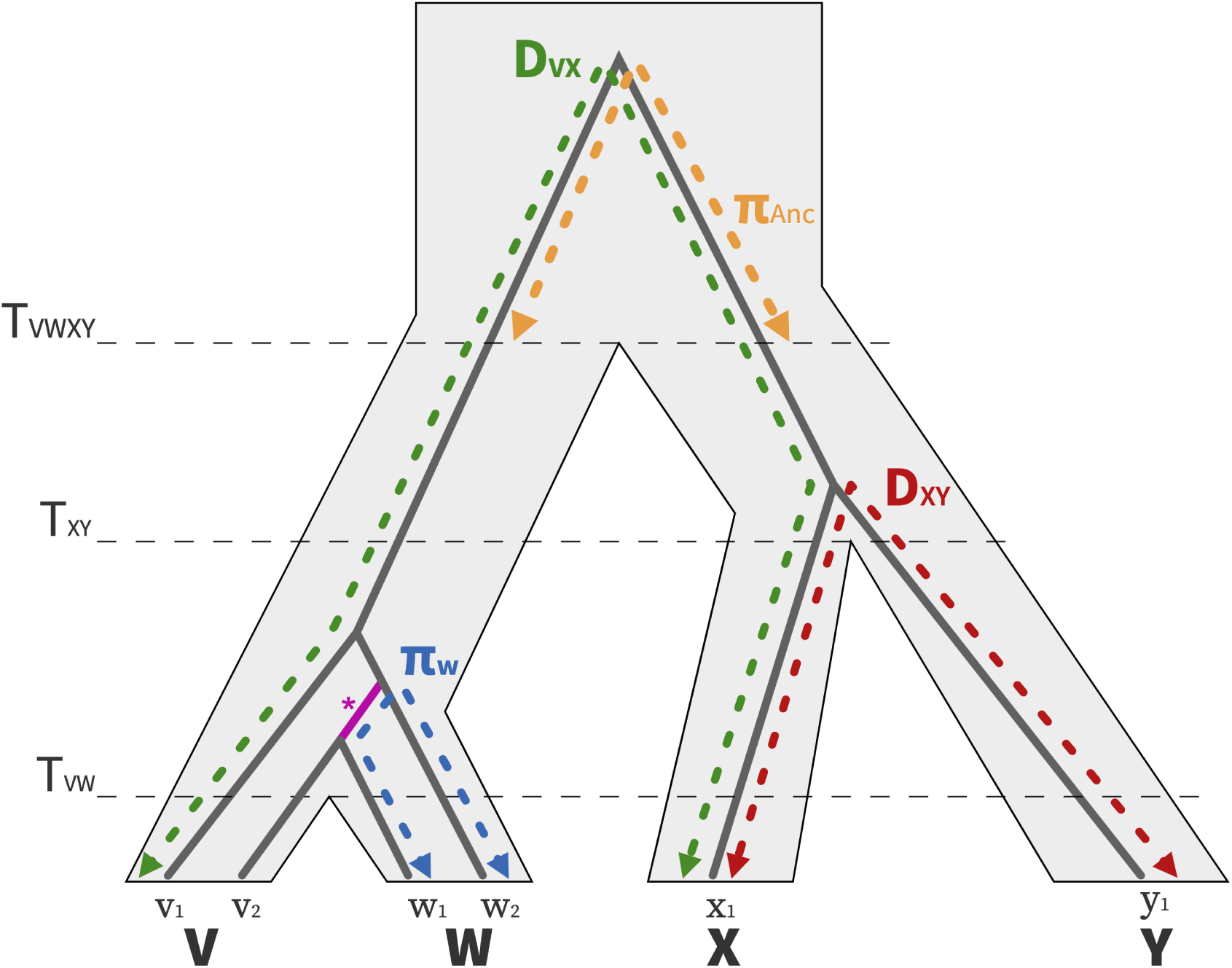
Visualizing the relationships between nucleotide diversity and divergence statistics between closely related taxa. A population and gene tree for four populations (V, W, X, Y) are depicted with the light gray polygon and gray solid line, respectively. The branch that is shared between *π_V_* and *π_W_* due to incomplete lineage sorting is highlighted in pink.

Figure 4 shows the pairwise correlations between great apes landscapes of diversity and divergence against phylogenetic distance (*dT*, which is computed from the split times of the model shown in Figure 1). We see ancestral variation seems to play a role in structuring correlations between landscapes: pairs of species that recently split have their landscapes of diversity highly correlated. Surprisingly, correlations still plateau at around 0.5. We expect ancestral variation to play a minor role when comparing orangutans and chimps, which separated around 60 *N_e_* generations ago, but their landscapes are still highly correlated. Population size history seems to affect the correlation between landscapes since the weakest correlations involve the landscape of diversity of one of the species with small historical population sizes (i.e., bonobos, eastern gorillas and western chimps).

**Figure 4:**
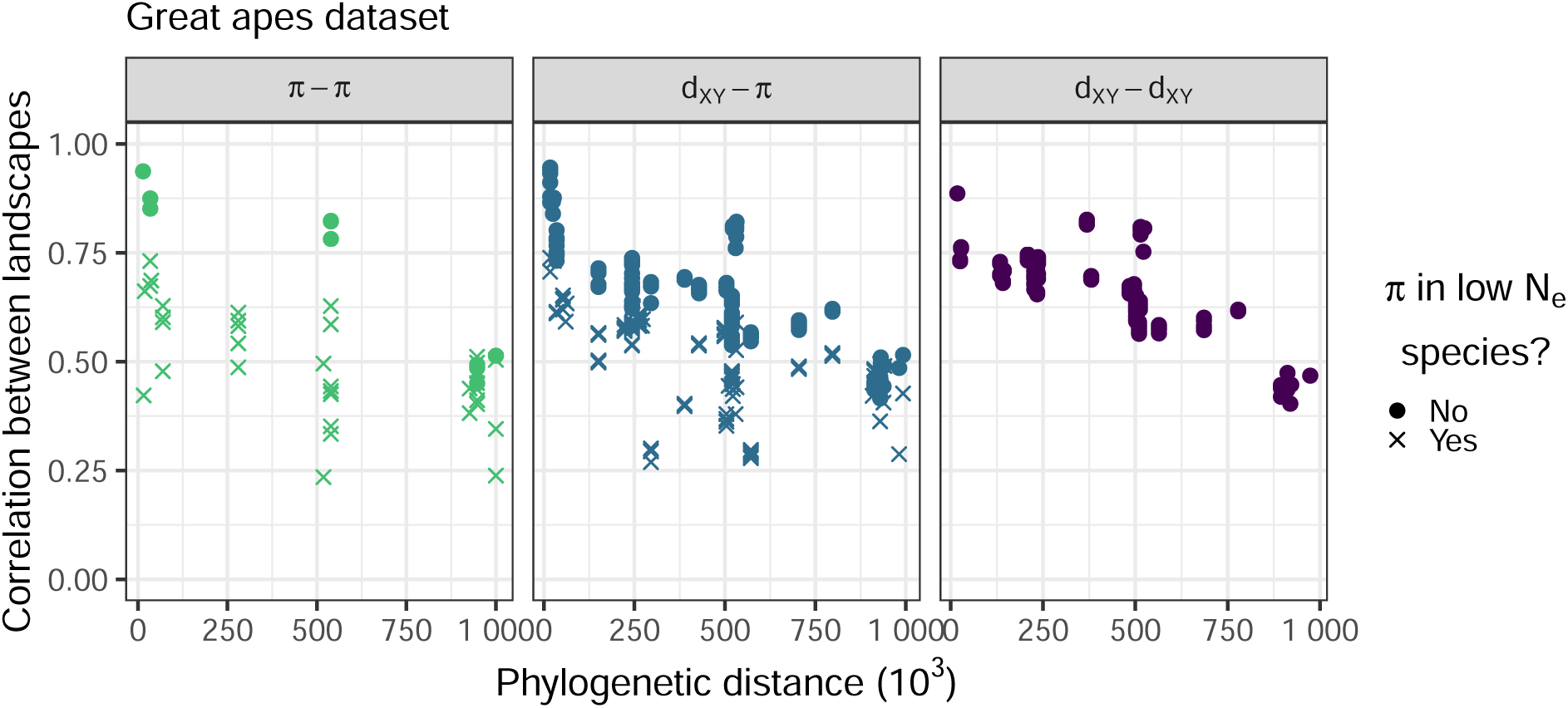
Correlations between landscapes of diversity and divergence across the great apes. Each point on the plots correspond to the (Spearman) correlation between two landscapes of diversity/divergence, computed on 1Mb windows across the entire chromosome 12. Correlations were split by type of landscapes compared (*π π*, *π d_XY_*, *d_XY_ d_XY_*). *dT* is the phylogenetic distance (in number of generations) between the most common recent ancestor of the two landscapes compared (e.g., the *dT* for correlation between landscapes of diversity in humans and divergence between eastern gorillas and orangutans is distance between the humans and the great apes nodes in the phylogenetic tree, Figure 1). Note that species with low *N_e_* — for which the estimated species *N_e_* was less than 8 10^3^: bonobos, eastern gorillas and western chimps — have a different point shape. Only comparisons for which the definition of the statistics do not overlap are shown, as explained in subsection 2.3.

Correlations between landscapes of divergence and diversity and between landscapes of divergence are also quite high, often surpassing 0.5, and they also decay with phylogenetic distance (*dT*) (see middle and right most plots in Figure 4). In theory, these landscapes can also be correlated due to ancestral variation. To see how ancestral variation can create correlations even between landscapes with no overlap in the tree, consider Figure 3: divergence between X and Y and divergence between V and W can each contain contributions from ancestral diversity if lineages have not coalesced in both branches leading from the ancestor. If a particular portion of the genome happens to have higher diversity in the ancestor, it will also have higher divergence. Since this correlation is produced by sharing of ancestral variation, it is expected to have a very small effect except when branches are short. As discussed in subsection 2.3, two divergences can also be correlated by definition (because they share branches in the tree). For example, when comparing human-Bornean orangutan and gorilla-Bornean orangutan divergence we expect some correlation because these divergences share the large African apes and orangutan branches in the tree (Figure 1). In Figure 4 we excluded these comparisons where branches are shared. Such comparisons can be seen in Figure S2. We found that even these comparisons that share branches have an excess of correlation compared to a theoretical expectation (derived from a simplified neutral model), that is the correlations are above the *y* = *x* line in Figure S2 even for distantly related species.

There are many processes that could maintain landscapes correlated. Above, we discussed how we expect ancestral variation to explain these correlations. The alternative would be to have a process that structures variation along chromosomes which is shared across species. Using forward-in-time simulations, we set out to (i) confirm that ancestral variation alone is not causing landscapes to remain correlated, and (ii) test which process or processes that when shared among a group of species could maintain correlations in similar ways to what we observed in the great apes’ data.

### 3.3 Neutral demographic processes

To assess the extent to which ancestral variation alone could explain our observations, we performed a forward-in-time simulation of the great apes’ evolutionary history. As expected, the resulting landscapes of diversity and divergence are not well correlated (Figure 5). Ancestral variation seems to maintain correlations between some landscapes; for instance, the landscapes of diversity in central and eastern chimps have a 0.61 correlation, the highest across all pairs of comparisons (Figure 5A, point *a*). Nevertheless, correlations between landscapes of diversity and divergence decay quickly with phylogenetic distance to 0. Some distant comparisons are moderately correlated (e.g., the landscape of diversity in Bornean orangutans and divergence between central and western chimps have a correlation coefficient of 0.23, see Figure 5A, point *b*), but that seems to be driven by the outlier window around 80Mb. This outlier window has a recombination rate close to 0 (Figure 2C), so the average nucleotide diversity over the window has a higher variance because of coalescent noise (see the extreme peaks and valleys in Figure 5). Recombination rate variation can create some moderate correlations, but when we look at multiple species at once it becomes clear that the mean correlation goes to 0.

**Figure 5:**
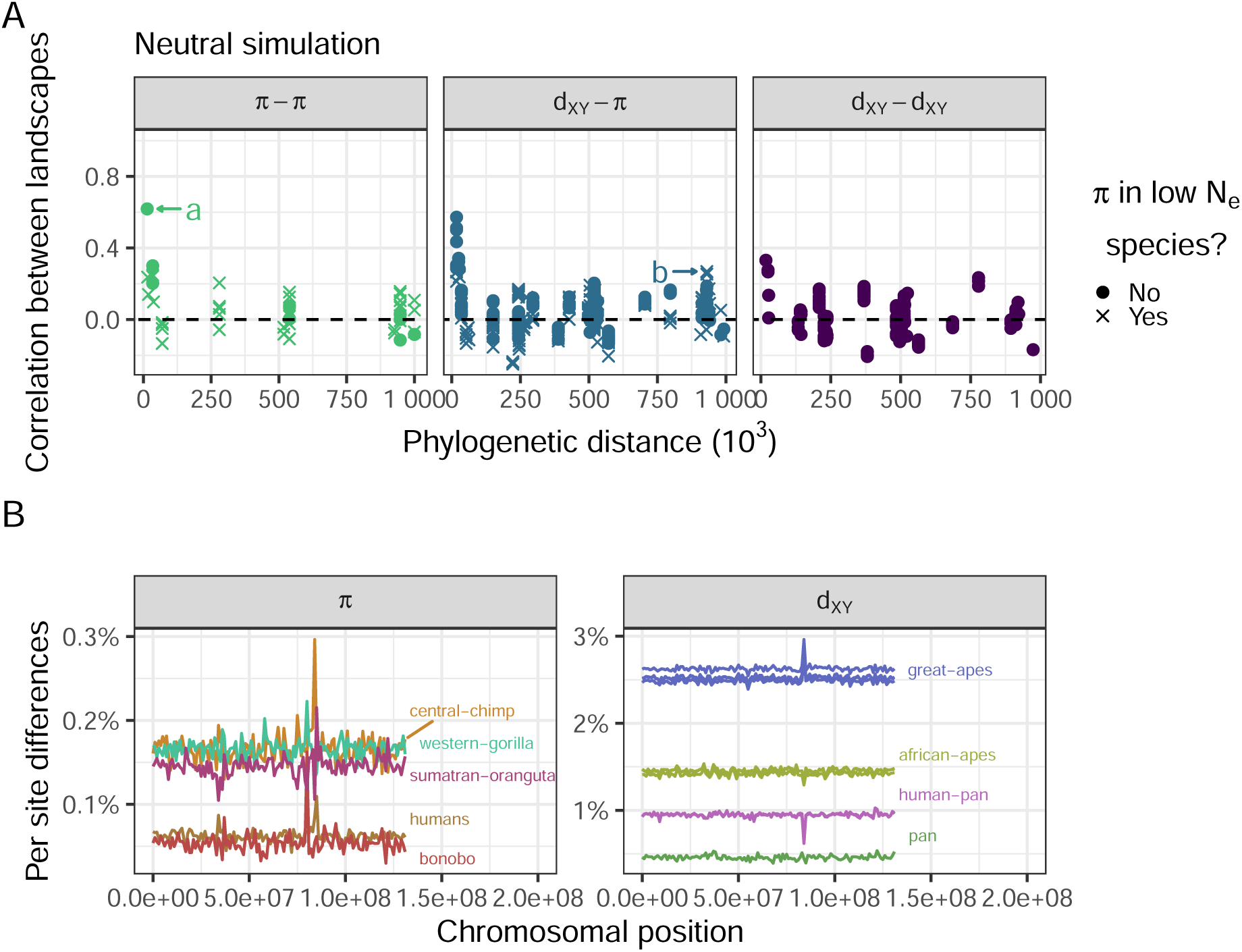
Landscapes are not well correlated in a neutral simulation. (A) Correlations between landscapes of diversity and divergence in a neutral simulation. See Figure 4 for more details. (B) Nucleotide diversity and divergence along the simulated neutral chromosome. See Figure 2A for details.

**Figure 6:**
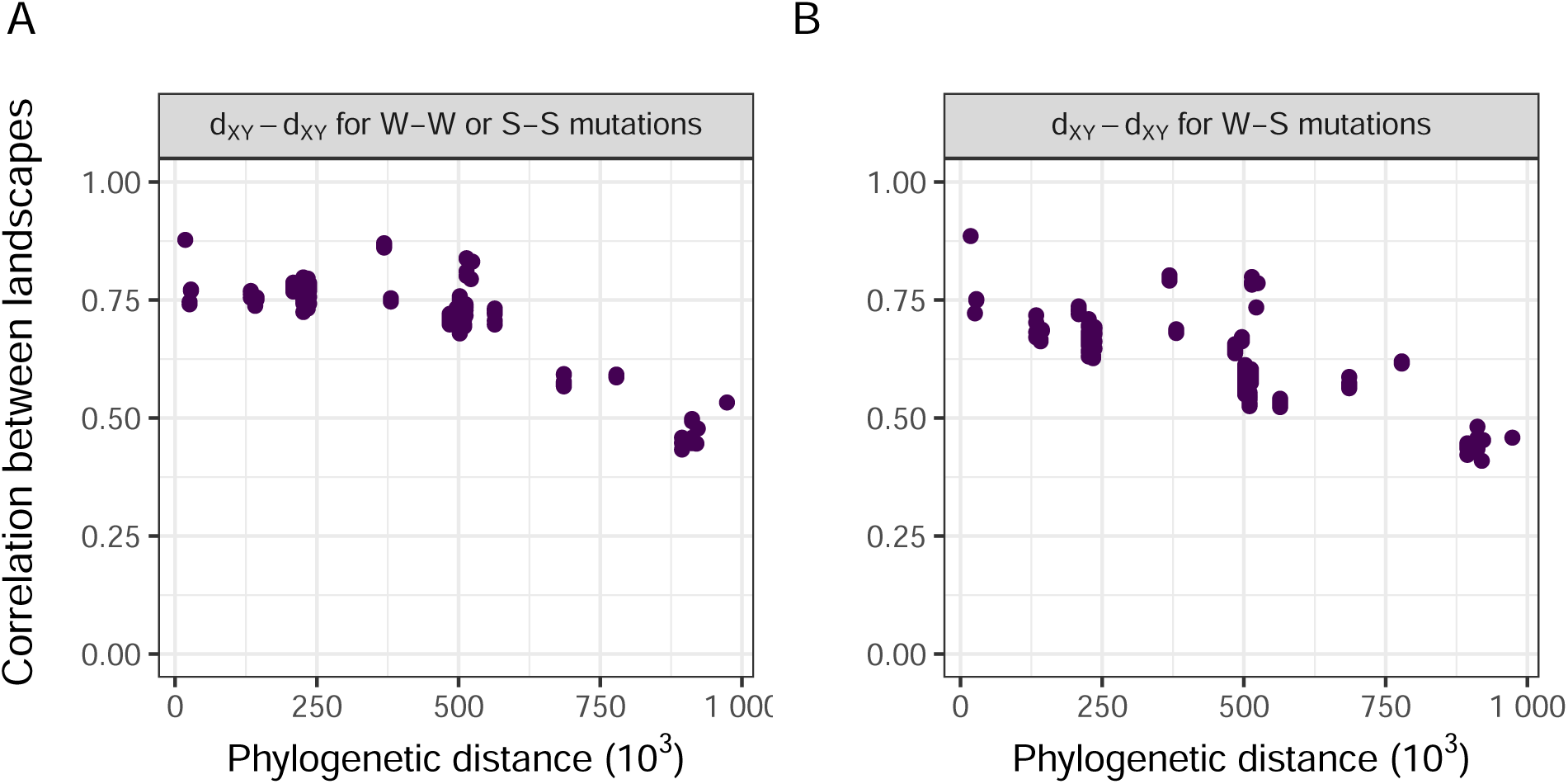
Correlations between landscapes of divergence partitioned by site type (W-W/SS and W-S). W-W sites are sites in which the state did not change between-species (and remained weak which corresponds to A or T). Similar logic applies to S-S sites (S or strong states are G or C). W-S sites are sites in which a new mutation appeared either going from weak to strong or from strong to weak. Note these definitions do not rely on identifying the exact ancestral state, we simply compare the current states in the four species involved (two species per *d_XY_* landscape). For example, if by looking at the four species we see the following states A,T,A,T the site would be classified as W-W. If we saw G,A,A,A the site would be classified as W-S. Other details are the same as in the rightmost panel in Figure 4.

### 3.4 GC-biased gene conversion

A prominent feature of the landscapes of divergence in the great apes is the faster accumulation of divergence in the ends of the chromosomes (Figure 2). This feature was not present in any of our simulations, so we sought to understand its possible causes. Double strand breaks are more common at the ends of chromosomes (Kong et al., 2002), and these can be repaired either by crossover or gene conversion events. GC-biased gene conversion (gBGC), the process whereby weak alleles (A and T) are replaced by strong alleles (G and C) in the repair of double-stranded breaks in heterozygotes, mimics positive selection – in that it increases the probability of fixation of G and C alleles (e.g., Galtier et al., 2009). We suspected gBGC could have caused the increased rate of accumulation divergence in the ends of chromosomes, as has been observed previously (Katzman et al., 2010), and contributes to the maintenance of correlations between landscapes over long time scales.

To tease apart the effects of gBGC on correlated landscapes, we partitioned divergence by mutation type (weak to weak, strong to strong and weak to strong). If correlations are being driven by gBGC, then we would expect the correlation between landscapes of divergence to be stronger for weak to strong mutations. We found that the overall correlations are very similar across mutation types, suggesting gBGC does not play a strong role in structuring the correlations between landscapes (Figure 6).

### 3.5 Positive and negative natural selection

Another process whose intensity is likely correlated across all branches in the great apes tree is natural selection. If targets of selection and recombination maps are shared across species, then we would expect both the direct and indirect effects of selection to be shared across branches. It can be difficult to model natural selection in a realistic manner because we do not know precisely which locations of the genome are subject to stronger selection. Nevertheless, exons are expected to have higher density of functional mutations than other places in the genome. Thus, we ran simulations in which beneficial and deleterious mutations can happen only within exons. Using human annotations, we simulated the great apes’ history assuming a common recombination map and exon locations. See the landscapes from the simulations in Figure 7.

**Figure 7:**
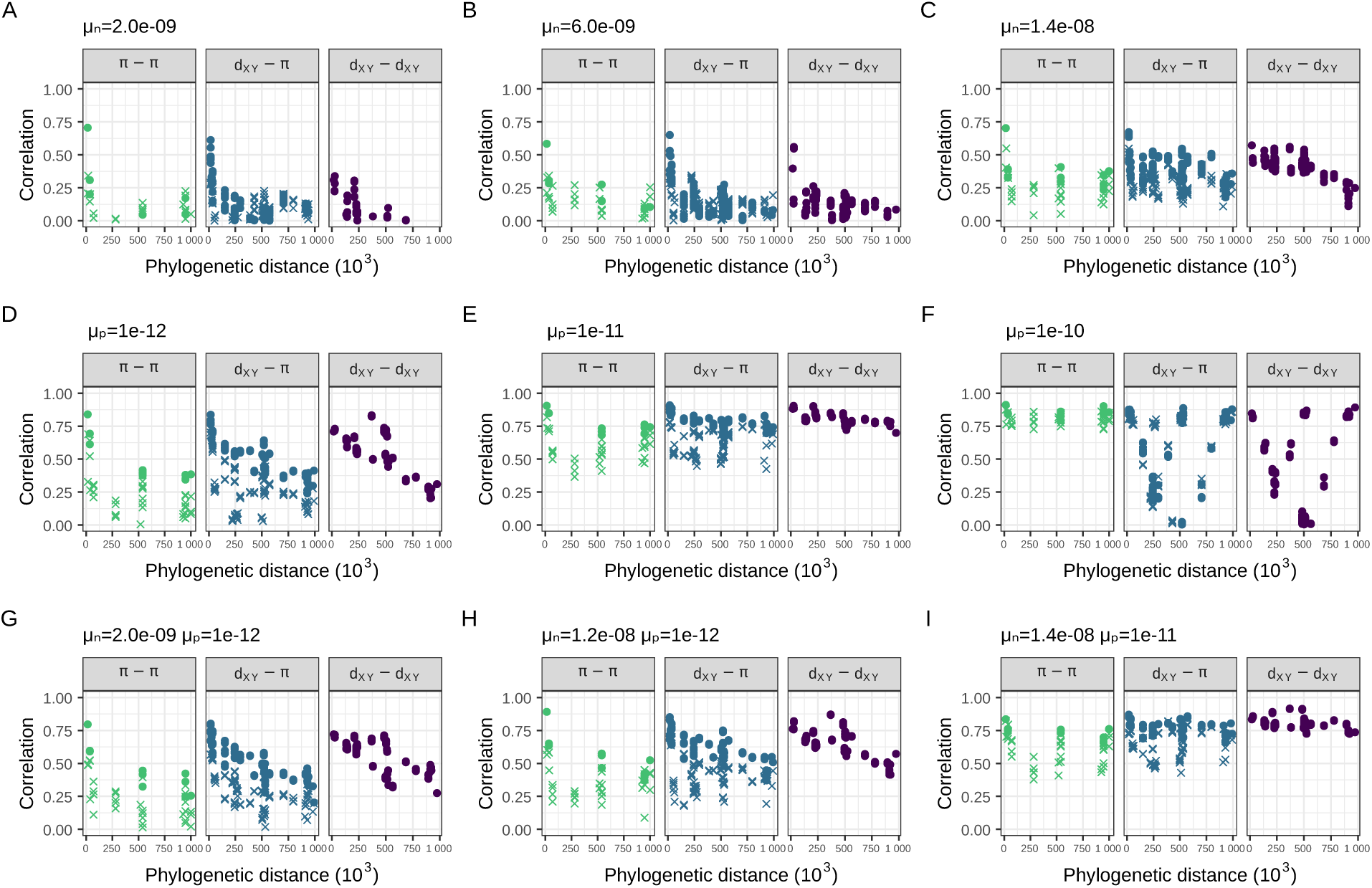
Correlations between landscapes of diversity and divergence in simulations with natural selection. (A-C) Simulations with negative selection. (D-F) Simulations with positive selection. (G-I) Simulations with both negative and positive selection. The selection parameters *µ_n_* and *µ_p_* are the rate of mutations in exons with negative and positive fitness effects, respectively. The mean fitness effect was *s̄* = 0.03for deleterious mutations and *s̄* = 0.01 for beneficial mutations (see subsection 2.2 for more details). See how panel H looks the most like the data (Figure 4). *µ*_SD_ = 7% could plausibly create the correlations observed in the real data (Figure 8D-I). The neutral and deleterious simulations with mutation rate variation fail to recover one aspect of the real data: the lower correlations between landscapes that include at least one low *N_e_* species (seen in the *π π* and *π d_XY_* comparisons of Figure 4). This feature, however, is seen in the simulations with both beneficial and deleterious mutations (Figure 8G-I).

We found that negative selection can slightly increase correlations between landscapes (Figure 7A-C). If 30% of all mutations within exons were strongly deleterious (mean selection coefficient *s̄* = *−*0.03), landscapes would be weakly correlated (Figure 7B). The correlations between landscapes rarely surpass 0.5, even with 70% of all mutations within exons being strongly deleterious (Figure 7C).

Positive selection, on the other hand, can quickly increase correlations between land-*_−_*_12_ scapes. A beneficial mutation rate within exons of *µ̄_p_* = 1 *×* 10^-12^ produced moderate correlations between landscapes (Figure 7D). With too much positive selection, correlations can break down because of the contrasting effects of positive selection on diversity and divergence. That is, while positive selection increases fixation rates and hence divergence between-species, its linked effects decrease diversity within the species. This can create negative correlations between landscapes, as can be seen in Figure 7F. Note that some correlations between landscapes of diversity and divergence remain high when the divergence is computed between closely related species (e.g., central and eastern chimps). Divergence is *d_XY_* = *π*^anc^ + 2*rT*, where *π*_anc_ is diversity in the ancestor, *r* is the substitution rate and *T* is the time since species split. Thus, for the divergences in which the two species split recently are dominated by genetic diversity in the ancestor, correlations between *π d_XY_*remain high because *d_XY_ π*^anc^.

Positive and negative selection can work synergistically to produce correlated landscapes that look like the real data. For example, comparing figures Figure 7D,G,H which differ in rate of negatively selected mutations *µ_n_*, it is possible to see that the correlations between landscapes start to resemble the real data with more deleterious mutations. Figure 7H seems to resemble the data fairly well, with *π d_XY_* and *d_XY_ d_XY_* correlations plateauing around 0.5. The *π − π* correlations are a bit lower than the real data, however. Recent demographic events can affect genetic diversity and although our simulations are heavily parameterized with respect to the effects of selection, we are not capturing all the variation caused by more realistic demographic models. Figure 7D and H look very similar to each other. These have the same amount of positive selection, but the first did not have any negative selection. The major difference between them is that with negative selection there is a more clear separation between the correlations involving low *N_e_*species, similar to what is seen in the data.

### 3.6 Mutation rate variation

Since mutation rate can vary along chromosomes, if this mutation rate map were shared across species, it would maintain correlations between landscapes over longer periods of time. To assess this, we used three of our previous simulated genealogies of the great apes and replaced all neutral mutations assuming a common neutral mutation rate map across the phylogeny: for each window, we drew a mutation rate from a normal distribution with mean 2 10*^−^*^8^ (the same as all other simulations) and standard deviation *µ*_SD_. We found that, under neutrality, a mutation rate map with *µ*_SD_ close to 7% 2 10*^−^*^8^ would be needed to get correlations similar to the data (Figure 8A-C). Although mean correlations look similar to the data, we see that correlations tend to increase slightly with time in the simulations with mutation rate variation. This is expected because windows with higher mutation rate accumulate divergence faster, creating a correlation with mutation rate that gets stronger with time. In the great apes’ data, however, we see a slow but steady decrease in correlations with time.

**Figure 8:**
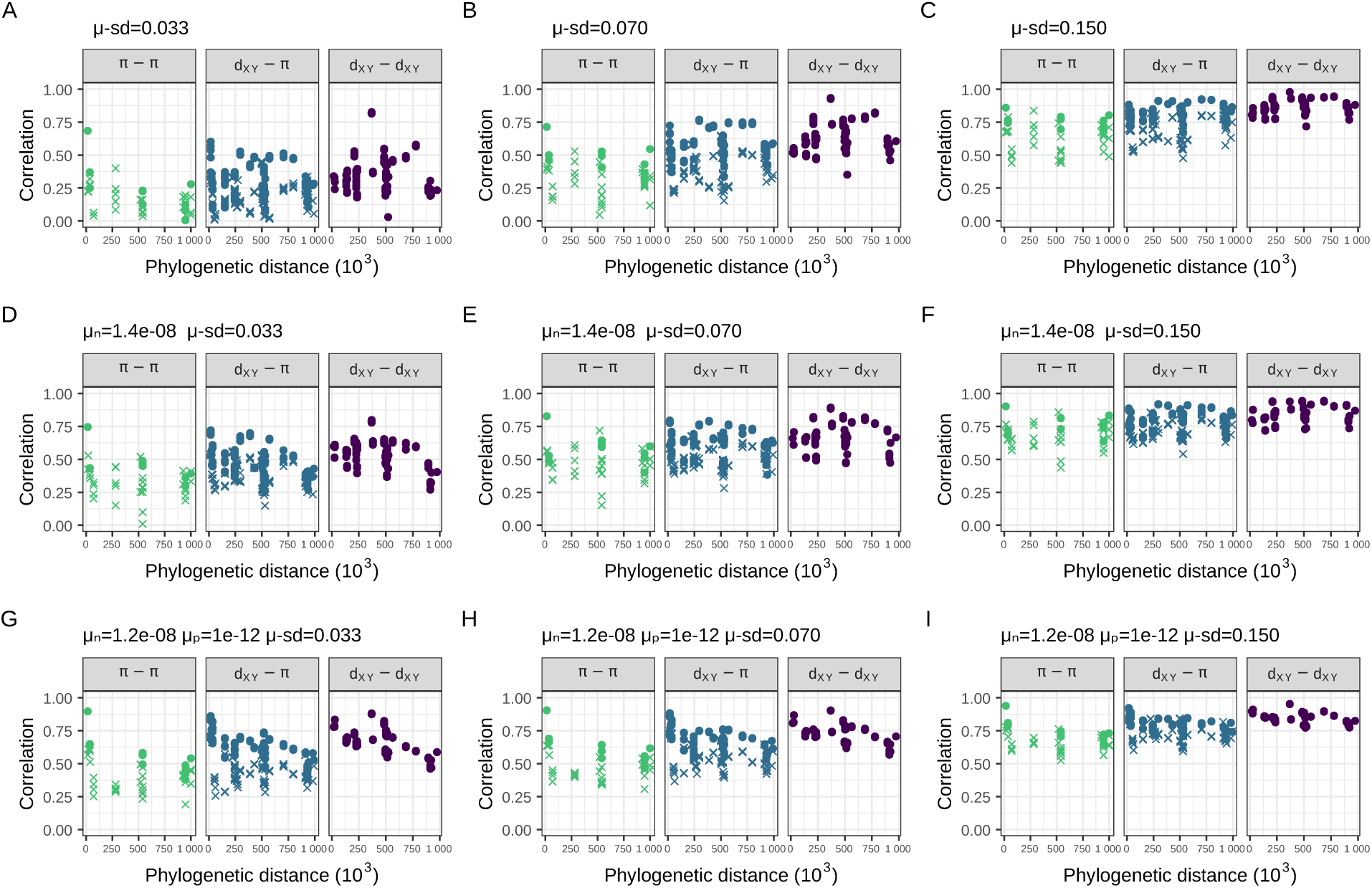
Correlations between landscapes of diversity and divergence across the great apes for simulations with variation in mutation rate along the chromosome. Panels A through I show different simulations in which we varied the standard deviation in neutral mutation rate between 1Mb windows, in each setting the standard deviation to the mean mutation rate (2 10*^−^*^8^) multiplied by *µ*_SD_. First row (A-C) use a neutral simulation, second row (D-F) a simulation with negative selection, and third row (G-I) a simulation with both positive and negative selection. The selection parameters *µ_n_* and *µ_p_* are the rate of mutations in exons with negative and positive fitness effects, respectively. The mean fitness effect was *s̄* = *−*0.03 for deleterious mutations and *s̄* = 0.01 for beneficial mutations (see subsection 2.2 for more details). See how without selection (A-C), the simulation with *µ*_SD_ = 7% (panel B) looks close to the data (Figure 4). With selection, the simulation with both positive and negative selection and *µ*_SD_ = 3.3% looks even more similar to the data (correlations between divergences decay over time, and there is a more pronounced differentiation between low and high *N_e_* comparisons).

When we added variation in the neutral mutation rate to simulations with selection, we found that a mutation rate map with a standard deviation of rates of slightly less than

### 3.7 Visualizing similarity between simulations and data

To see how a particular simulation resembles the real data, we can use figures Figure 4 and Figure 7 to compare how the patterns of all 1260 pairwise correlations between landscapes match the real data. However, it is difficult to assess the fit of the simulated scenarios to real data from such a comparison. Instead, we use principal component analysis (PCA) and create a low dimensional visualization, shown in Figure 9, in which each point is a simulation or the real data (shown in yellow). We created this PCA from the 57 1260 matrix in which rows are the simulations and the data, and columns are the pairwise Spearman correlations between landscapes. Unlike in the plots above, here we include the correlations between overlapping landscapes (as detailed in subsection 2.3) (Figure 9). In PC space, the data most closely resembles a subset of our simulations with both positive and negative selection (e.g., *µ̄_p_* = 1 10*^−^*^12^ and *µ̄_n_* = 1.2 10*^−^*^8^), including no or very little variation in mutation rates (less than 4%).

**Figure 9:**
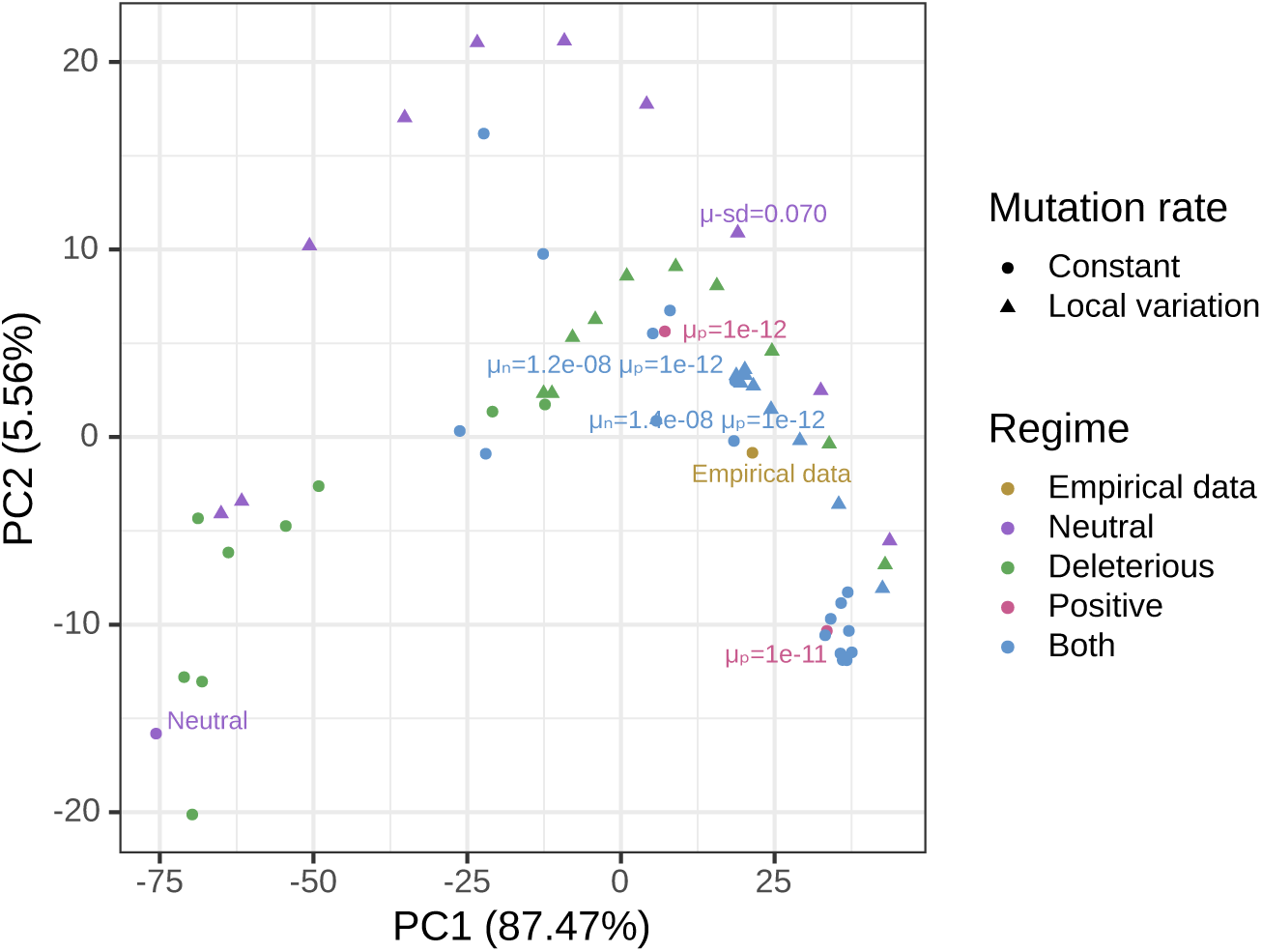
Principal component analysis (PCA) visualization of data and simulations. Colors differentiate the empirical data from simulations with different parameters: “Neutral” refers to the simulation without any selection, “Deleterious” refers to simulations with deleterious mutations, “Positive” refers to simulations with beneficial mutations, “Both” refers to simulations with both beneficial and deleterious mutations. The shape of the points differentiate simulations with constant mutation rate along the genome and variable local mutation rates. Local variation in mutation rates were added on top of three simulations: neutral, deleterious with *µ_n_* = 1.4 10*^−^*^8^, and both with *µ_n_* = 1.2 10*^−^*^8^ and *µ_p_* = 1 10*^−^*^12^. The PCA performed on a matrix containing all pairwise correlations between landscapes across the great apes (i.e., all *π π*, *π d_XY_* and *d_XY_ d_XY_* comparisons) for the great apes dataset and simulations (with selection and with mutation rate variation). We excluded simulations with *µ_p_* 1 10*^−^*^10^ from the PCA analysis because PC2 was capturing negative correlations caused by strong positive selection — as seen in Figure 7F.

We also performed PCA on correlations computed at two different scales, 500Kb and 5Mb, in addition to the previously shown results for 1Mb (Figure S5, Figure S6). At 500Kb, the observed data are slightly more distant from simulations than at the higher scales, possibly because the recombination map used in simulations had a coarser resolution. Nevertheless, the observed data most closely match the simulations with both positive and negative selection in all scales.

### 3.8 Correlations between genomic features and diversity and divergence

Next, we describe how two important genomic features (i.e., exon density and recombination rate) are related to diversity and divergence in the real great apes data set. The correlations between recombination rate and genetic diversity are positive in all great apes (Figure 10A). The strongest correlation between genetic diversity and recombination rate is seen in humans, which is unsurprising given our recombination map was estimated for humans. Recent demographic events also seem to impact the strength of the correlation; for example, the correlation between recombination rate and diversity is higher in Nigerian chimps than in western chimps, which have a much lower recent effective population size. We found that diversity is negatively correlated with exon density across all species (Figure 10D). Contrary to what we observed with recombination rate, the correlation between exon density and diversity was even stronger in most other apes than in humans. Species with smaller *N_e_* tend to show weaker correlation between diversity and exon density (see Nam et al., 2017 for related findings). A striking feature of the correlations of between-species divergence and genomic features, shown in Figure 10, is that the correlations get stronger with the amount of phylogenetic time that goes into the comparison (i.e., the *T_MRCA_*), in a way that is roughly linear with time.

**Figure 10:**
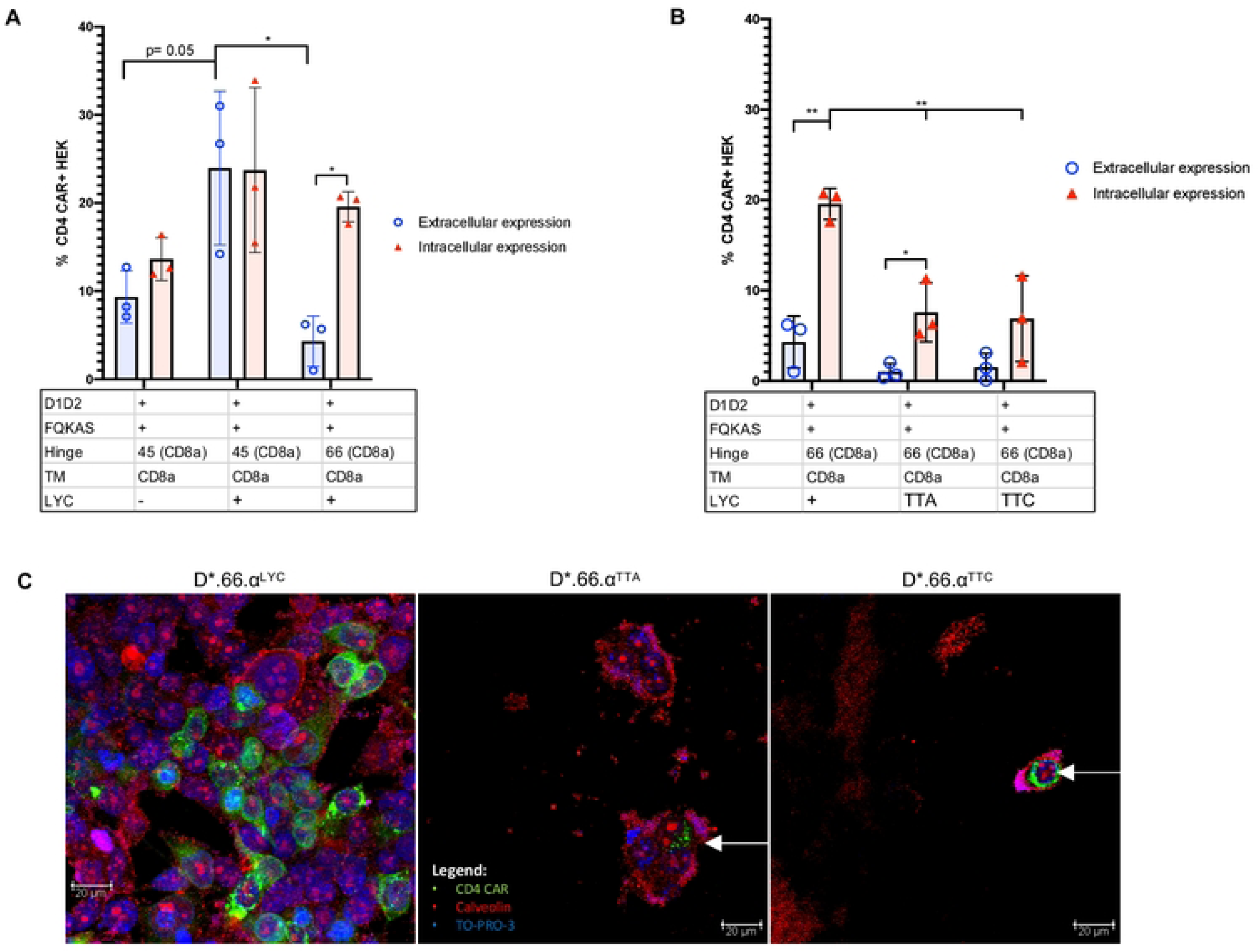
Correlations and covariances between landscapes of diversity and divergence and annotation features in the real great apes data. Exon density and recombination rates were obtained as detailed in Figure 2. Split time is the time distance between the two species involved in the divergence. Points are colored by the species of within-species diversity (*π*) in plots A and D. In plots B,C,E,F, the points are colored by the most common recent ancestor of the species for which between-species divergence was computed. Species with low *N_e_* — for which the estimated species *N_e_* was less than 8 10^3^: bonobos, eastern gorillas and western chimps — have a different point shape.

To describe why this increase in correlation with time might occur, we turn to an analytic approach. Genetic divergence (*D*) in the *i*^th^ window between two species that split *t* generations ago can be decomposed as:

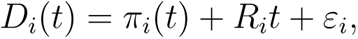

where *π_i_*(*t*) is the genetic diversity in the ancestor at time *t*, *R_i_*is the substitution rate in the window and *ε_i_* is a contribution from genealogical and mutational noise (which has mean zero). This decomposition follows from the definition of genetic divergence as the number of mutations since the common ancestor, as depicted in Figure 3 (see how *D_V X_* = *π*^anc^ + 2*RT_V WXY_*).

The covariance between *D*(*t*), the vector of divergences along windows, and a genomic feature *X* is, using bilinearity of covariance,

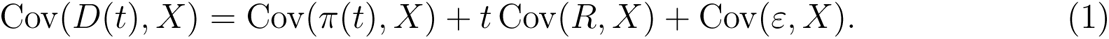

Happily, this equation predicts the linear change of the covariance with time that is seen in Figure 10C and perhaps Figure 10F. However, caution is needed because the correlation between diversity and the genomic feature (Cov(*π*(*t*), *X*)) may be different in different ancestors, and indeed the inferred effective population size is greater in older ancestors in the great apes (Figure 1).

Next consider covariances of diversity with recombination rate, Figure 10C. Consulting the equation above, the fact that the covariance between divergence and recombination rate increases with time can be caused by two factors (taking *X* to be the vector of mean recombination rates along the genome): (i) a positive covariance between substitution rates and recombination rates (Cov(*R, X*) *>* 0), and/or (ii) greater genetic diversity in longer ago ancestors (*N_e_*(*t*) larger for larger *t*). It is unlikely that the increase in *N_e_* in more ancient ancestors was sufficient to produce the dramatic increase in covariance seen in Figure 10C, since it would require Cov(*π*(*t*), *X*) to be far larger in the ancestral species than is seen in any modern species. On the other hand, there are various plausible mechanisms that would affect Cov(*R, X*). One factor that certainly contributes is the “smile”: we found that divergence increases faster near the ends of the chromosomes where recombination rate is greater, probably in part because of GC-biased gene conversion. Interestingly, positive and negative selection are predicted to have opposite effects here: greater recombination rate increases the efficacy of both through reduced interference among selected alleles, so positive selection would increase substitution rate and hence increase Cov(*R, X*), while negative selection would decrease Cov(*R, X*). When considering only the middle half of the chromosome (i.e., excluding the effect of gBGC) (Figure S7), the covariances between divergence and recombination rate flip to negative, and they continue to decrease over time. Thus, it seems that negative selection is the most important driver of divergence in the middle, whereas gBGC strongly affects the tails of the chromosome.

The covariance of diversity and exon density has a less clear pattern (Figure 10F), although it generally gets more strongly negative with time. This decrease could be a result of a negative covariance between substitution rates and exon density and/or an increase in the population sizes of the ancestors (if Cov(*ν, X*) *<* 0, as expected since *ν* is relative diversity and *X* is now exon density). As before, positive selection in exons would be expected to produce a positive covariance between exon density and substitution rate, while negative selection would produce a negative covariance. It is hard to determine *a priori* which is likely to be stronger, because although negative selection is thought to be much more ubiquitous, a small amount of positive selection can have a strong effect on substitution rates. The fact that covariance generally goes down with time suggests that negative selection (i.e., constraint) is more strongly affecting substitution rates.

It is at first surprising that the correlations between exon density and divergence go up with time, but the covariances go down with time (Figure 10E,F). However, correlation is defined as Cor(*D_t_, X*) = Cov(*D_t_, X*)*/* SD(*D_t_*) SD(*X*). Thus, if the variance in divergences increases over time the correlations will decrease over time. Indeed, we see this happening as gBGC increases divergences on the ends of the chromosome faster than in the middle, leading to an increase in variance of divergence along the genome. This also explains why correlations of landscapes of very recent times are very noisy, but covariances are not. Indeed, the patterns are clearer when we exclude the tails of the chromosome (Figure S7): there is only a modest increase in the correlation between exon density and divergence over time and the covariances go down with time more linearly.

## 4 Discussion

A central goal of population genetics is to understand the balance of evolutionary forces at work in shaping the origin and maintenance of variation within and between-species (Lewontin, 1974). While the field has been historically data-limited, with the current flood of genome sequencing data, we are poised to make progress on such old questions. Over the past decades, an important lever in understanding the relative impact of genetic drift versus selection in shaping genomic patterns of variation has been to examine the relationship between *levels* of diversity and genomic features, such as recombination rate and exon density. The overarching observation has been that regions of reduced crossing over generally harbor less variation than regions of increased crossing over in many but not all species (e.g., Begun & Aquadro, 1992; Corbett-Detig et al., 2015). This observation is consistent with a role for linked selection shaping patterns of variation in recombining genomes, but the relative contributions of deleterious and beneficial mutations is still largely unknown. Indeed, it seems likely that some complex mixture of both processes shapes variation in natural populations (Kern & Hahn, 2018).

In this paper, we moved beyond genetic diversity within a single species to look at how divergence between closely related species changes with time and how this correlates with genomic features. Previous studies (e.g., Stankowski et al., 2019) looked at similar patterns (in monkeyflowers) and found strong correlations between landscapes of diversity and divergence between related species, despite deep split times. Landscapes of closely related species can remain correlated for two main reasons (i) shared ancestral variation or (ii) shared heterogeneous process. If two species recently split, their landscapes of diversity are expected to be correlated due to shared ancestral variation. If the process that structures genetic diversity along chromosomes is heterogeneous and somewhat shared between-species, then their landscapes are expected to remain correlated over longer periods of time. For example, if the effects of selection are concentrated in the same genomic regions in two species, then their landscapes of diversity will be correlated. By incorporating information from multiple species at once, we are able to pool information across species and thus increase our power to disentangle the role of different evolutionary forces. Patterns across multiple species are more likely to be robust to the idiosyncrasies of any one species, such as demographic history. For instance within-species metrics can be confounded by demography: demographic events can create spurious troughs of diversity (Simonsen et al., 1995) or exacerbate the effects of background selection on diversity (Torres et al., 2018). However, correlations between landscapes can only be produced due to shared ancestral variation or a shared heterogeneous process.

In the great apes, we found that landscapes of within-species diversity and between-species divergence are highly correlated across the phylogeny. Those correlations are often stronger than those that have been historically used as evidence for the effects of selection on genetic variation. For example, the correlation between genetic diversity in humans and exon density is 0.2, yet the correlation between diversity in humans and diversity in western gorillas is 0.48. This stronger correlation may not be entirely due to shared landscape of selection — it may also be a result of shared ancestral variation (and incomplete lineage sorting), mutation rate variation, and/or GC-biased gene conversion. To understand how much of the correlation between landscapes can be attributed to ancestral variation, we performed extensive simulations of the great apes’ evolutionary history, and found that ancestral variation explains very little of the correlations we observed. Thus, a shared heterogeneous process seems to be needed to explain the data.

Two neutral processes can be heterogeneous along the genome and shared across species: GC-biased gene conversion and mutation. GC-biased gene conversion (gBGC) is thought to be an important factor in shaping levels of variation in humans (Chen et al., 2007; Gĺemin et al., 2015; Pouyet et al., 2018), and it has similar effects to those of natural selection. However, if gBGC were a major driver of correlations we would expect to see a difference in overall levels of correlation between different classes of substitution, and we do not (Figures S4 and 6). As such gBGC seems to be a minor contributor to the correlations we observe, although it does seem to be leading to increased substitution rates near the telomeres (where divergences are increasing roughly 5% faster; see Figure 2 and Figure S1). In birds, an excess of divergence near telomeres has been attributed to meiotic drives (Ellegren et al., 2012).

When the history of the great apes is simulated with a shared heterogeneous mutation map, correlations between landscapes do emerge. These were as strong as seen in the data when the rates were drawn from a normal distribution with a standard deviation of the mutation rate of at least a 7% of the mean mutation rate. However, our mutation map was perfectly shared among was species in our simulations, so it is possible that a mutation map which changes over time might move closely to match the data. T. C. A. Smith et al. (2018) estimated the standard deviation of de novo mutation rate in humans at the 1Mb scale to be around 25% of the mean mutation rate. However, the lack of congruency in de novo mutations identified in different data sets raises questions about the role of ascertainment biases that need to be addressed in future studies (Castellano et al., 2020). Our simulations showed a facet of shared mutational heterogeneity along the genome that we do not observe in real data: with variable mutation rate correlations increase over time, whereas in the real data they decrease. It is unknown how conserved mutation rate heterogeneity is across the great apes, so it remains to be seen how an evolving heterogeneous mutation rate map affects landscapes of diversity and divergence. A major driver of mutation rate variation stems from CpG dinucleotides, which have much higher mutation rates than other sites (Agarwal & Przeworski, 2021; Hodgkinson & Eyre-Walker, 2011; Nachman & Crowell, 2000). Nevertheless, when we partitioned the landscapes of divergence by mutation types, we did not see an excess of correlation between landscapes with mutations that can be affected by CpG-induced mutation rate variation (Figures S4 and 6).

Natural selection can also structure genetic variation heterogeneously along the genome. In simulations, both positive and negative selection are needed for the correlations between landscapes to resemble the data. By examining the correlations between landscapes (summarized in Figure 9), we found that the best fitting simulation is the one with a beneficial mutation rate within exons of 1 10*^−^*^12^ and deleterious rate within exons of 1.2 10*^−^*^8^. Positive selection seems to be needed to explain one particular feature of the data: the separation between correlations involving a low *N_e_* species (i.e., correlations are lower if diversity is computed in a species with a low *N_e_*– as seen in humans, bonobos, western chimps and eastern gorillas; see Figure 4). Bottlenecks can erase sweep signatures (Jensen et al., 2005; Nielsen et al., 2005; Przeworski, 2002), but demography does not affect local variation in mutation rates, and it can exacerbate signatures of background selection (Torres et al., 2018). Thus, if sweeps are causing correlations between landscapes, we expect it to be more sensitive to the strong bottleneck in humans than the other processes. This conclusion largely agrees with previous studies which found that positive selection is necessary to explain reduction in genetic diversity surrounding genes in the great apes (Nam et al., 2017).

Another way we might characterize our simulations is through examination of substitution processes. In our best fitting simulation, we get a fixation rate of beneficial mutations of around 1 10*^−^*^9^ per generation per exon base pair, what amounts to approximately 9% of the fixations within exons (along the human lineage), i.e., about one new fixation of a beneficial mutation every 250 generations. Fixation rate within exons is decreased to around 60% of the rate in our neutral simulation due to the constant removal of deleterious mutations within these regions. Indeed, previous studies (Boyko et al., 2008; Laval et al., 2021; Zhen et al., 2021) have estimated that between 10% and 16% of amino acid differences between humans and chimpanzees were caused by positive selection, which is strikingly similar to our best fitting simulation. We would expect to see the fixation of around 16 beneficial mutations in the past 4000 generations, which is close to the number of hard sweeps genome scans for selections have found in humans over this same time period (Schrider & Kern, 2016, 2017). Our best fitting simulation with selection assumes that 60% of new mutations within exons are deleterious, similar to estimates from the site frequency spectrum (Boyko et al., 2008; Huber et al., 2017; B. Y. Kim et al., 2017). Thus while we have not done exhaustive model fitting due to computational constraints, our simulations reproduce estimates from studies which model a different facet of genetic variation (i.e., the site frequency spectrum).

Heterogeneous processes that correlate with a genomic feature will create differences in rates of substitution along the genome that correlate with the genomic feature. As shown in Equation (1), this implies that the covariance along the genome between a genomic feature and divergence is expected to increase with time, and the rate of increase is equal to the covariance between that feature and the substitution rate. (It is important to note that varying covariances with ancestral diversity can be a confounding factor, and that the observation applies to covariance, not correlation.) Indeed, the covariance between divergence and recombination rate increases roughly linearly with time (see Figure 10C), as expected because the rate of gBGC-induced fixations are correlated with recombination rate. Once this effect is removed (see Figure S7F), the covariance between exon density and divergence decreases linearly with time, as we would expect due to the effects of negative selection directly removing deleterious mutations in or near exons. The magnitude of this slope might produce a quantitative estimate of the strength of this effect, although more work is needed to disentangle confounders. It is important to contrast this observation, which applies mostly to the direct effects of selection, to other observations which also include linked effects (as discussed in Phung et al., 2016).

Although simulations allow for more biological realism, we made assumptions to constrain the parameter space explored. Our simulations used randomly mating populations of constant size, as inferred in Prado-Martinez et al. (2013). These population sizes were inferred using neutral model, and so are likely affected by the effects of selection (Jensen et al., 2005). Mean levels of diversity and divergence in our neutral simulation match the data, but simulations with natural selection differ, at times substantially (Figure S9). On the other hand, our simulations with selection match the data more closely with respect to standard deviation in levels of diversity and divergence along genomes. However, inaccuracy of the demographic model should not affect any of our main observations, because the effects of demography on levels of variation along genomes are not shared across multiple species.

We chose exons to be the targets of selection in our simulations. Exons cover about 1% of the human genome, and in reality selection affects non-coding regions as well. However, a substantial portion of this selection affects *cis*-regulatory regions, whose density along the genome is well-predicted by coding sequence itself. Furthermore, highly conserved noncoding sequences have long been identified and characterized as functional (Bejerano et al., 2004; Katzman et al., 2007; Siepel et al., 2005). In the great apes, non-coding diversity is correlated with recombination rate, pointing to the role of selection (Castellano et al., 2020). However, it would be circular to include conserved noncoding elements in our simulations because such elements are identified based in part on levels of divergence, which themselves depend on ancestral levels of genetic diversity. Because conserved noncoding elements generally occur close to coding regions of the genome (at the 1Mb scale, the correlation between density of exons and PhastCons elements is around 0.6), we might expect a more realistic model to have the same amount of selection (in terms of total influx of selected mutations), but spread out over a somewhat wider region of the genome since we have omitted such sites. Even without considering all potential targets of selection, patterns of genetic diversity in our simulations match the data well: we see a correlation between simulated and observed diversity of 0.45 for chimps (Figure S8), for example.

While it has long been recognized that genetic variation among species might be structured similarly due to shared targets of selection, our results demonstrate that these correlations contain important information about the processes at work that has yet to be utilized fully. Here we have used large-scale simulations to demonstrate the combination of forces required to pattern shared divergence and diversity as we observe it in nature. Indeed, our results show that a combination of negative and positive selection, GC-biased gene conversion and mutation rate variation all contribute in shaping genetic variation in the great apes.

Although some processes are not necessarily needed to recapitulate the real data (e.g., mutation rate variation), positive selection seems to be the only force that can explain most of our observations. There is clearly a need for future analytical work that might describe expected correlations across the genome given variation in local mutation, recombination, and selection. Further, statistical model fitting, based on theory or simulation is clearly desirable, although our experience suggests that the latter approach would prove computationally expensive.

## Acknowledgements

We thank the Kern-Ralph CoLab for their invaluable support and input. Comments from reviewers also significantly improved this version of our paper. This work was supported by NIH awards R01GM117241 and R01HG010774 to A.D.K.

## Supplementary material

### 4.1 Correlation between divergences that share branches

Landscapes of divergence can be correlated by their definition, as they can share part of their histories. In most of our analyses (except for Figure S2), we do not show the correlations for such cases but below we describe how this sharing would affect correlations (using a simplified theory). For example, in Figure 3 *d_V X_* and *d_XY_* share the branch *X*; depending on how the length of the branch *X* compares to the total tree length, these two landscapes are bound to be correlated. Assuming that mutations follow a Poisson process and that coalescences happen instantaneously, we derive the following. There are three non-overlapping parts in the tree between these, the branch from the *XY* ancestor to *X* with length E[*τ_X_*] = *T_XY_*, the branch from the *XY* ancestor to *Y* with length E[*τ_Y_*] = *T_XY_* and the branch from *V* to the *XY* ancestor with length E[*τ_V_*] = 2*T_V W XY_ − T_XY_*. If we just consider the genealogical definition of divergence and assume *d_V X_* = *τ_V_* + *τ_X_* and *d_XY_* = *τ_X_* + *τ_Y_* (i.e., ignoring the contributions of ancestral diversity to divergence), then

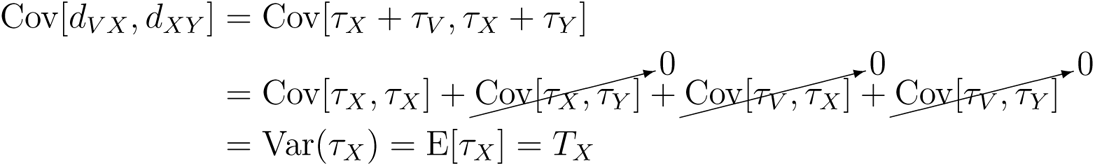

Therefore,

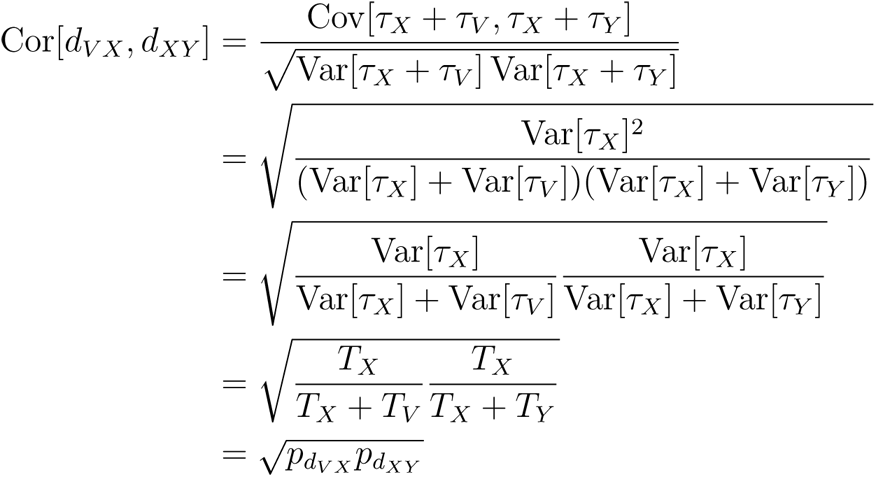

where 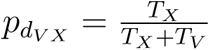 is the proportion of *d_V X_* that is shared with *d_XY_*, and 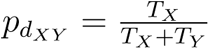 is the proportion of *d_XY_* that is shared with *d_V X_*.

---

**Figure S1:**
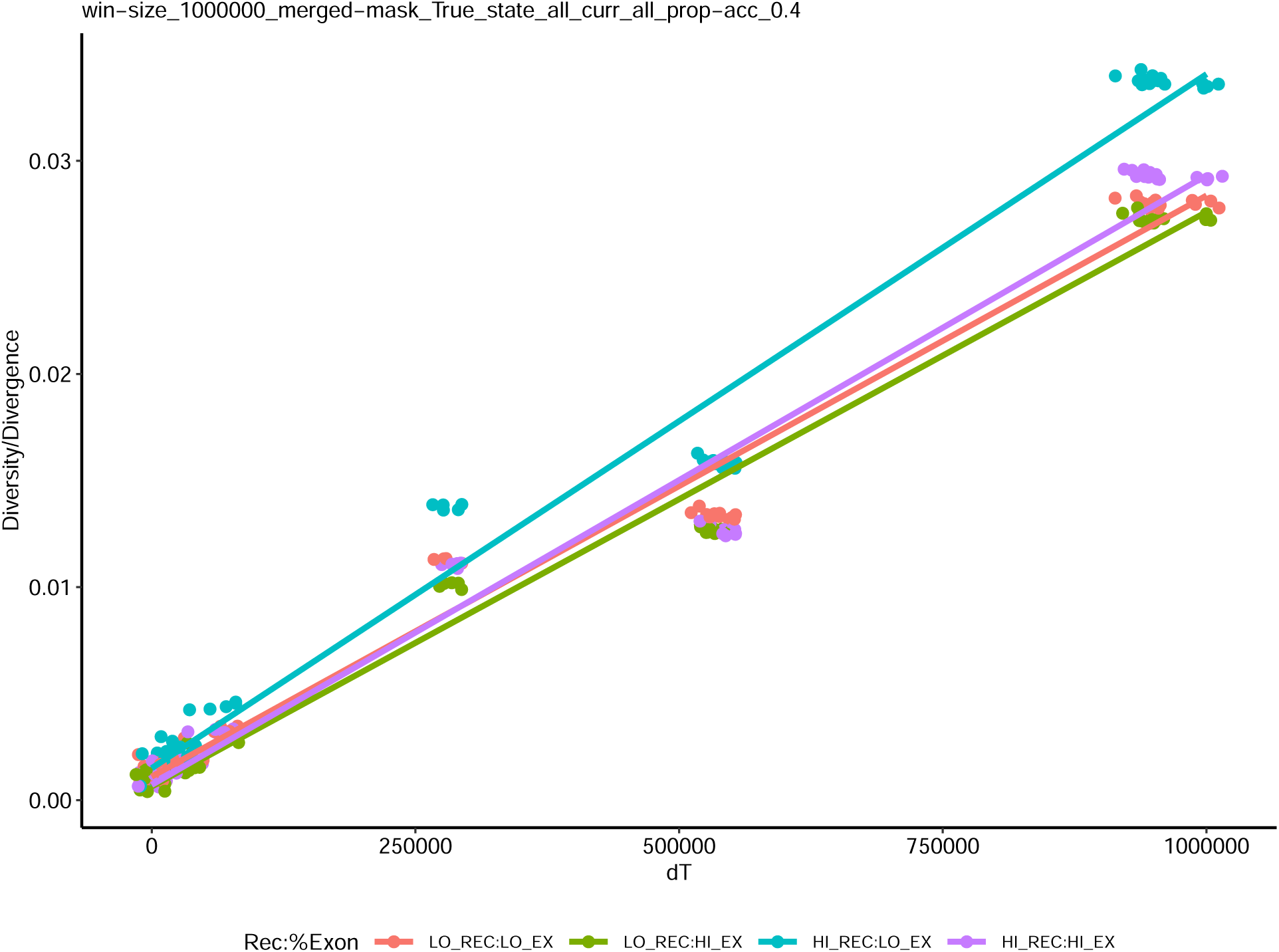
Effect of exon density and recombination rate on the accumulation of genetic divergence in chromosome 12 with phylogenetic distance. Within-species genetic diversities are shown at *dT* = 0. Mean diversity and divergences were computed for four groups depending on whether they fell or not on the top 90% percentile of recombination rate and exon density.

**Figure S2:**
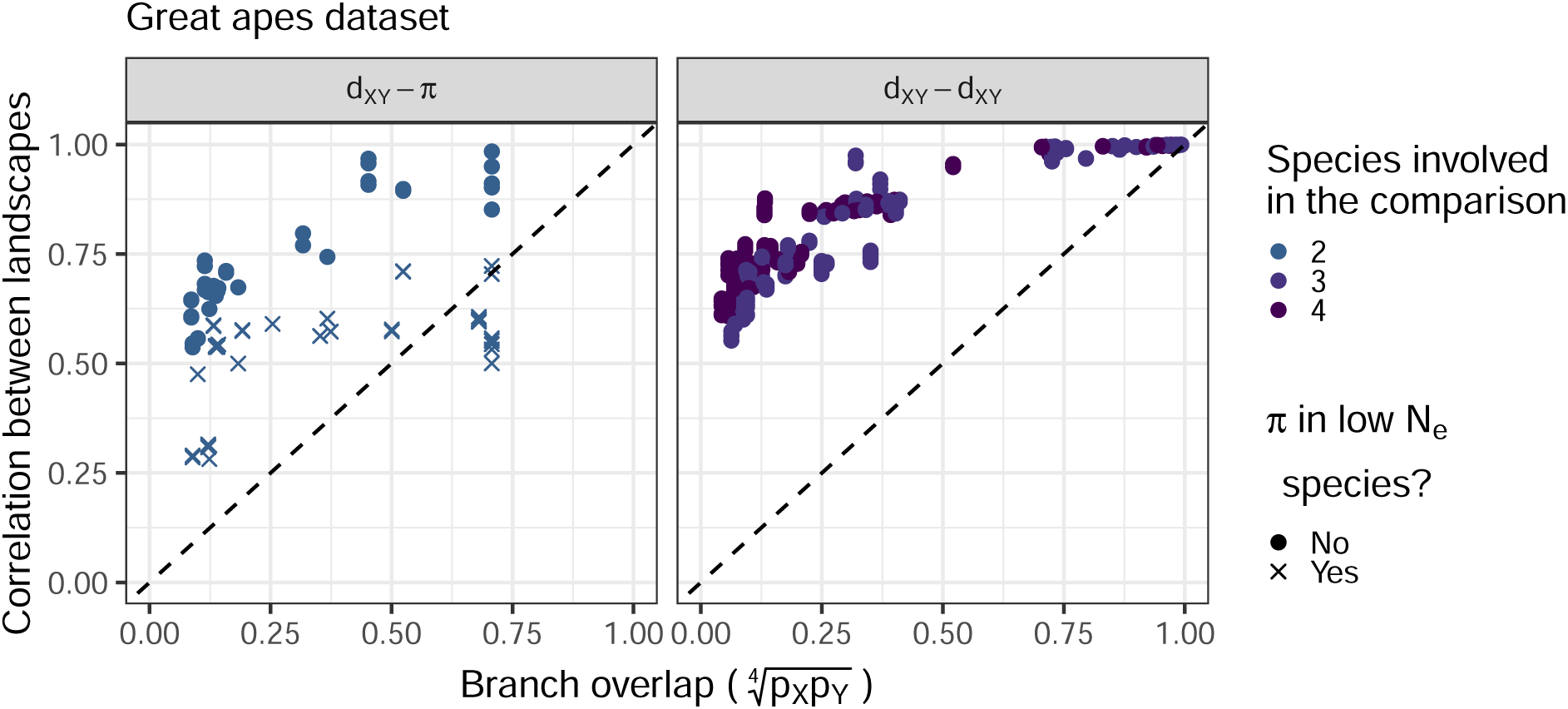
For example, diversity in humans and divergence between humans and bonobos share part of their history. Each point on the plots correspond to the (Spear-man) correlation between two landscapes of diversity/divergence, computed on 1Mb windows across the entire genome. Correlations were split by type of landscapes compared (*π − d_XY_*, *d_XY_ − d_XY_*). The x-axis is a metric of expected branch overlap between the landscapes. See subSection 4.1 for more information. Note that species with low *N_e_* (bonobos, eastern gorillas and western chimps) have a different point shape. The colors reflect the number of species involved in the comparison. For example, the comparison between human-western gorilla and eastern chimp-Sumatran orangutan divergences includes four different species. On the other hand, the comparison between human-western gorilla and human-Sumatran orangutan divergences includes just three species.

**Figure S3:**
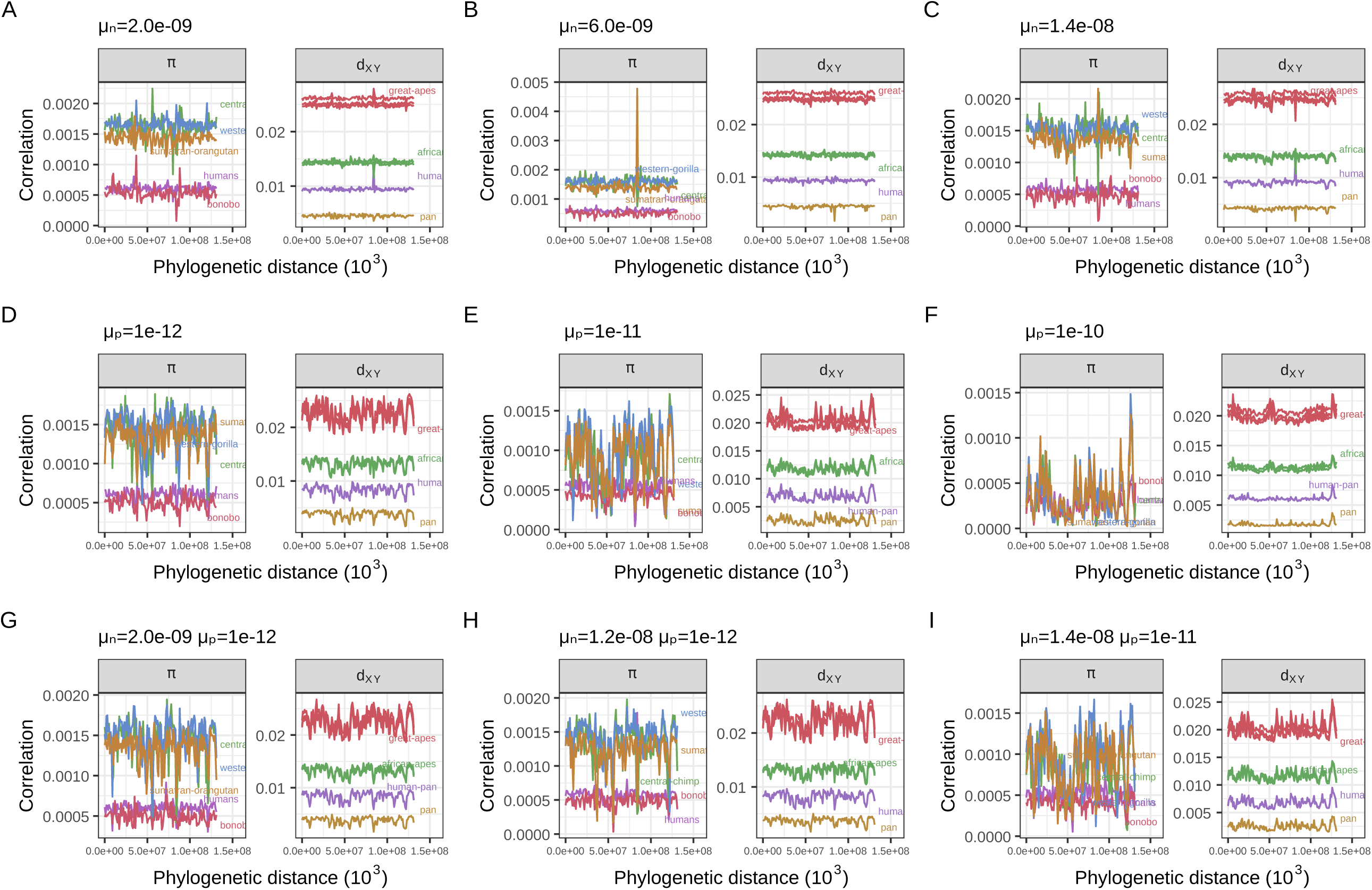
Landscapes of diversity and divergence in selected simulations with natural selection. The selection parameters *µ_n_* and *µ_p_*are the rate of mutations in exons with negative and positive fitness effects, respectively. The mean fitness effect was *s̄* = 0.03 for deleterious mutations and *s̄* = 0.01 for beneficial mutations (see subsection 2.2 for more details). Other details are as in Figure 2.

**Figure S4:**
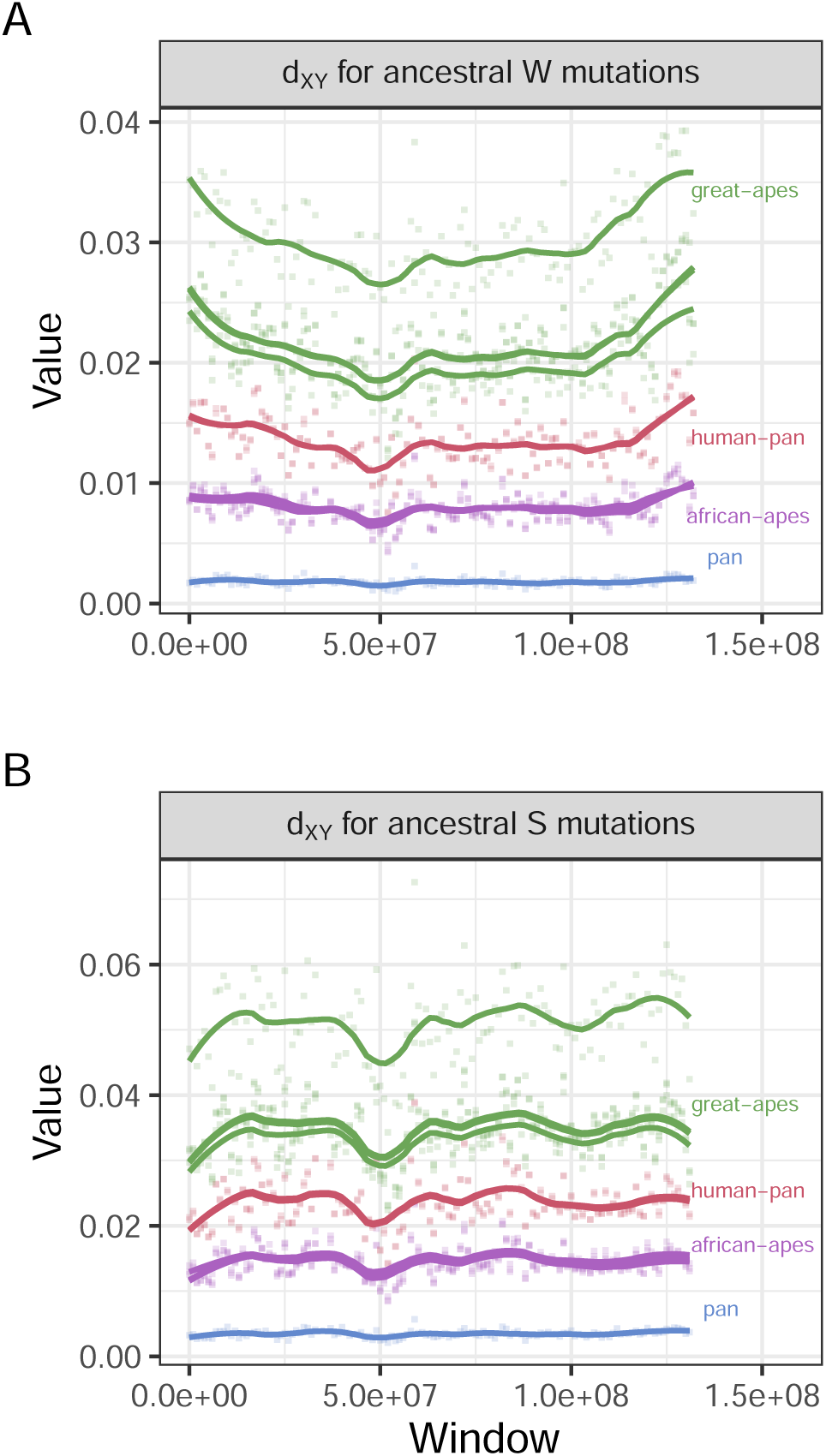
Landscapes of divergence partitioned by allele state in the ancestor. Ancestral states were assumed to be the same as seen in rhesus macaques (RheMac2), and sites not called in macaques were not used. *d_XY_* for W sites is simply the mean pairwise differences between samples in species *X* and *Y* per ancestral W sites (A/T). Similar reasoning applies for *d_XY_* for S ancestral sites, but only considering (G/C) sites. Points were colored by the most common recent ancestor of the two species compared in each divergence. Lines were fitted using local linear regression. Note that for ancestrally weak mutations (A) there is an increase in divergence at the ends of the chromosomes, but that is not seen for ancestrally strong mutations (B).

**Figure S5:**
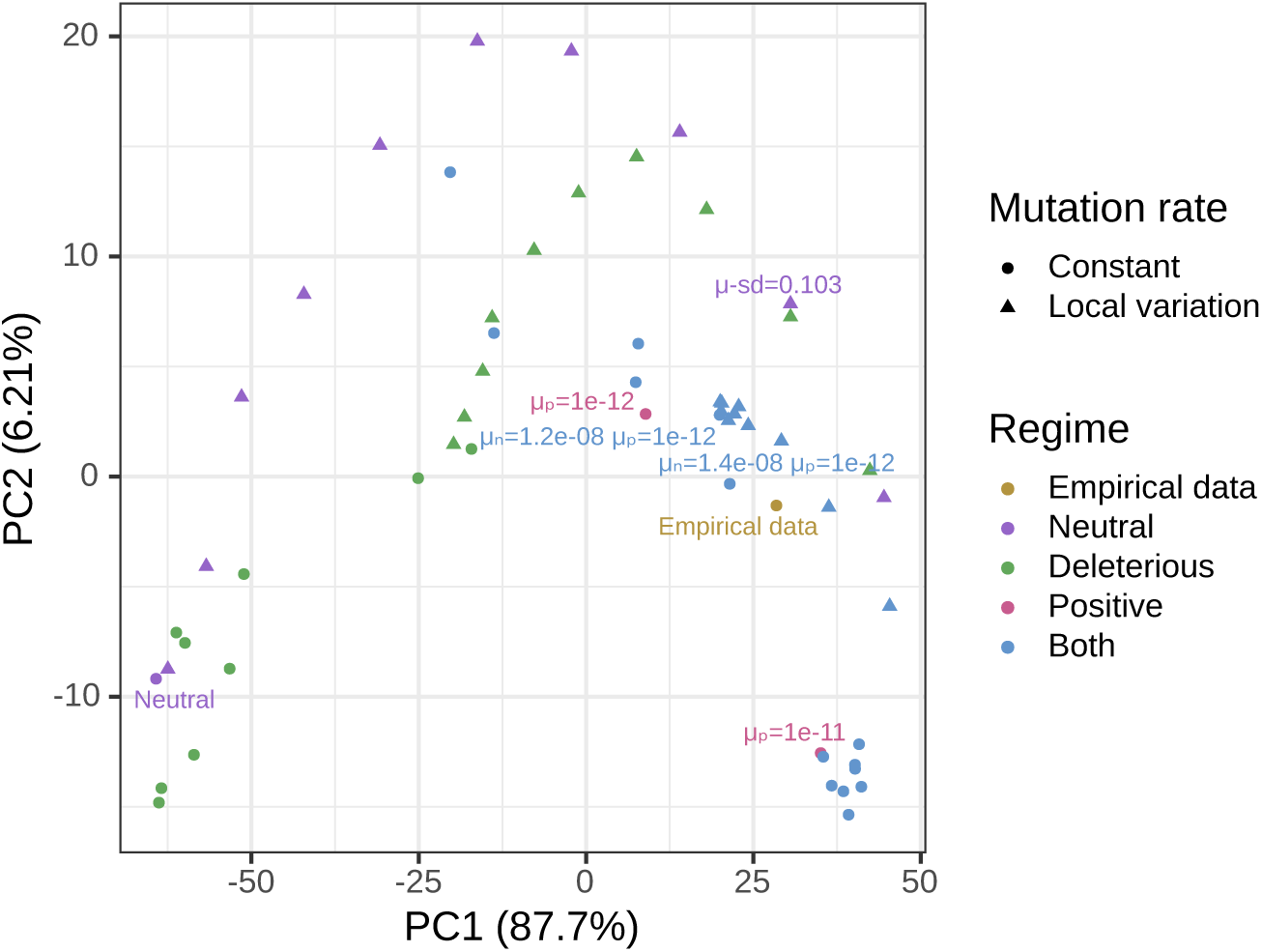
PCA visualization of data and simulations at 500Kb. The colors differentiate the empirical data from simulations with different parameters: Neutral refers to the simulation without any selection, Deleterious refers to simulations with deleterious mutations, Positive refers to simulations with beneficial mutations, Both refers to simulations with both beneficial and deleterious mutations. The shape of the points differentiate simulations with constant mutation rate along the genome and variable local mutation rates. Principal component analysis (PCA) applied to a matrix with all pairwise correlations between landscapes across the great apes (including *π π*, *π d_XY_* and *d_XY_ d_XY_* comparisons) for the great apes dataset and simulations (with selection and with mutation rate variation). We excluded simulations with *µ_p_* 1 10*^−^*^10^ from the PCA analysis because PC2 was capturing negative correlations caused by strong positive selection — as seen in Figure 7F.

**Figure S6:**
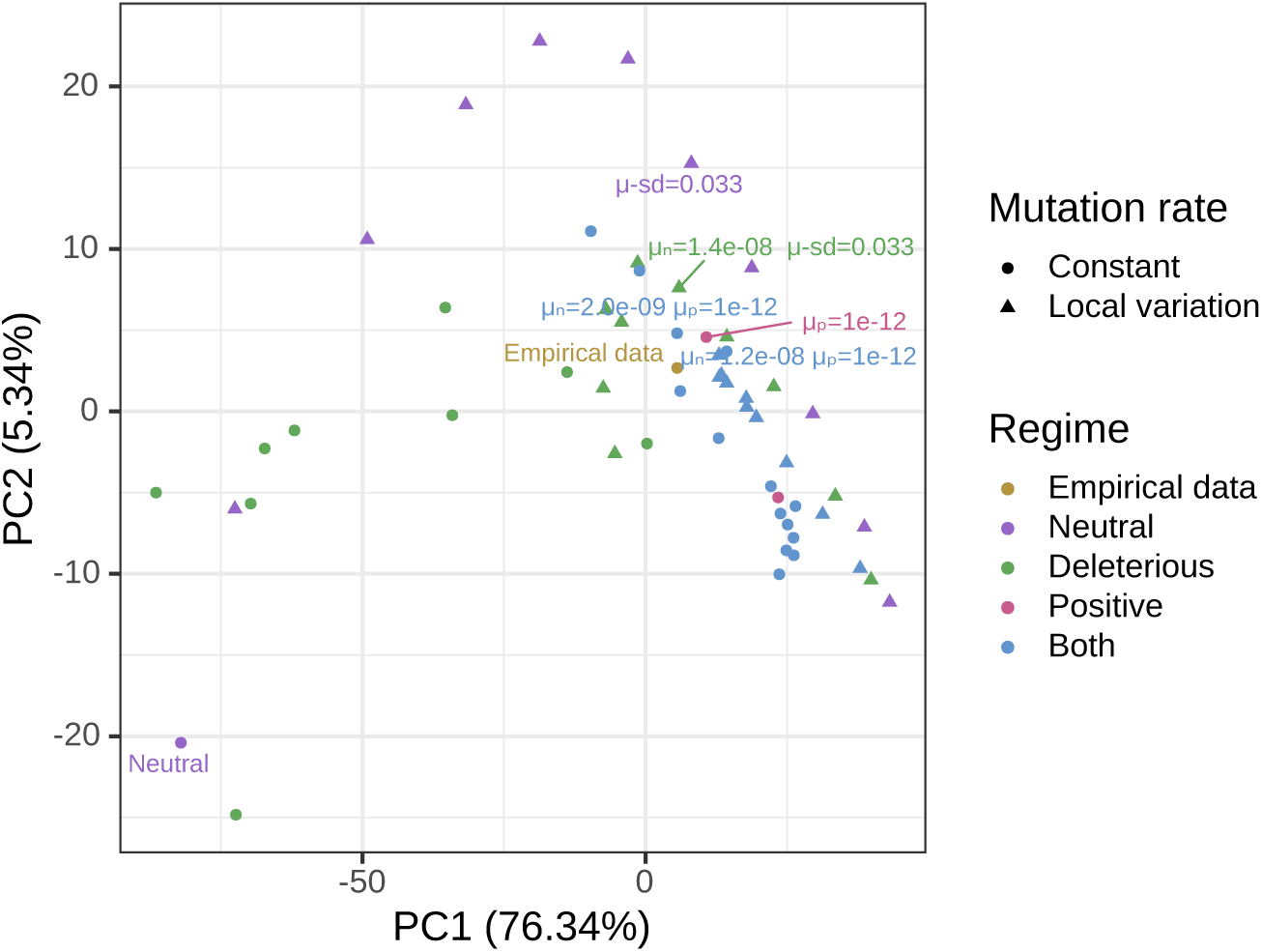
PCA visualization of data and simulations at 5Mb. The colors differentiate the empirical data from simulations with different parameters: Neutral refers to the simulation without any selection, Deleterious refers to simulations with deleterious mutations, Positive refers to simulations with beneficial mutations, Both refers to simulations with both beneficial and deleterious mutations. The shape of the points differentiate simulations with constant mutation rate along the genome and variable local mutation rates. Principal component analysis (PCA) applied to a matrix with all pairwise correlations between landscapes across the great apes (including *π π*, *π d_XY_* and *d_XY_ d_XY_* comparisons) for the great apes dataset and simulations (with selection and with mutation rate variation). We excluded simulations with *µ_p_* 1 10*^−^*^10^ from the PCA analysis because PC2 was capturing negative correlations caused by strong positive selection — as seen in Figure 7F.

**Figure S7:**
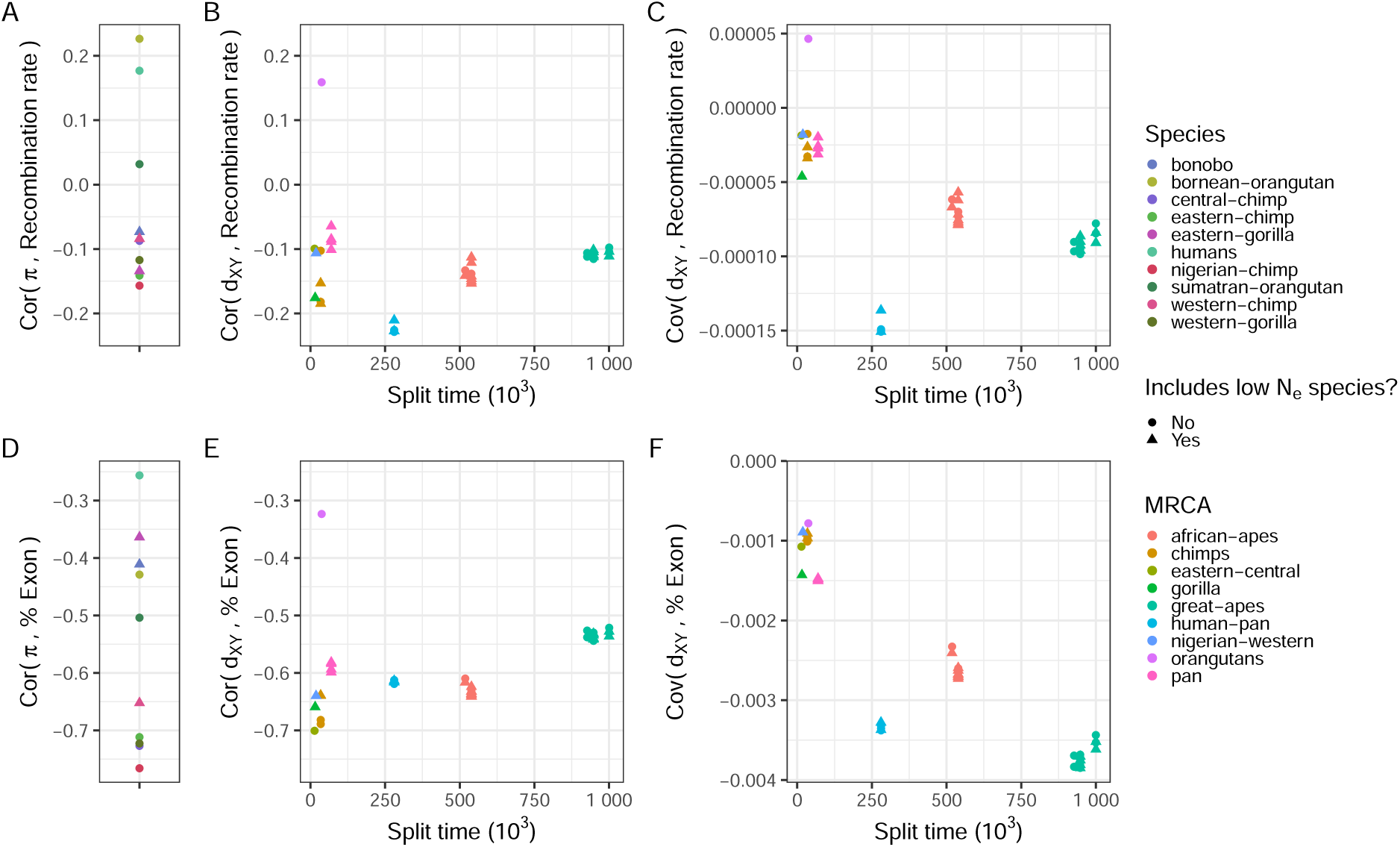
Correlations and covariances between landscapes of diversity and divergence and annotation features in the real great apes data. Only windows in the middle half of chromosome 12 were included. Compare to Figure 10.

**Figure S8:**
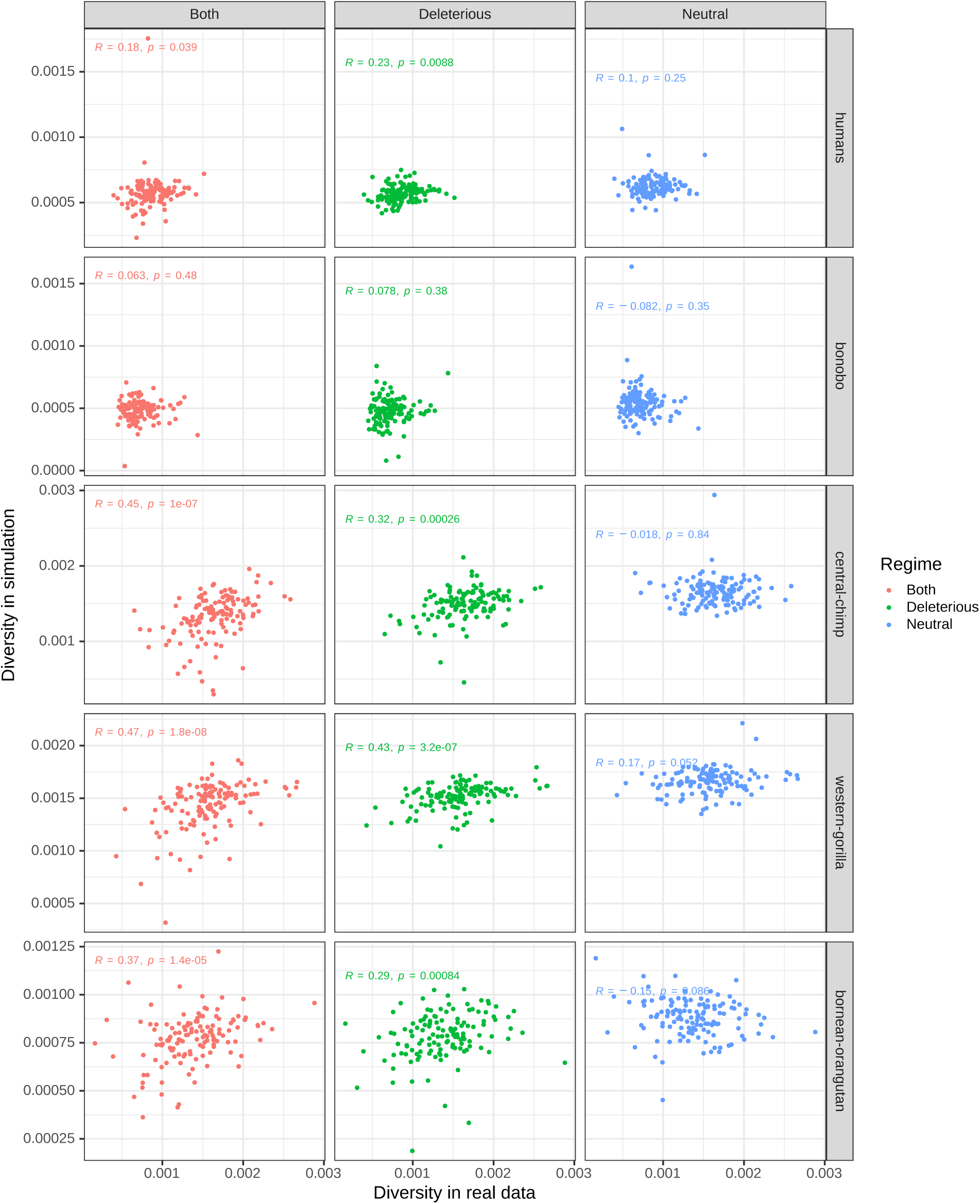
Scatterplots of genetic diversity in the real great apes data against diversity seen in simulations. Simulation with deleterious mutations had *µ_n_* = exp 1.4*−*8, and the simulation with both deleterious and beneficial mutations had *µ_n_*= exp 1.4*−*8 and *µ_p_* = exp 1*−*12.

**Figure S9:**
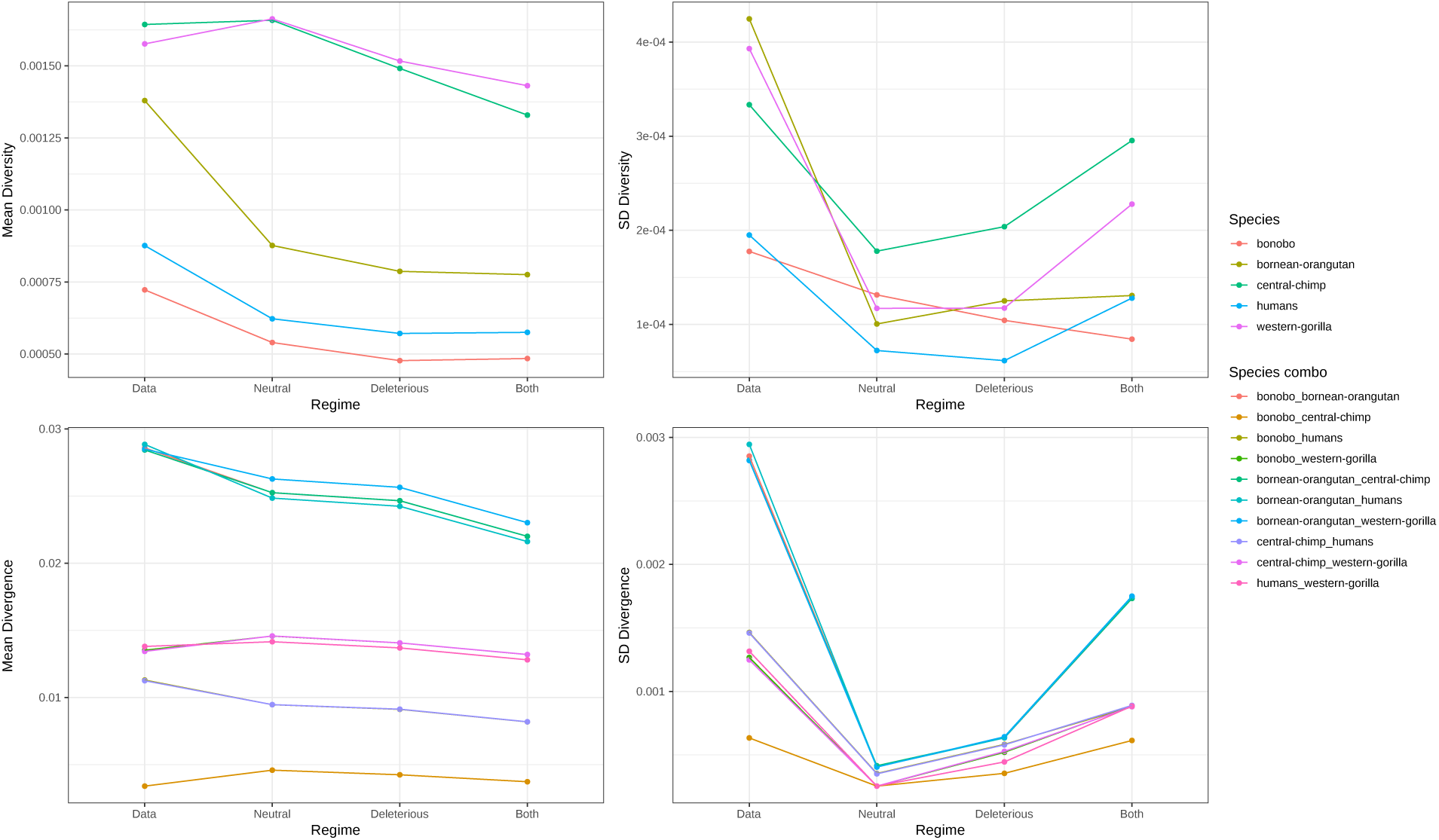
Summaries of genetic diversity (top) and divergence (bottom) across all species for the data and simulations. Mean is shown on the left and standard deviation on the right. “Neutral” refers to the simulation without any selection, “Deleterious” refers to the simulation with deleterious mutations ocurring at a rate of 1.4 10*^−^*^8^, “Both” refers to the simulation with both beneficial and deleterious mutations, with rates 1 10*^−^*^12^ and 1.4 10*^−^*^8^ respectively.

**Figure S10:**
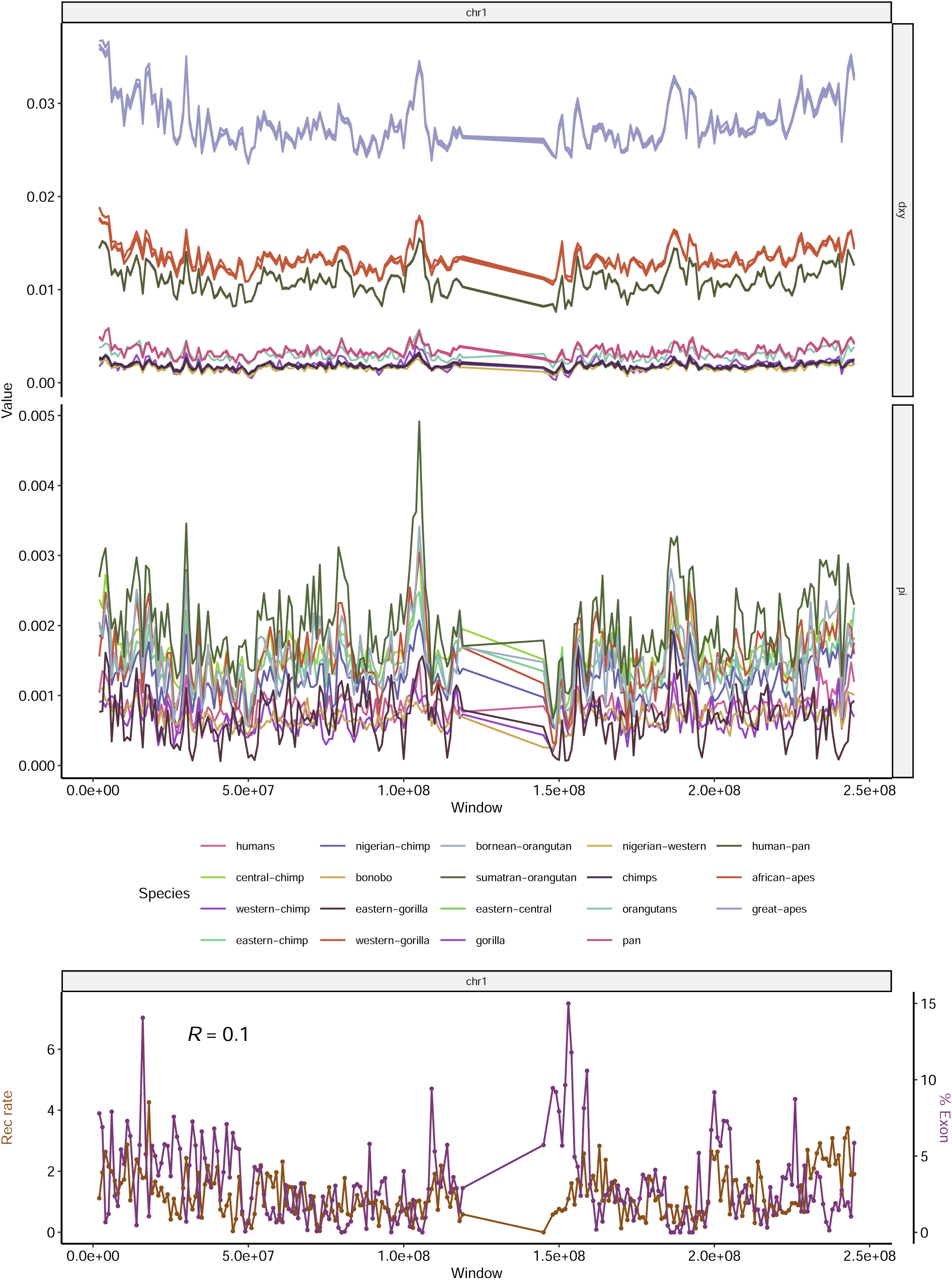
Landscapes of diversity, divergence, exon density and recombination rate across chromosome 1. See Figure 2 for more details.

**Figure S11:**
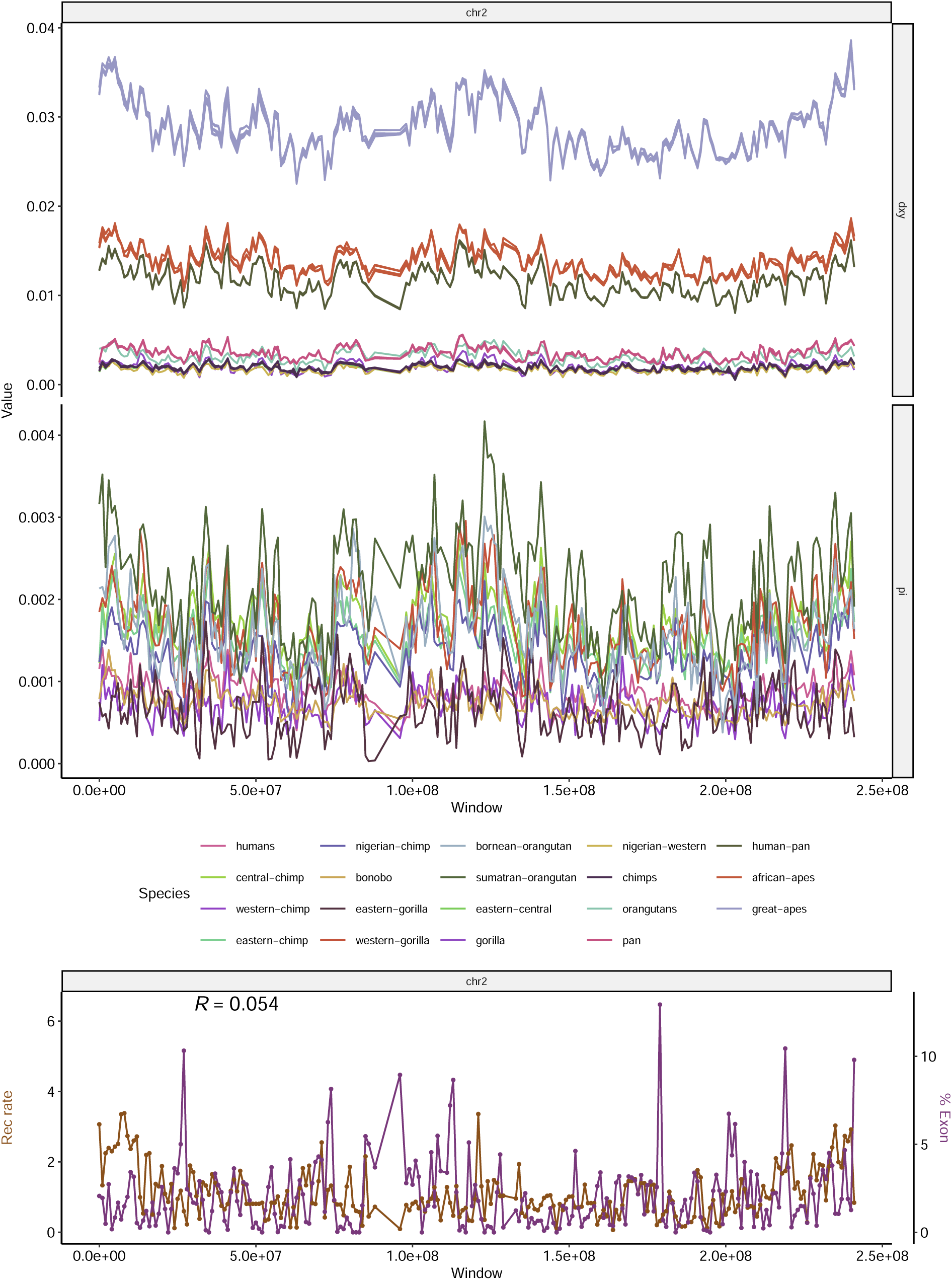
Landscapes of diversity, divergence, exon density and recombination rate across chromosome 2. See Figure 2 for more details.

**Figure S12:**
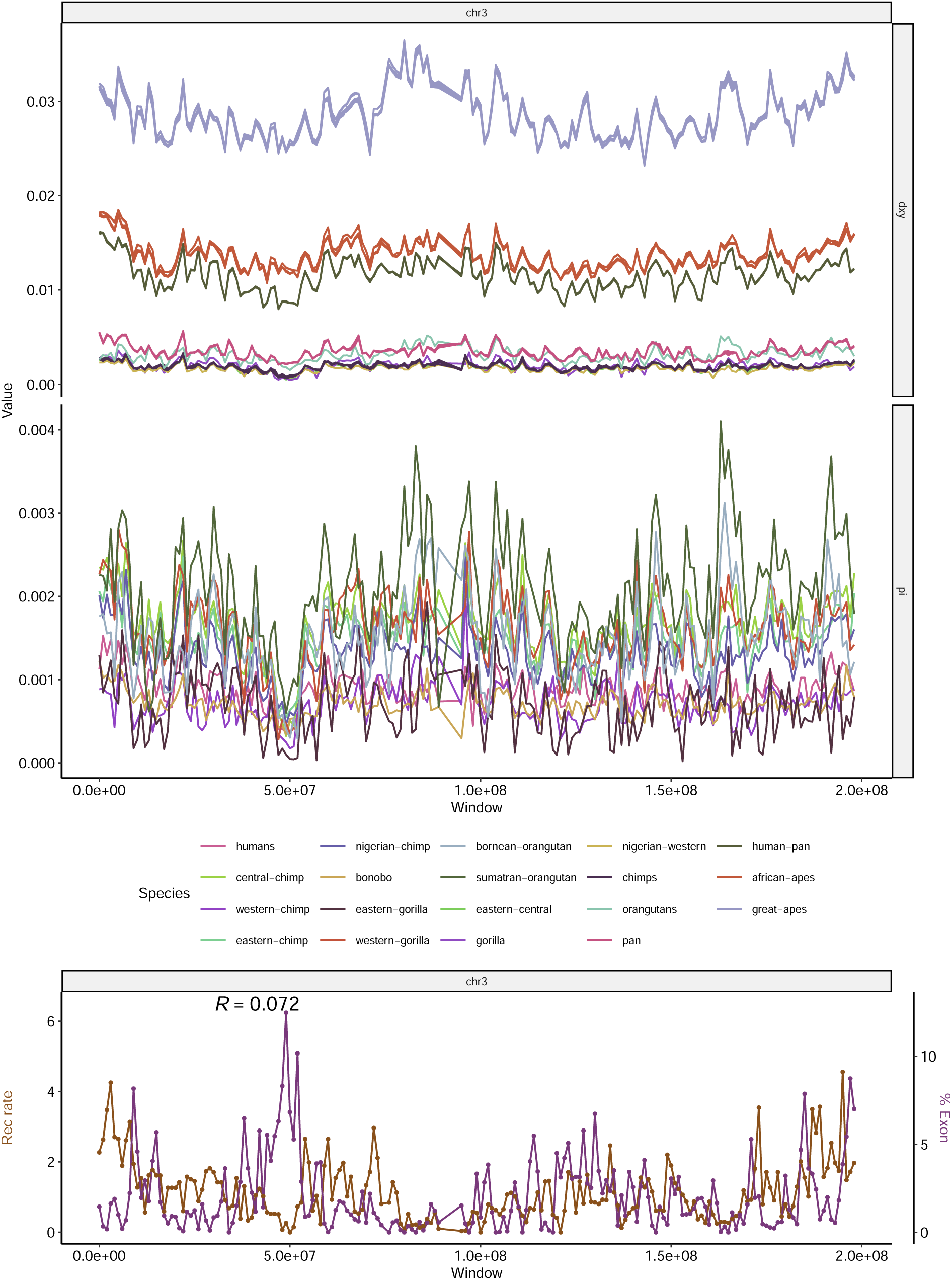
Landscapes of diversity, divergence, exon density and recombination rate across chromosome 3. See Figure 2 for more details.

**Figure S13:**
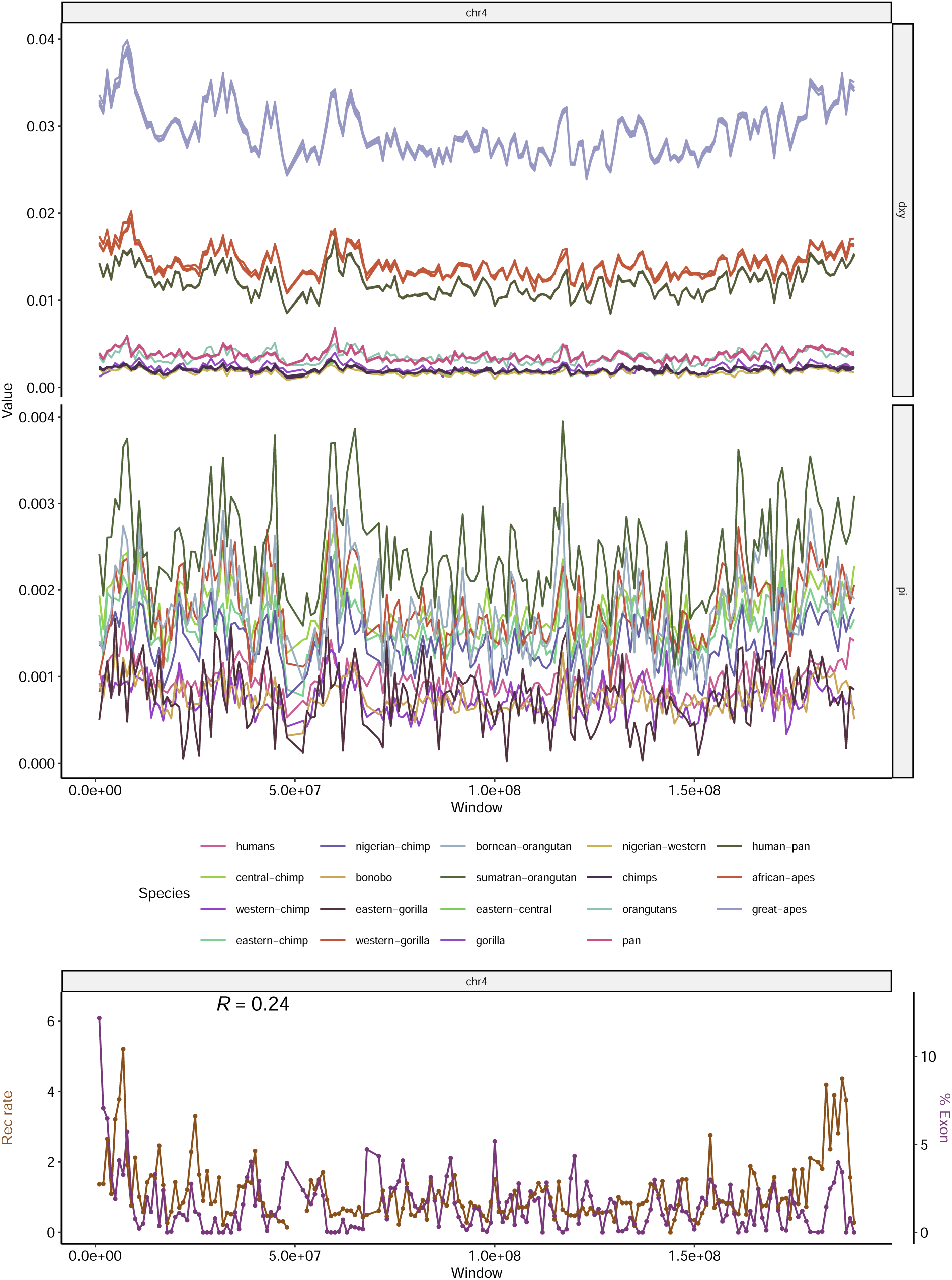
Landscapes of diversity, divergence, exon density and recombination rate across chromosome 4. See Figure 2 for more details.

**Figure S14:**
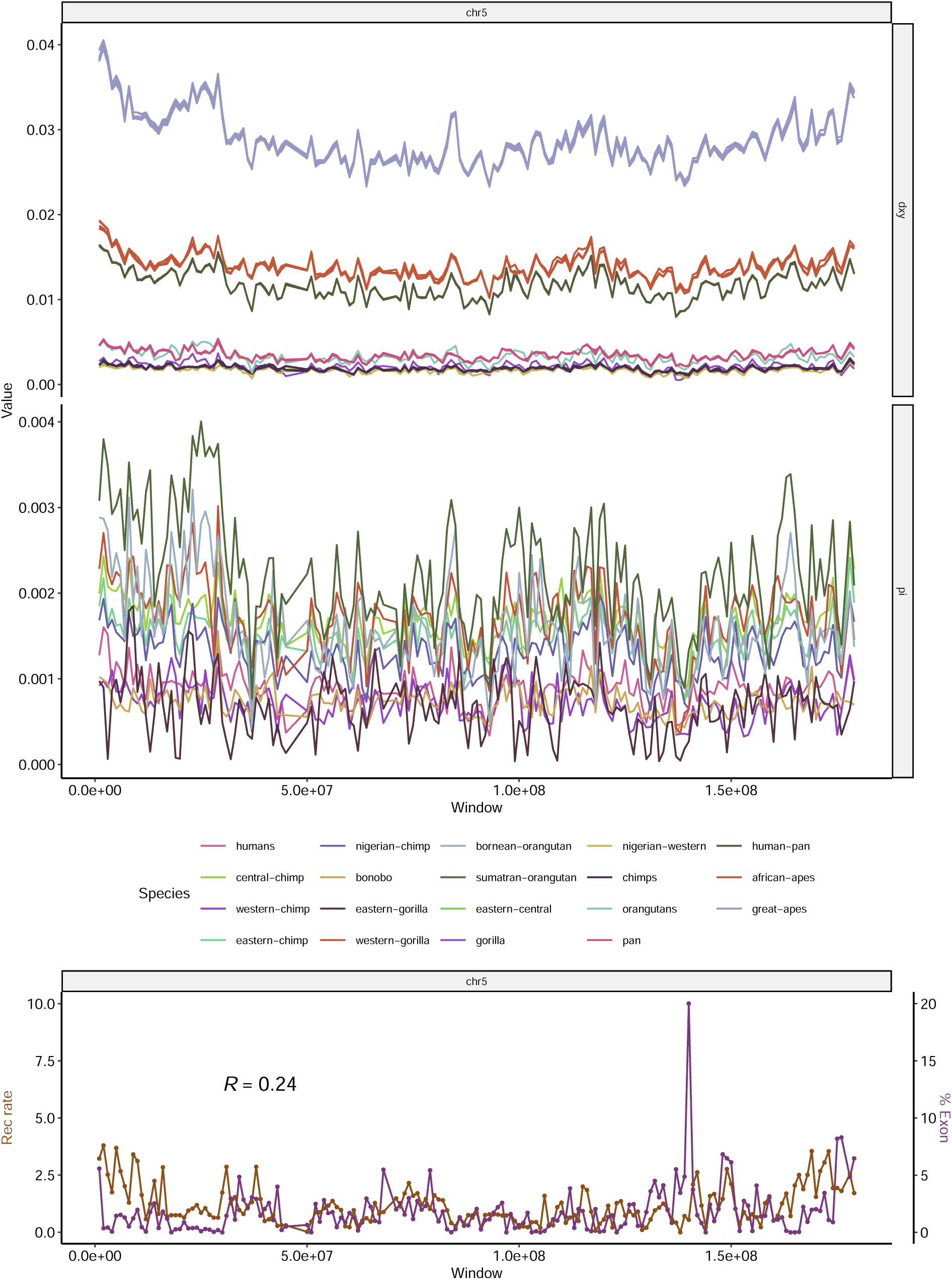
Landscapes of diversity, divergence, exon density and recombination rate across chromosome 5. See Figure 2 for more details.

**Figure S15:**
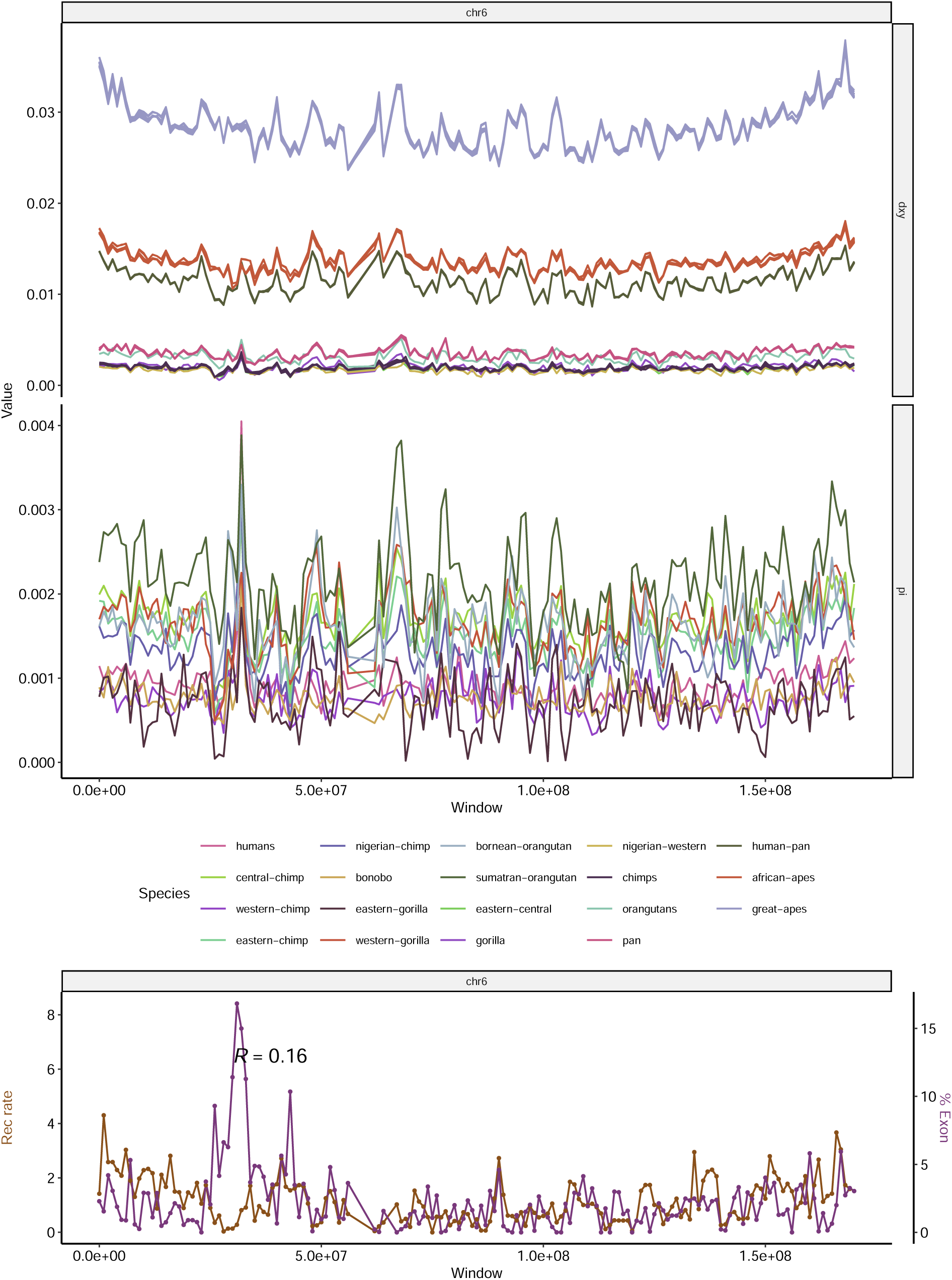
Landscapes of diversity, divergence, exon density and recombination rate across chromosome 6. See Figure 2 for more details.

**Figure S16:**
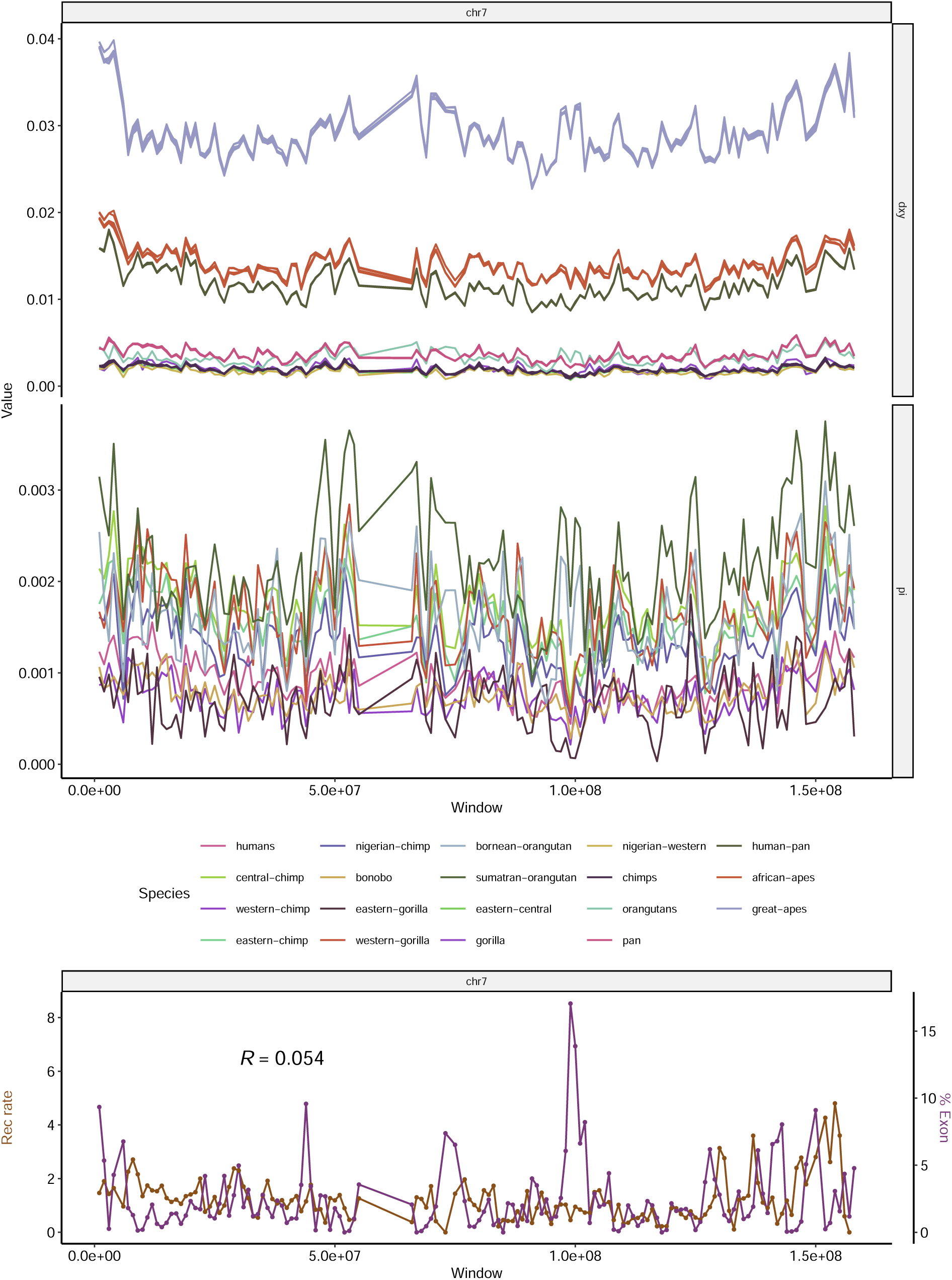
Landscapes of diversity, divergence, exon density and recombination rate across chromosome 7. See Figure 2 for more details.

**Figure S17:**
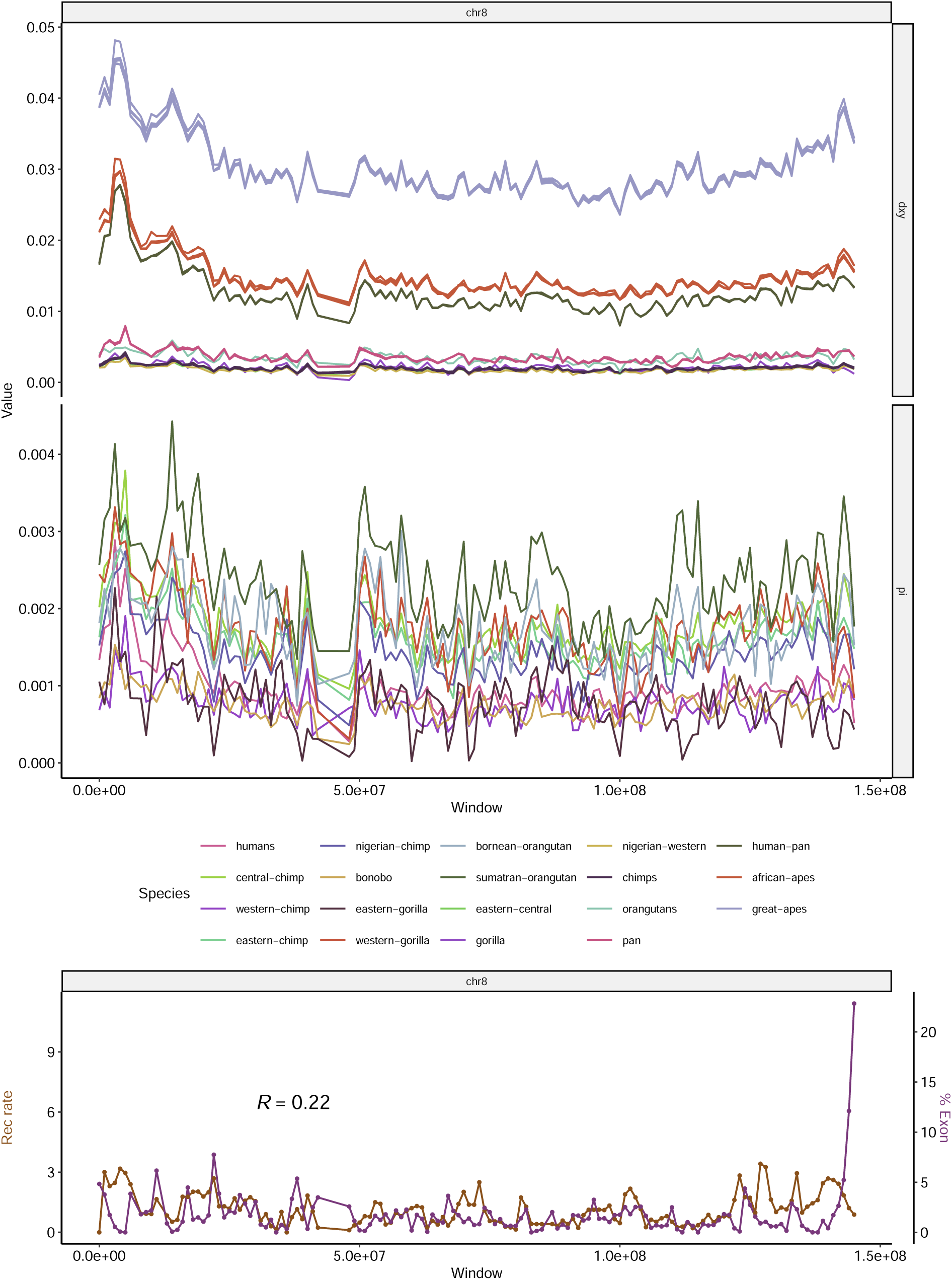
Landscapes of diversity, divergence, exon density and recombination rate across chromosome 8. See Figure 2 for more details.

**Figure S18:**
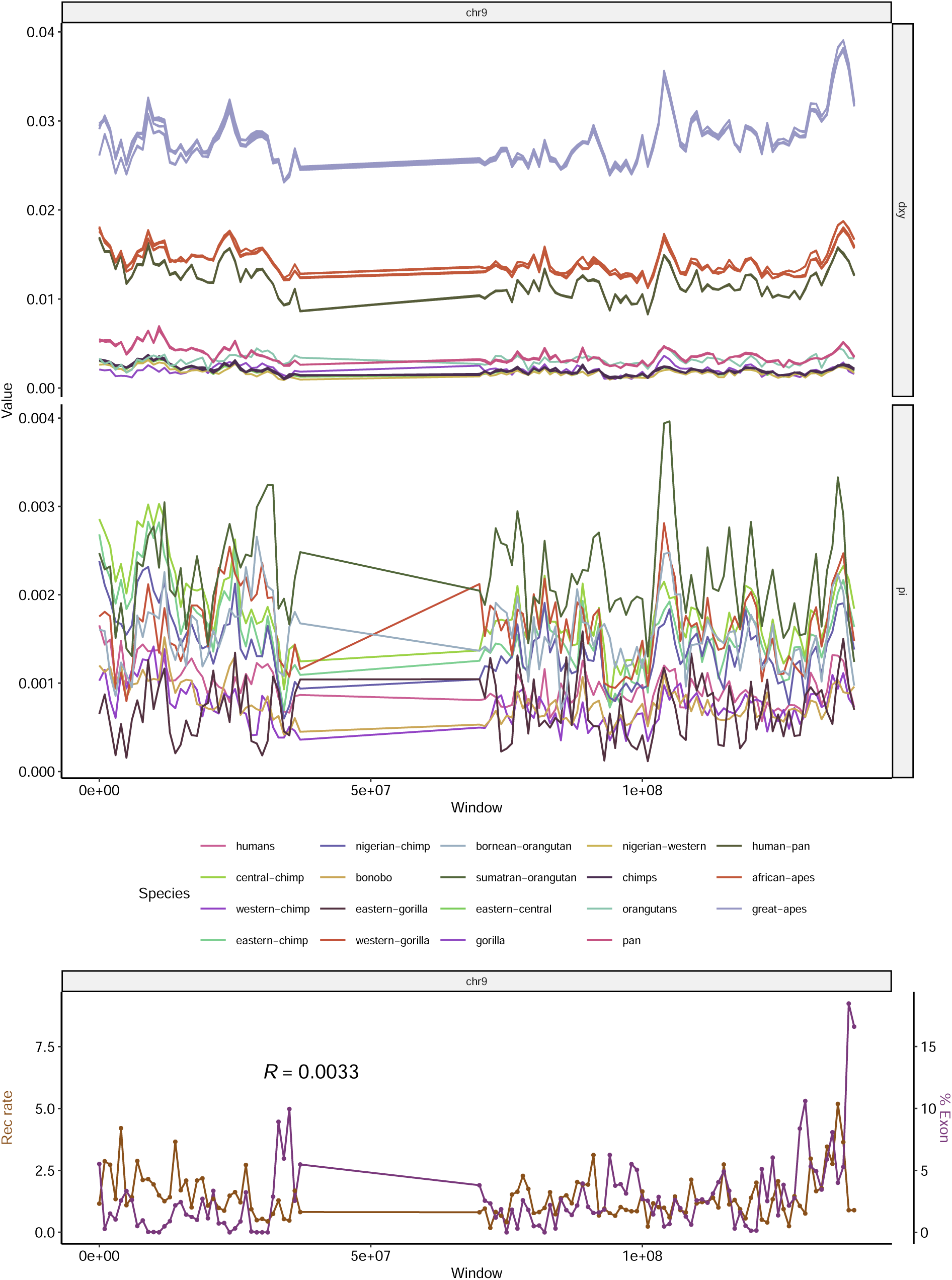
Landscapes of diversity, divergence, exon density and recombination rate across chromosome 9. See Figure 2 for more details.

**Figure S19:**
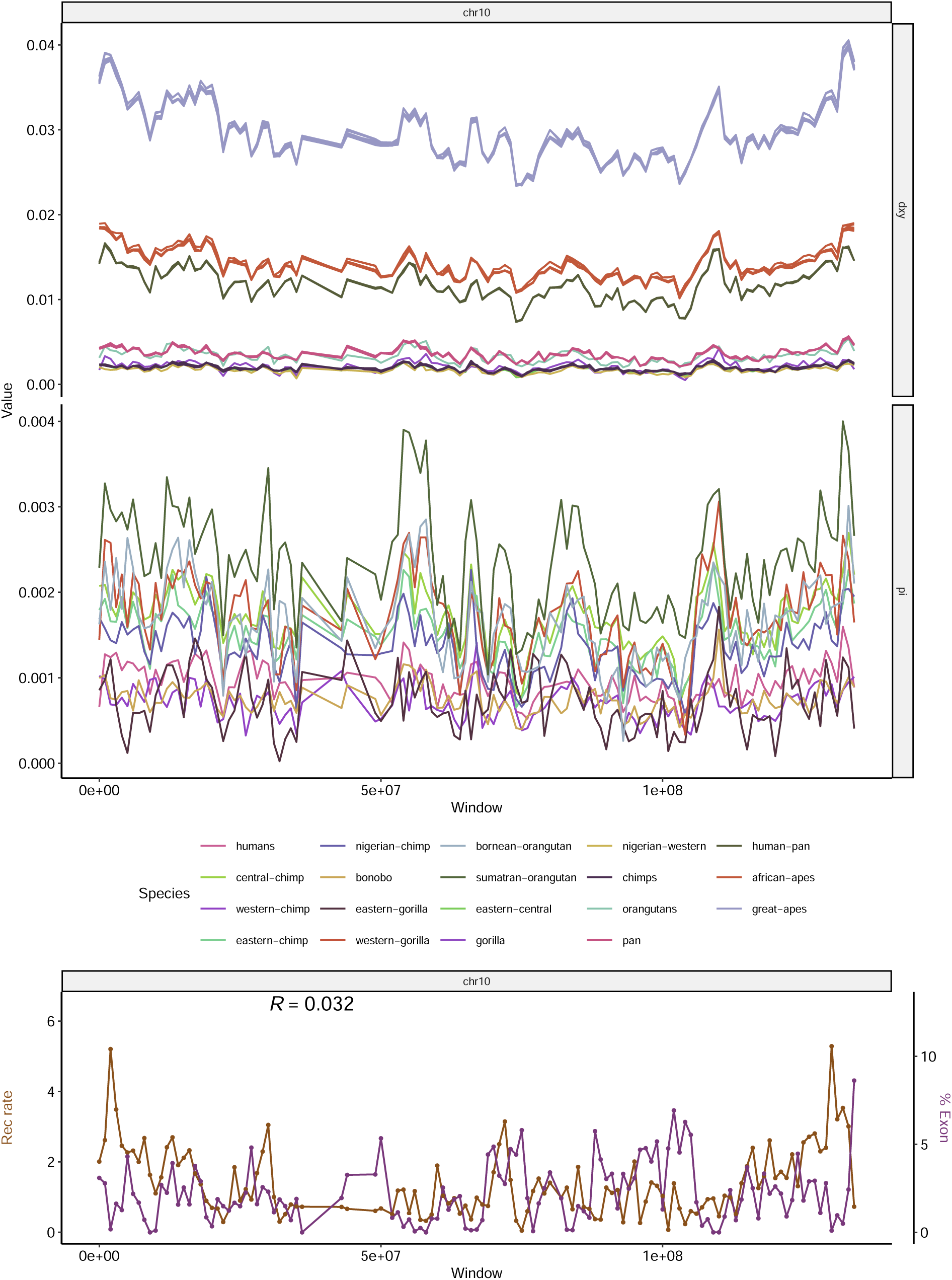
Landscapes of diversity, divergence, exon density and recombination rate across chromosome 10. See Figure 2 for more details.

**Figure S20:**
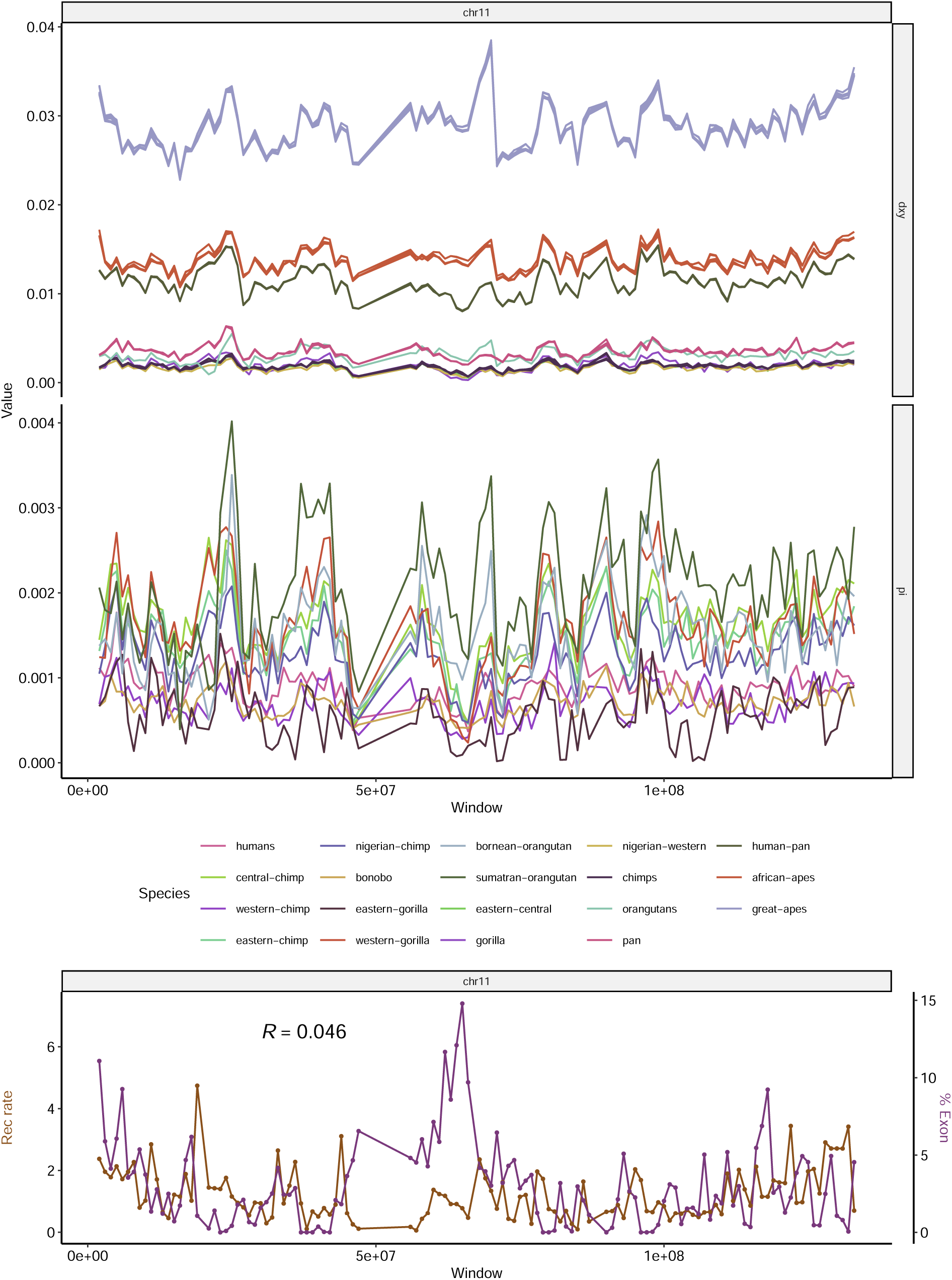
Landscapes of diversity, divergence, exon density and recombination rate across chromosome 11. See Figure 2 for more details.

**Figure S21:**
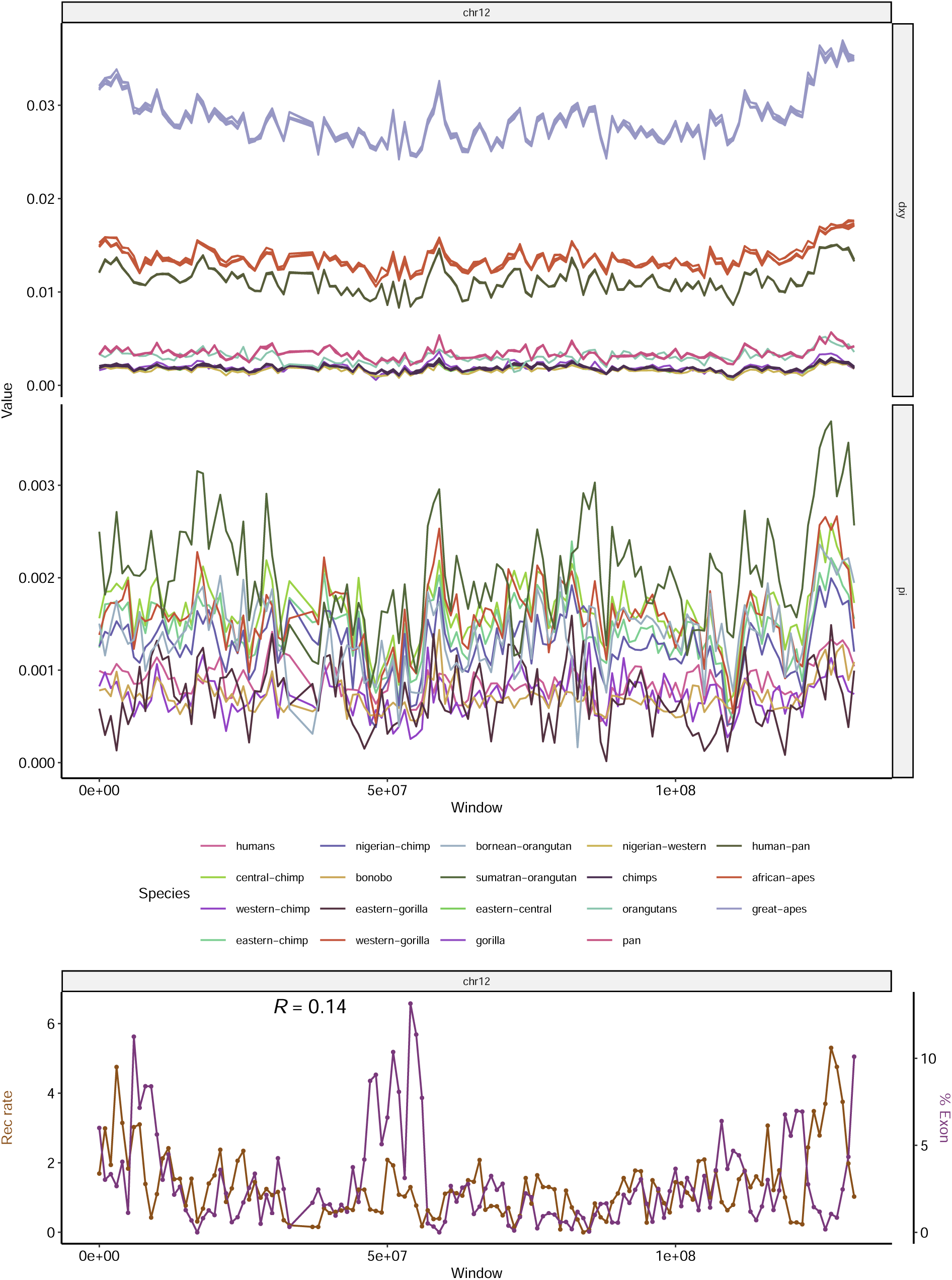
Landscapes of diversity, divergence, exon density and recombination rate across chromosome 12. See Figure 2 for more details.

**Figure S22:**
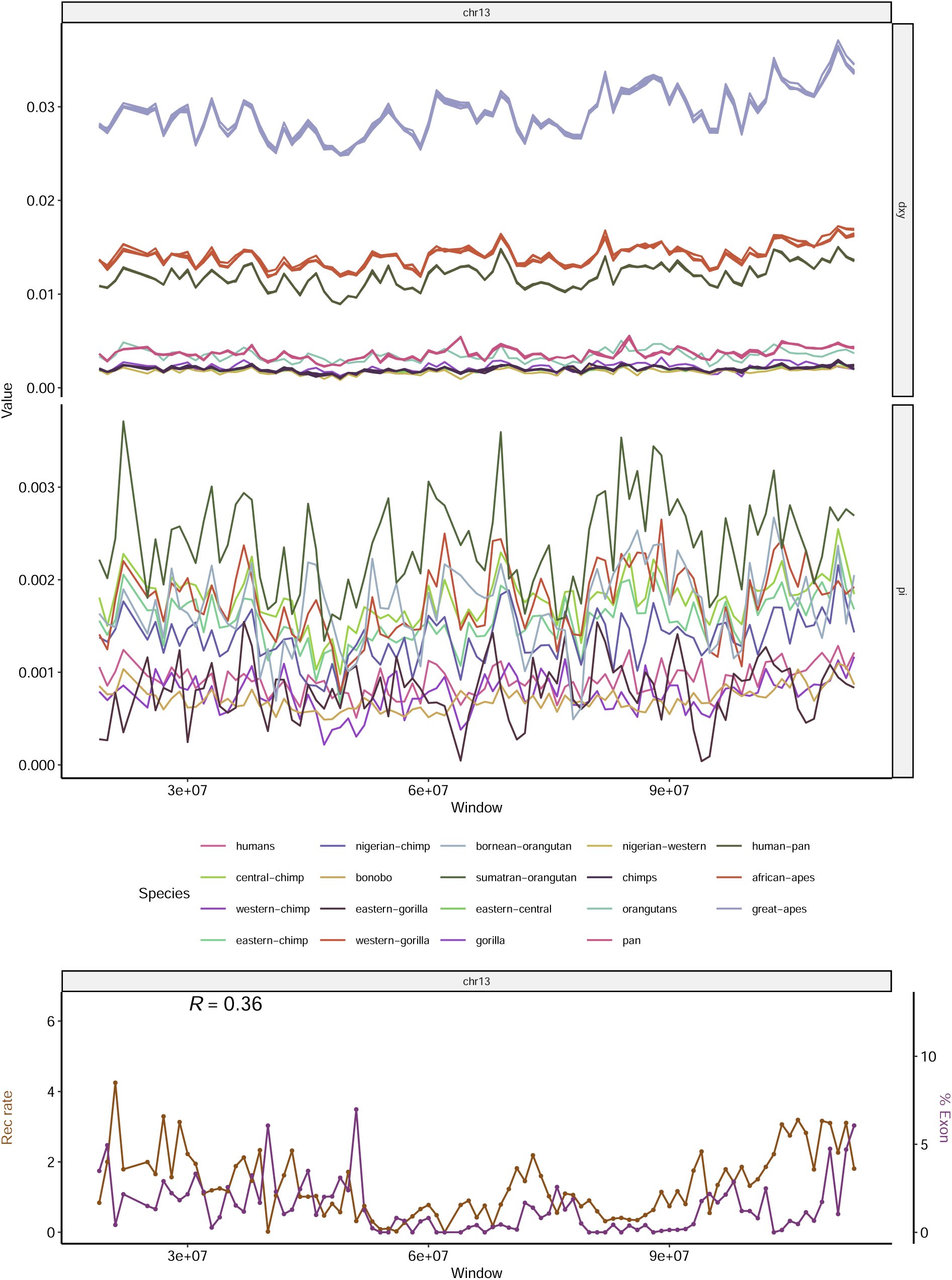
Landscapes of diversity, divergence, exon density and recombination rate across chromosome 13. See Figure 2 for more details.

**Figure S23:**
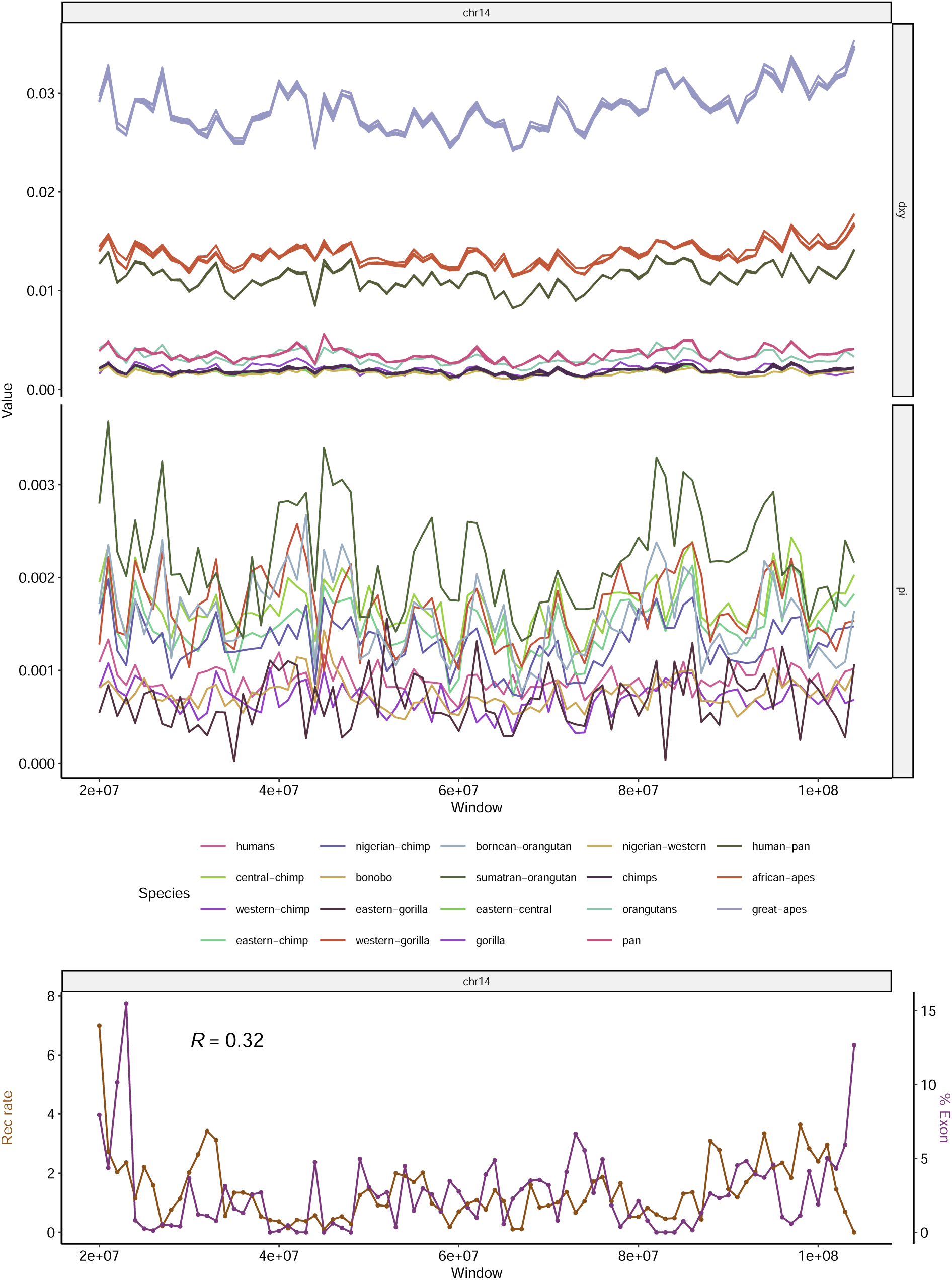
Landscapes of diversity, divergence, exon density and recombination rate across chromosome 14. See Figure 2 for more details.

**Figure S24:**
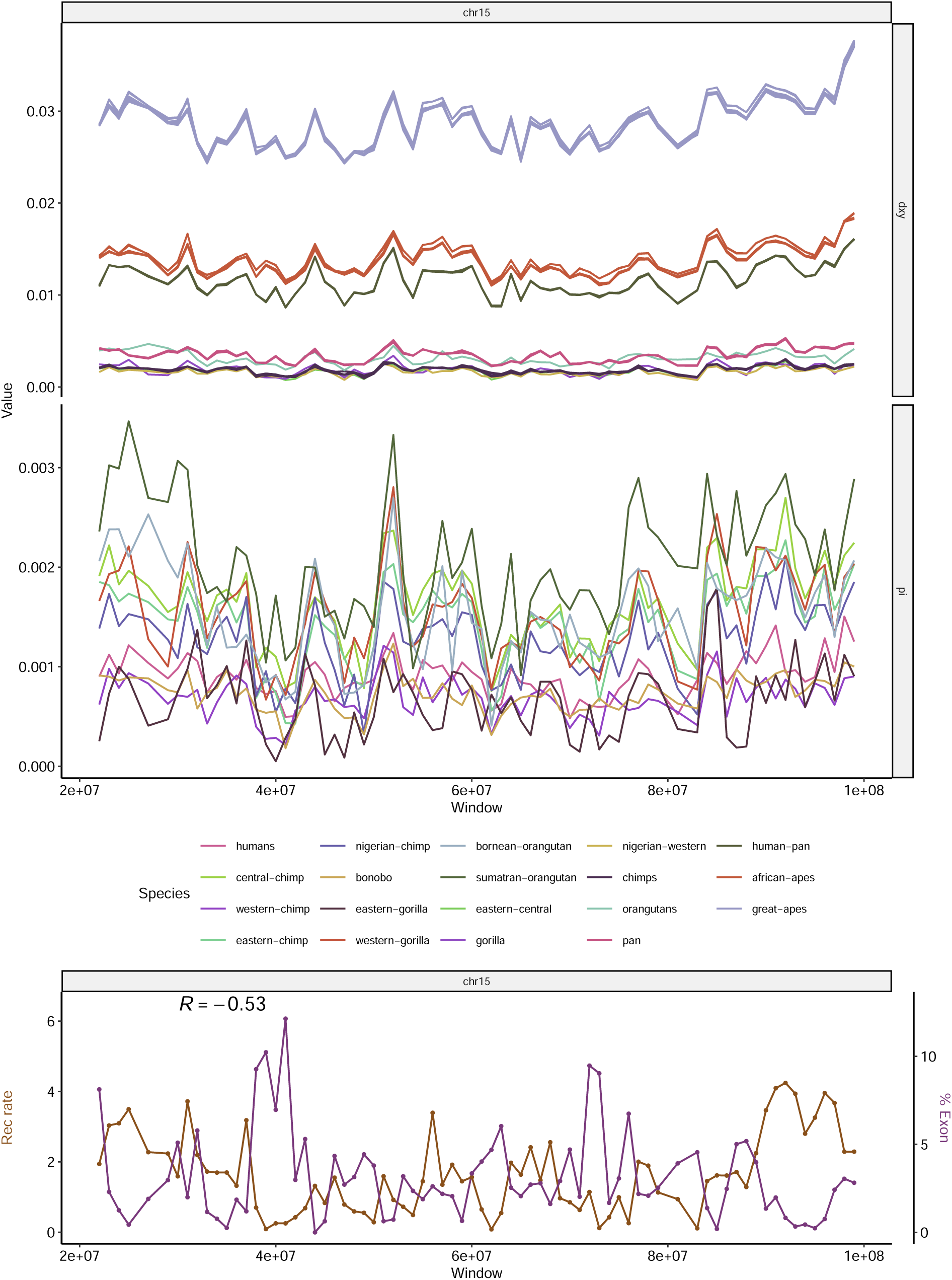
Landscapes of diversity, divergence, exon density and recombination rate across chromosome 15. See Figure 2 for more details.

**Figure S25:**
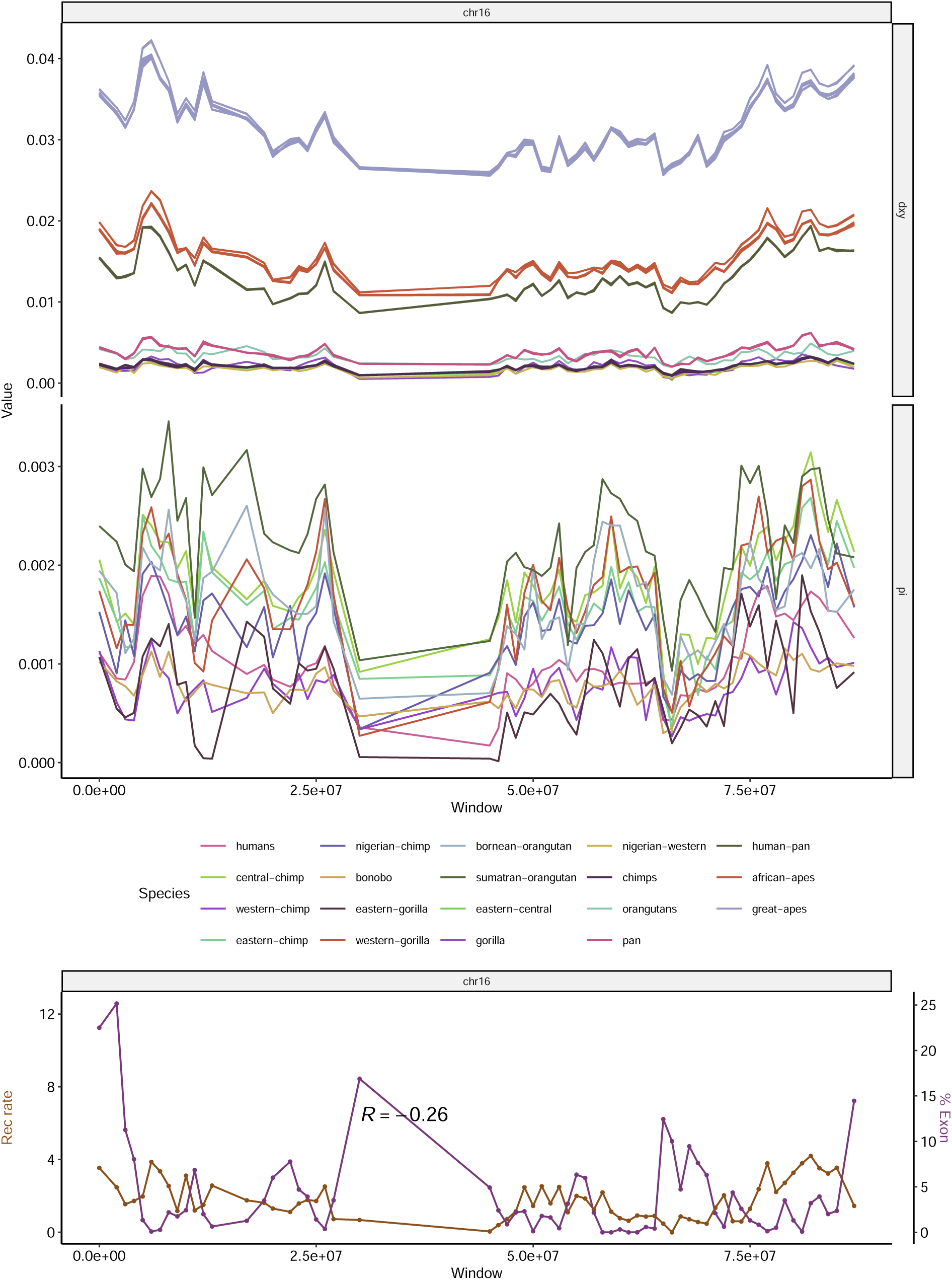
Landscapes of diversity, divergence, exon density and recombination rate across chromosome 16. See Figure 2 for more details.

**Figure S26:**
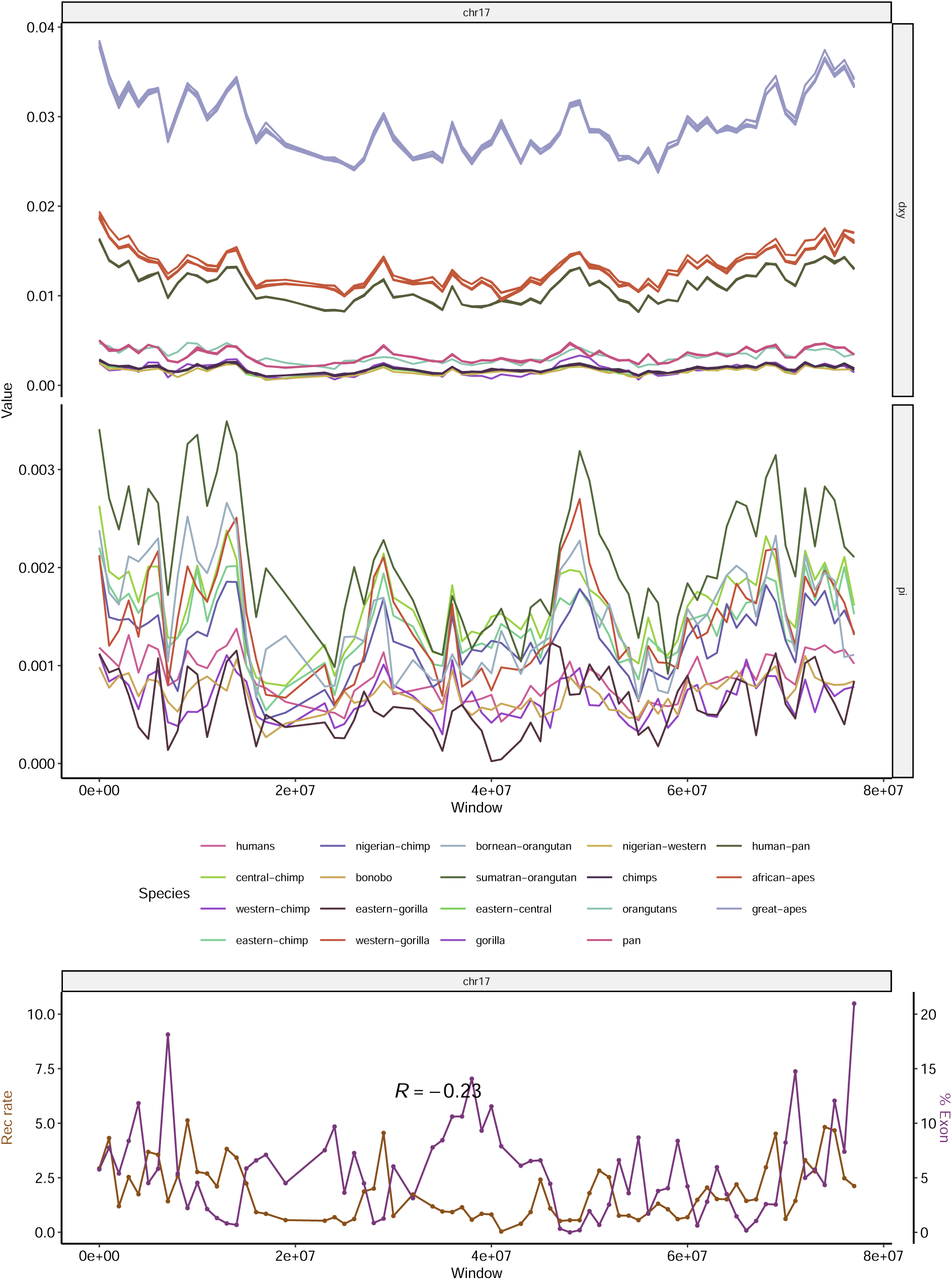
Landscapes of diversity, divergence, exon density and recombination rate across chromosome 17. See Figure 2 for more details.

**Figure S27:**
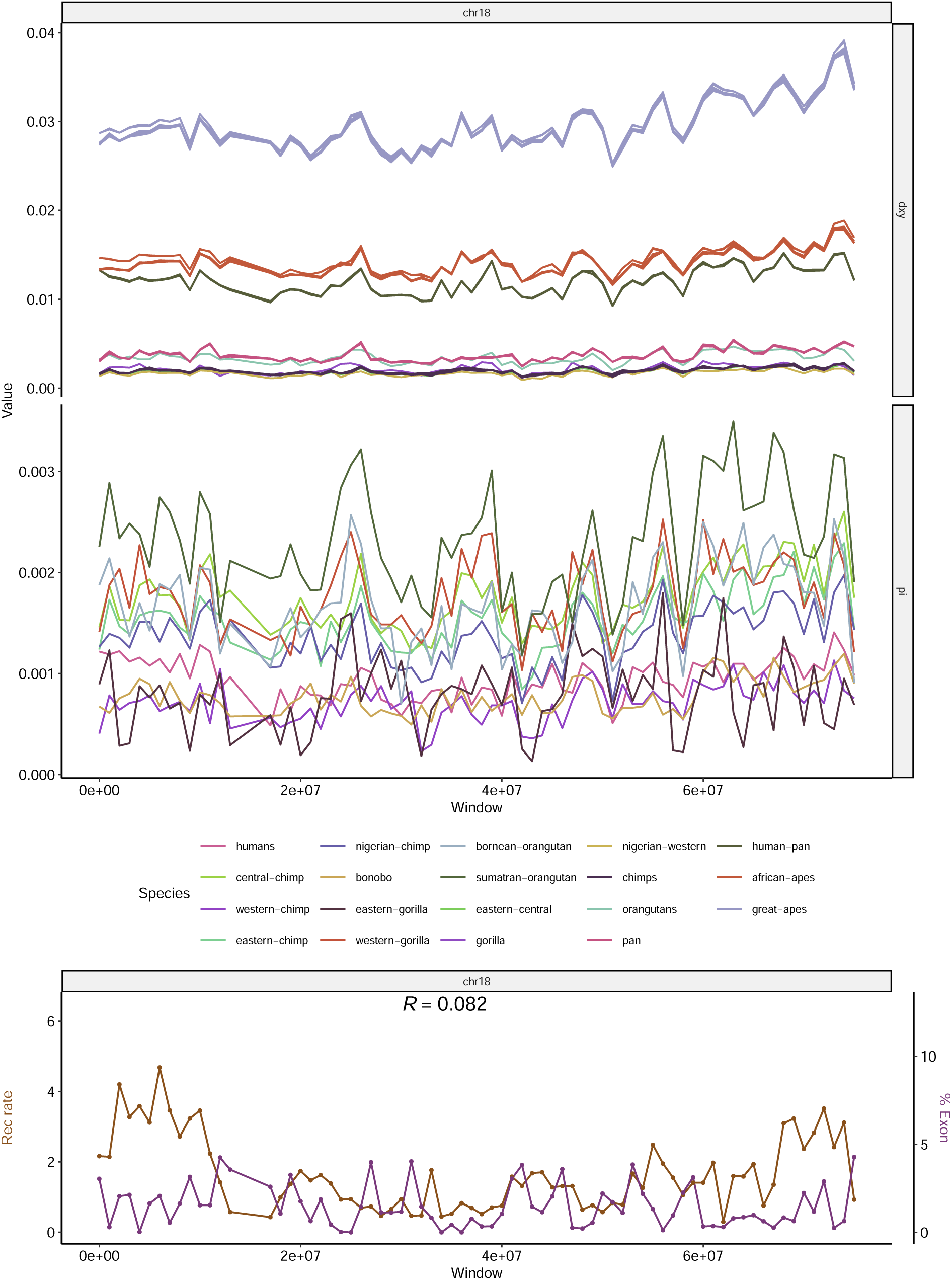
Landscapes of diversity, divergence, exon density and recombination rate across chromosome 18. See Figure 2 for more details.

**Figure S28:**
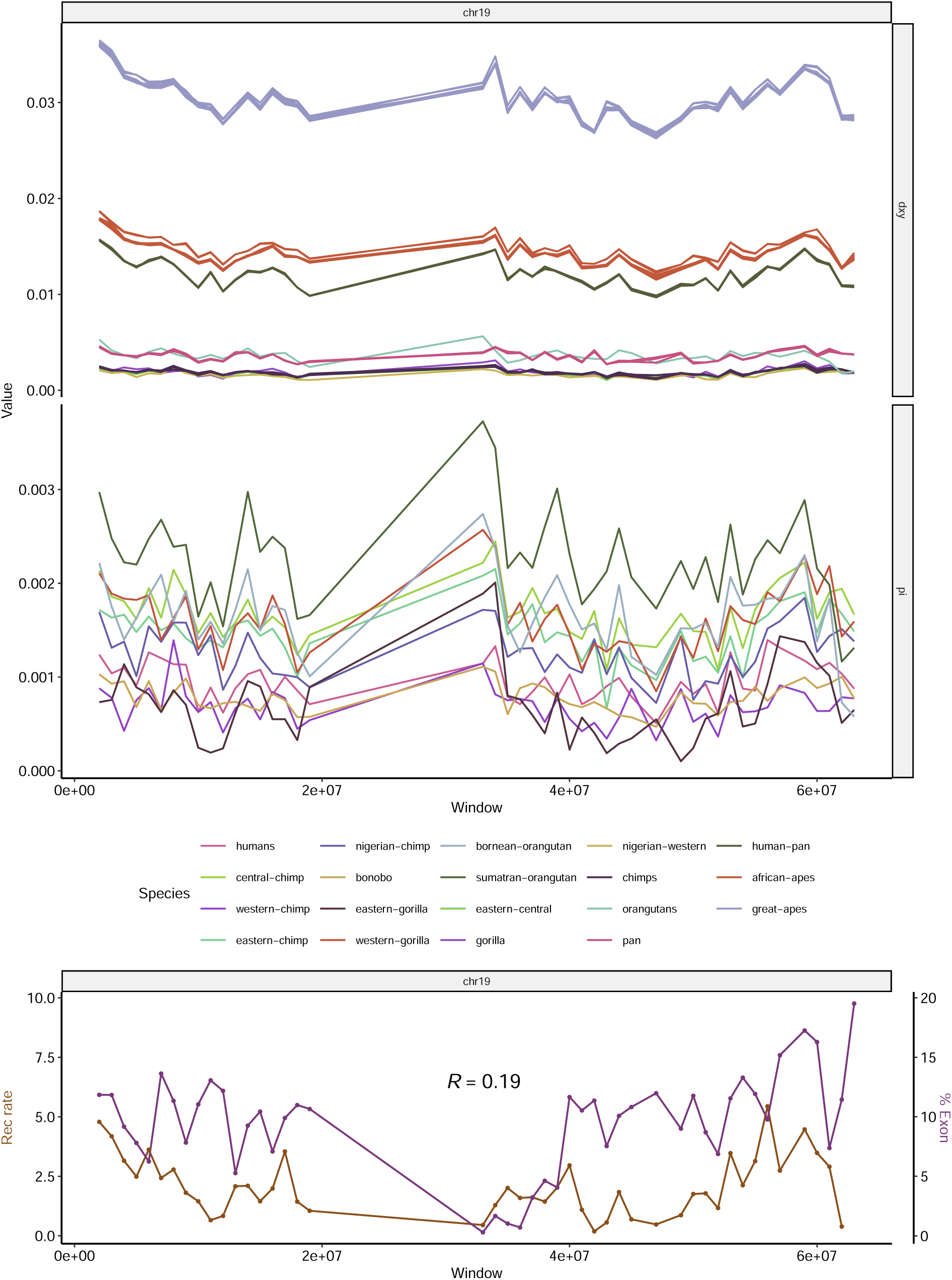
Landscapes of diversity, divergence, exon density and recombination rate across chromosome 19. See Figure 2 for more details.

**Figure S29:**
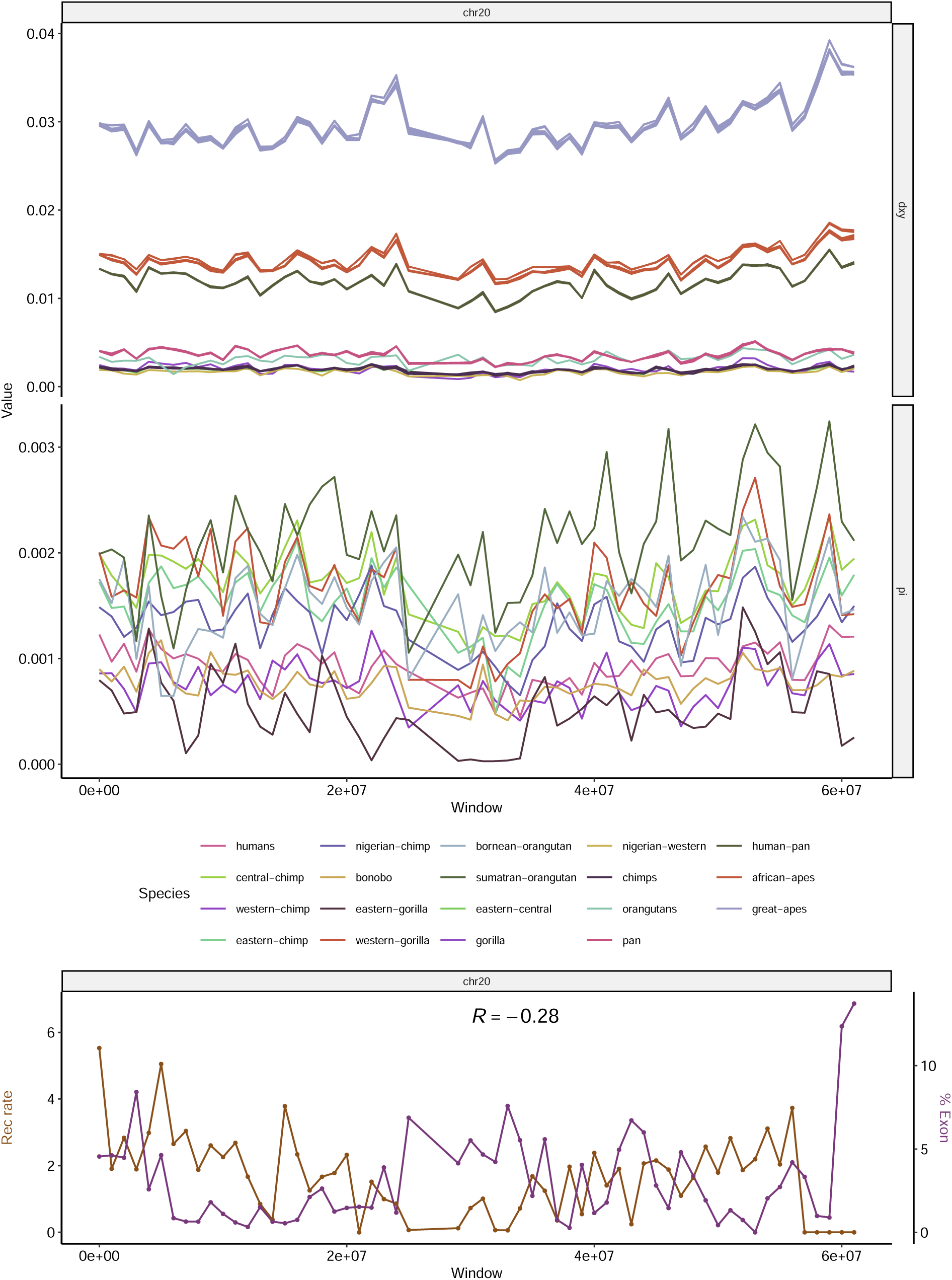
Landscapes of diversity, divergence, exon density and recombination rate across chromosome 20. See Figure 2 for more details.

**Figure S30:**
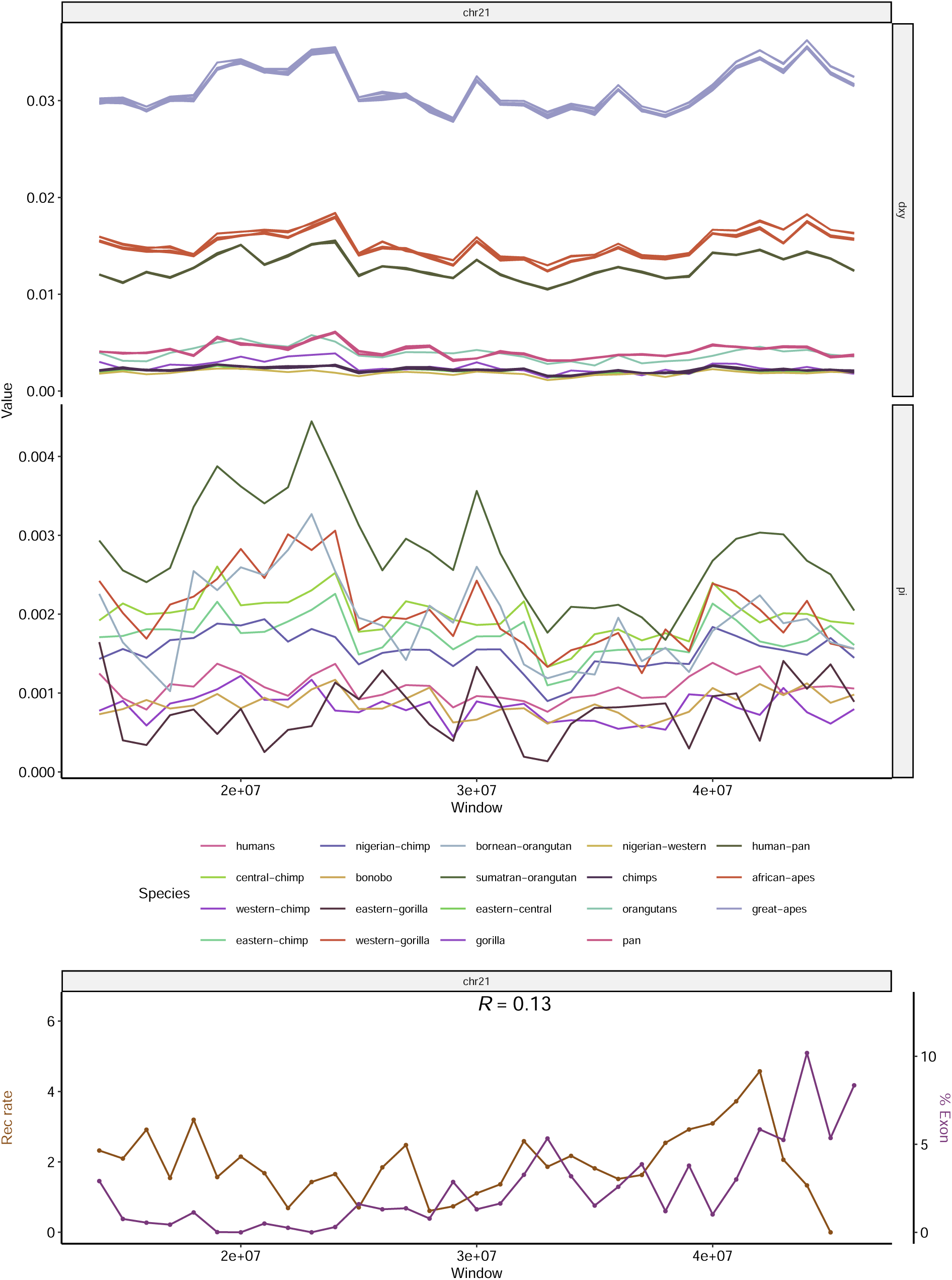
Landscapes of diversity, divergence, exon density and recombination rate across chromosome 21. See Figure 2 for more details.

**Figure S31:**
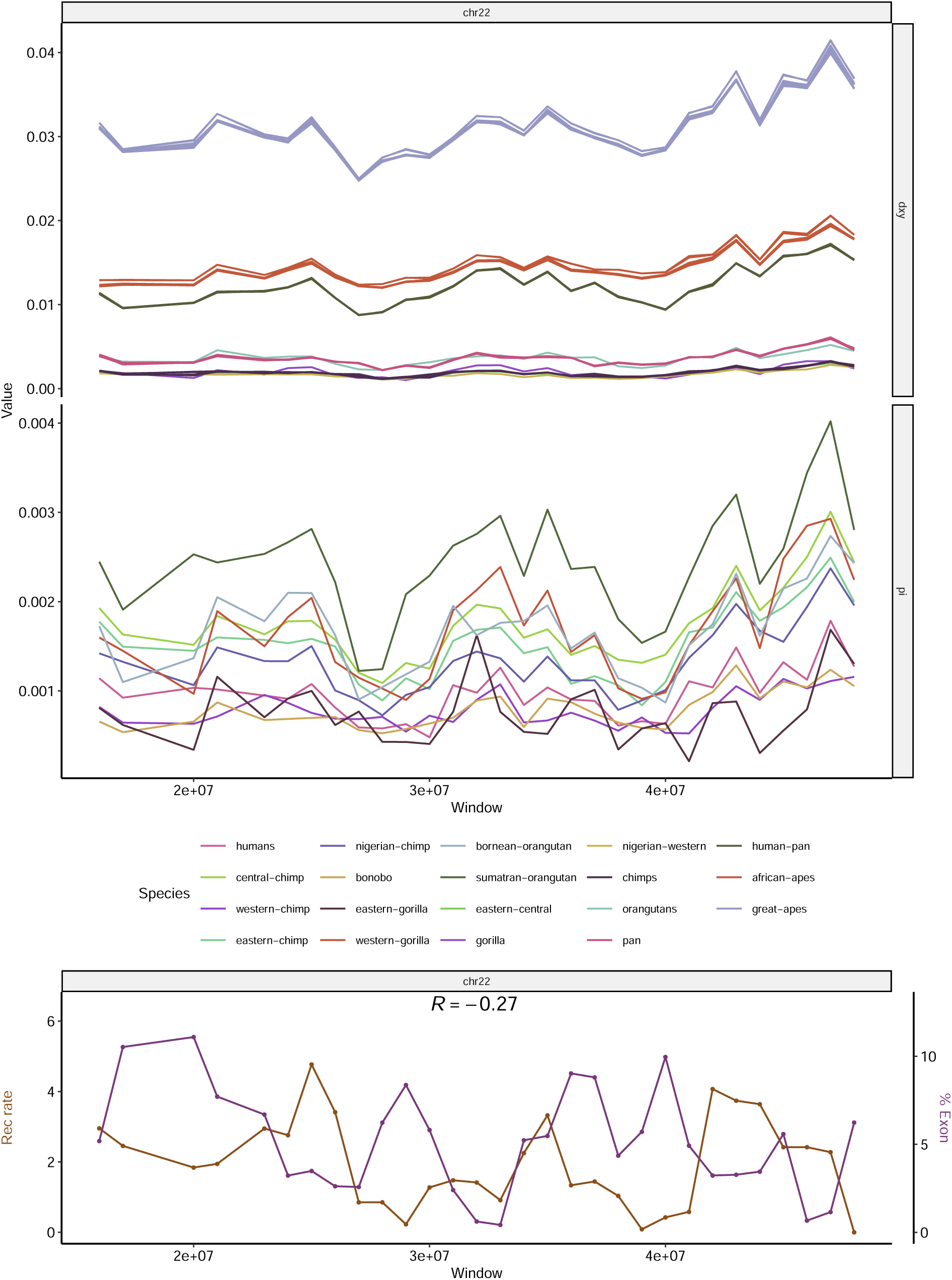
Landscapes of diversity, divergence, exon density and recombination rate across chromosome 22. See Figure 2 for more details.

**Table S1:** Parameter space explored with simulations. *µ_N_* and *µ_P_* are the rates of mutations under negative and positive selection, respectively. *s̄_N_* and *s̄_P_* and the mean fitness effects of negatively and positively selected mutations. *µ_SD_* is the scaled standard deviation of the mutation rate map. See Table 1 and subsection 2.2 for more details.

| $\mu_N$ | $\mu_P$ | $\bar{s}_N$ | $\bar{s}_P$ | Regime | $\mu_{SD}$ |
| --- | --- | --- | --- | --- | --- |
| 0 | 0 | 0 | 0 | Neutral | 0 |
| 0 | 0 | 0 | 0 | Variable $\mu$ | 0.005 |
| 0 | 0 | 0 | 0 | Variable $\mu$ | 0.007 |
| 0 | 0 | 0 | 0 | Variable $\mu$ | 0.011 |
| 0 | 0 | 0 | 0 | Variable $\mu$ | 0.016 |
| 0 | 0 | 0 | 0 | Variable $\mu$ | 0.023 |
| 0 | 0 | 0 | 0 | Variable $\mu$ | 0.033 |
| 0 | 0 | 0 | 0 | Variable $\mu$ | 0.048 |
| 0 | 0 | 0 | 0 | Variable $\mu$ | 0.070 |
| 0 | 0 | 0 | 0 | Variable $\mu$ | 0.103 |
| 0 | 0 | 0 | 0 | Variable $\mu$ | 0.150 |
| 0 | $1 \times 10^{-12}$ | 0 | $1 \times 10^{-2}$ | Beneficial | 0 |
| 0 | $1 \times 10^{-11}$ | 0 | $1 \times 10^{-2}$ | Beneficial | 0 |
| $2 \times 10^{-9}$ | 0 | $-3 \times 10^{-2}$ | 0 | Deleterious | 0 |
| $2 \times 10^{-9}$ | $1 \times 10^{-11}$ | $-3 \times 10^{-2}$ | $1 \times 10^{-2}$ | Both | 0 |
| $2 \times 10^{-9}$ | 0 | $-1.5 \times 10^{-2}$ | 0 | Deleterious | 0 |
| $2 \times 10^{-9}$ | $1 \times 10^{-11}$ | $-1.5 \times 10^{-2}$ | $1 \times 10^{-2}$ | Both | 0 |
| $2 \times 10^{-9}$ | 0 | $-1 \times 10^{-2}$ | 0 | Deleterious | 0 |
| $2 \times 10^{-9}$ | $1 \times 10^{-12}$ | $-1 \times 10^{-2}$ | $5 \times 10^{-3}$ | Both | 0 |
| $2 \times 10^{-9}$ | $1 \times 10^{-12}$ | $-1 \times 10^{-2}$ | $1 \times 10^{-2}$ | Both | 0 |
| $2 \times 10^{-9}$ | 0 | $-3 \times 10^{-3}$ | 0 | Deleterious | 0 |
| $2 \times 10^{-9}$ | $1 \times 10^{-12}$ | $-3 \times 10^{-3}$ | $5 \times 10^{-3}$ | Both | 0 |
| $2 \times 10^{-9}$ | $1 \times 10^{-12}$ | $-3 \times 10^{-3}$ | $1 \times 10^{-2}$ | Both | 0 |
| $2 \times 10^{-9}$ | 0 | $-1 \times 10^{-3}$ | 0 | Deleterious | 0 |
| $2 \times 10^{-9}$ | $1 \times 10^{-11}$ | $-1 \times 10^{-3}$ | $1 \times 10^{-2}$ | Both | 0 |
| $6 \times 10^{-9}$ | 0 | $-3 \times 10^{-2}$ | 0 | Deleterious | 0 |
| $6 \times 10^{-9}$ | $1 \times 10^{-11}$ | $-3 \times 10^{-2}$ | $1 \times 10^{-2}$ | Both | 0 |
| $6 \times 10^{-9}$ | 0 | $-1.5 \times 10^{-2}$ | 0 | Deleterious | 0 |
| $6 \times 10^{-9}$ | $1 \times 10^{-11}$ | $-1.5 \times 10^{-2}$ | $1 \times 10^{-2}$ | Both | 0 |
| $1 \times 10^{-8}$ | $1 \times 10^{-11}$ | $-1 \times 10^{-3}$ | $1 \times 10^{-2}$ | Both | 0 |
| $1.2 \times 10^{-8}$ | 0 | $-3 \times 10^{-2}$ | 0 | Deleterious | 0 |
| $1.2 \times 10^{-8}$ | $1 \times 10^{-12}$ | $-3 \times 10^{-2}$ | $1 \times 10^{-2}$ | Both | 0 |
| $1.2 \times 10^{-8}$ | $1 \times 10^{-12}$ | $-3 \times 10^{-2}$ | $1 \times 10^{-2}$ | Both | 0.005 |
| $1.2 \times 10^{-8}$ | $1 \times 10^{-12}$ | $-3 \times 10^{-2}$ | $1 \times 10^{-2}$ | Both | 0.007 |
| $1.2 \times 10^{-8}$ | $1 \times 10^{-12}$ | $-3 \times 10^{-2}$ | $1 \times 10^{-2}$ | Both | 0.011 |
| $1.2 \times 10^{-8}$ | $1 \times 10^{-12}$ | $-3 \times 10^{-2}$ | $1 \times 10^{-2}$ | Both | 0.016 |
| $1.2 \times 10^{-8}$ | $1 \times 10^{-12}$ | $-3 \times 10^{-2}$ | $1 \times 10^{-2}$ | Both | 0.023 |
| $1.2 \times 10^{-8}$ | $1 \times 10^{-12}$ | $-3 \times 10^{-2}$ | $1 \times 10^{-2}$ | Both | 0.033 |
| $1.2 \times 10^{-8}$ | $1 \times 10^{-12}$ | $-3 \times 10^{-2}$ | $1 \times 10^{-2}$ | Both | 0.048 |
| $1.2 \times 10^{-8}$ | $1 \times 10^{-12}$ | $-3 \times 10^{-2}$ | $1 \times 10^{-2}$ | Both | 0.070 |
| $1.2 \times 10^{-8}$ | $1 \times 10^{-12}$ | $-3 \times 10^{-2}$ | $1 \times 10^{-2}$ | Both | 0.103 |
| $1.2 \times 10^{-8}$ | $1 \times 10^{-12}$ | $-3 \times 10^{-2}$ | $1 \times 10^{-2}$ | Both | 0.150 |
| $1.2 \times 10^{-8}$ | $1 \times 10^{-11}$ | $-3 \times 10^{-2}$ | $1 \times 10^{-2}$ | Both | 0 |
| $1.4 \times 10^{-8}$ | 0 | $-3 \times 10^{-2}$ | 0 | Deleterious | 0 |
| $1.4 \times 10^{-8}$ | 0 | $-3 \times 10^{-2}$ | 0 | Deleterious | 0.005 |
| $1.4 \times 10^{-8}$ | 0 | $-3 \times 10^{-2}$ | 0 | Deleterious | 0.007 |
| $1.4 \times 10^{-8}$ | 0 | $-3 \times 10^{-2}$ | 0 | Deleterious | 0.011 |
| $1.4 \times 10^{-8}$ | 0 | $-3 \times 10^{-2}$ | 0 | Deleterious | 0.016 |
| $1.4 \times 10^{-8}$ | 0 | $-3 \times 10^{-2}$ | 0 | Deleterious | 0.023 |
| $1.4 \times 10^{-8}$ | 0 | $-3 \times 10^{-2}$ | 0 | Deleterious | 0.033 |
| $1.4 \times 10^{-8}$ | 0 | $-3 \times 10^{-2}$ | 0 | Deleterious | 0.048 |
| $1.4 \times 10^{-8}$ | 0 | $-3 \times 10^{-2}$ | 0 | Deleterious | 0.070 |
| $1.4 \times 10^{-8}$ | 0 | $-3 \times 10^{-2}$ | 0 | Deleterious | 0.103 |
| $1.4 \times 10^{-8}$ | 0 | $-3 \times 10^{-2}$ | 0 | Deleterious | 0.150 |
| $1.4 \times 10^{-8}$ | $1 \times 10^{-12}$ | $-3 \times 10^{-2}$ | $1 \times 10^{-2}$ | Both | 0 |
| $1.4 \times 10^{-8}$ | $1 \times 10^{-11}$ | $-3 \times 10^{-2}$ | $1 \times 10^{-2}$ | Both | 0 |

## Notes

### Competing Interest Statement

The authors have declared no competing interest.

### Summary of Updates

New simulations with both mutation rate variation and selection. Text changed for clarity and to improve conclusions. Minor changes throughout the text.

## References

Agarwal, I., & Przeworski, M. (2021). Mutation saturation for fitness effects at human CpG sites (J. Ross-Ibarra & P. J. Wittkopp, Eds.). eLife, 10, e71513. 10.7554/eLife.71513

Andolfatto, P. (2001). Adaptive hitchhiking effects on genome variability. Current Opinion in Genetics & Development, 11 (6), 635–641. 10.1016/S0959-437X(00)00246-X

Andolfatto, P. (2007). Hitchhiking effects of recurrent beneficial amino acid substitutions in the drosophila melanogaster genome. Genome Research, 17 (12), 1755–1762. 10.1101/gr.6691007

Battey, C. J. (2020). Evidence of linked selection on the z chromosome of hybridizing hummingbirds*. Evolution, 74 (4), 725–739. 10.1111/evo.13888

Begun, D. J., & Aquadro, C. F. (1992). Levels of naturally occurring DNA polymorphism correlate with recombination rates in d. melanogaster. Nature, 356 (6369), 519–520. 10.1038/ 356519a0

Begun, D. J., Holloway, A. K., Stevens, K., Hillier, L. W., Poh, Y.-P., Hahn, M. W., Nista, P. M., Jones, C. D., Kern, A. D., Dewey, C. N., Pachter, L., Myers, E., & Langley, C. H. (2007). Population genomics: Whole-genome analysis of polymorphism and divergence in drosophila simulans. PLOS Biology, 5 (11), e310. 10.1371/journal.pbio.0050310

Beissinger, T. M., Wang, L., Crosby, K., Durvasula, A., Hufford, M. B., & Ross-Ibarra, J. (2016). Recent demography drives changes in linked selection across the maize genome. Nature Plants, 2 (7), 1–7. 10.1038/nplants.2016.84

Bejerano, G., Pheasant, M., Makunin, I., Stephen, S., Kent, W. J., Mattick, J. S., & Haussler, D. (2004). Ultraconserved elements in the human genome. *Science (New York*, N.Y.), 304 (5675), 1321–1325. 10.1126/science.1098119

Birky, C. W., & Walsh, J. B. (1988). Effects of linkage on rates of molecular evolution. Proceedings of the National Academy of Sciences, 85 (17), 6414–6418. 10.1073/pnas.85.17.6414

Boyko, A. R., Williamson, S. H., Indap, A. R., Degenhardt, J. D., Hernandez, R. D., Lohmueller, K. E., Adams, M. D., Schmidt, S., Sninsky, J. J., Sunyaev, S. R., White, T. J., Nielsen, R., Clark, A. G., & Bustamante, C. D. (2008). Assessing the evolutionary impact of amino acid mutations in the human genome. PLoS genetics, 4 (5), e1000083. 10.1371/journal.pgen.1000083

Burri, R. (2017). Interpreting differentiation landscapes in the light of long-term linked selection. Evolution Letters, 1 (3), 118–131. 10.1002/evl3.14

Burri, R., Nater, A., Kawakami, T., Mugal, C. F., Olason, P. I., Smeds, L., Suh, A., Dutoit, L., Burěs, S., Garamszegi, L. Z., Hogner, S., Moreno, J., Qvarnström, A., Ružić, M., Sæther, S.-A., Sætre, G.-P., Török, J., & Ellegren, H. (2015). Linked selection and recombination rate variation drive the evolution of the genomic landscape of differentiation across the speciation continuum of ficedula flycatchers. Genome Research, 25 (11), 1656–1665. 10.1101/gr.196485.115

Cai, J. J., Macpherson, J. M., Sella, G., & Petrov, D. A. (2009). Pervasive hitchhiking at coding and regulatory sites in humans. PLOS Genetics, 5 (1), e1000336. 10.1371/journal.pgen1000336.

Castellano, D., Eyre-Walker, A., & Munch, K. (2020). Impact of mutation rate and selection at linked sites on DNA variation across the genomes of humans and other homininae. Genome Biology and Evolution, 12 (1), 3550–3561. 10.1093/gbe/evz215

Castellano, D., Macìa, M. C., Tataru, P., Bataillon, T., & Munch, K. (2019). Comparison of the full distribution of fitness effects of new amino acid mutations across great apes. Genetics, 213 (3), 953–966. 10.1534/genetics.119.302494

Charlesworth, B., Morgan, M. T., & Charlesworth, D. (1993). The effect of deleterious mutations on neutral molecular variation. Genetics, 134 (4), 1289–1303.

Chen, J.-M., Cooper, D. N., Chuzhanova, N., Férec, C., & Patrinos, G. P. (2007). Gene conversion: Mechanisms, evolution and human disease. Nature Reviews Genetics, 8 (10), 762–775. 10.1038/nrg2193

Corbett-Detig, R. B., Hartl, D. L., & Sackton, T. B. (2015). Natural selection constrains neutral diversity across a wide range of species. PLOS Biology, 13 (4), e1002112. 10.1371/journalpbio.1002112.

Cruickshank, T. E., & Hahn, M. W. (2014). Reanalysis suggests that genomic islands of speciation are due to reduced diversity, not reduced gene flow. Molecular Ecology, 23 (13), 3133–3157. 10.1111/mec.12796

Delmore, K. E., Lugo Ramos, J. S., van Doren, B. M., Lundberg, M., Bensch, S., Irwin, D. E., & Liedvogel, M. (2018). Comparative analysis examining patterns of genomic differentiation across multiple episodes of population divergence in birds. Evolution Letters, 2 (2), 76–87. 10.1002/evl3.46

Ellegren, H., Smeds, L., Burri, R., Olason, P. I., Backström, N., Kawakami, T., Künstner, A., Mäkinen, H., Nadachowska-Brzyska, K., Qvarnström, A., Uebbing, S., & Wolf, J. B. W. (2012). The genomic landscape of species divergence in ficedula flycatchers. Nature, 491 (7426), 756–760. 10.1038/nature11584

Enard, D., Messer, P. W., & Petrov, D. A. (2014). Genome-wide signals of positive selection in human evolution. Genome Research, 24 (6), 885–895. 10.1101/gr.164822.113

Galtier, N. (2016). Adaptive protein evolution in animals and the effective population size hypothesis. PLOS Genetics, 12 (1), e1005774. 10.1371/journal.pgen.1005774

Galtier, N., Duret, L., Gĺemin, S., & Ranwez, V. (2009). GC-biased gene conversion promotes the fixation of deleterious amino acid changes in primates. Trends in genetics: TIG, 25 (1), 1–5. 10.1016/j.tig.2008.10.011

Gĺemin, S., Arndt, P. F., Messer, P. W., Petrov, D., Galtier, N., & Duret, L. (2015). Quantification of GC-biased gene conversion in the human genome. Genome Research, 25 (8), 1215–1228. 10.1101/gr.185488.114

Gower, G., Ragsdale, A. P., Bisschop, G., Gutenkunst, R. N., Hartfield, M., Noskova, E., Schiffels, S., Struck, T. J., Kelleher, J., & Thornton, K. R. (2022). Demes: A standard format for demographic models. Genetics, 222 (3), iyac131. 10.1093/genetics/iyac131

Haller, B. C., Galloway, J., Kelleher, J., Messer, P. W., & Ralph, P. L. (2019). Tree-sequence recording in SLiM opens new horizons for forward-time simulation of whole genomes. Molecular Ecology Resources, 19 (2), 552–566. 10.1111/1755-0998.12968

Haller, B. C., & Messer, P. W. (2019). SLiM 3: Forward genetic simulations beyond the wright-fisher model. Molecular Biology and Evolution, 36 (3), 632–637. 10.1093/molbev/msy228

Halligan, D. L., Kousathanas, A., Ness, R. W., Harr, B., Ëory, L., Keane, T. M., Adams, D. J., & Keightley, P. D. (2013). Contributions of protein-coding and regulatory change to adaptive molecular evolution in murid rodents. PLoS Genetics, 9 (12). 10.1371/journal.pgen.1003995

Harr, B. (2006). Genomic islands of differentiation between house mouse subspecies. Genome Research, 16 (6), 730–737. 10.1101/gr.5045006

Hernandez, R. D., Kelley, J. L., Elyashiv, E., Melton, S. C., Auton, A., McVean, G., 1000 Genomes Project, Sella, G., & Przeworski, M. (2011). Classic selective sweeps were rare in recent human evolution. Science (New York, N.Y.), 331 (6019), 920–924. 10.1126/science.1198878

Hodgkinson, A., & Eyre-Walker, A. (2011). Variation in the mutation rate across mammalian genomes. Nature Reviews Genetics, 12 (11), 756–766. 10.1038/nrg3098

Huber, C. D., Kim, B. Y., Marsden, C. D., & Lohmueller, K. E. (2017). Determining the factors driving selective effects of new nonsynonymous mutations. Proceedings of the National Academy of Sciences, 114 (17), 4465–4470. 10.1073/pnas.1619508114

Hudson, R. R. (1983). Testing the constant-rate neutral allele model with protein sequence data. Evolution, 37 (1), 203–217. 10.2307/2408186

Hudson, R. R., Kreitman, M., & Aguadé, M. (1987). A test of neutral molecular evolution based on nucleotide data. Genetics, 116 (1), 153–159. 10.1093/genetics/116.1.153

Ingvarsson, P. K. (2010). Natural selection on synonymous and nonsynonymous mutations shapes patterns of polymorphism in populus tremula. Molecular Biology and Evolution, 27 (3), 650–660. https://doi.org/10.1093/molbev/msp255

Irwin, D. E., Alcaide, M., Delmore, K. E., Irwin, J. H., & Owens, G. L. (2016). Recurrent selection explains parallel evolution of genomic regions of high relative but low absolute differentiation in a ring species. Molecular Ecology, 25 (18), 4488–4507. 10.1111/mec.13792

Jauch, A., Wienberg, J., Stanyon, R., Arnold, N., Tofanelli, S., Ishida, T., & Cremer, T. (1992). Reconstruction of genomic rearrangements in great apes and gibbons by chromosome painting. Proceedings of the National Academy of Sciences of the United States of America, 89 (18), 8611–8615. Retrieved February 21, 2020, from https://www.ncbi.nlm.nih.gov/pmc/articles/PMC49970/

Jensen, J. D., Kim, Y., DuMont, V. B., Aquadro, C. F., & Bustamante, C. D. (2005). Distinguishing between selective sweeps and demography using DNA polymorphism data. Genetics, 170 (3), 1401–1410. 10.1534/genetics.104.038224

Kaplan, N. L., Hudson, R. R., & Langley, C. H. (1989). The “hitchhiking effect” revisited. Genetics, 123 (4), 887–899.

Katzman, S., Kern, A. D., Bejerano, G., Fewell, G., Fulton, L., Wilson, R. K., Salama, S. R., & Haussler, D. (2007). Human genome ultraconserved elements are ultraselected. Science, 317 (5840), 915–915. 10.1126/science.1142430

Katzman, S., Kern, A. D., Pollard, K. S., Salama, S. R., & Haussler, D. (2010). GC-biased evolution near human accelerated regions (J. Zhang, Ed.). PLoS Genetics, 6 (5), e1000960. 10.1371/journal.pgen.1000960

Kelleher, J., Etheridge, A. M., & McVean, G. (2016). Efficient coalescent simulation and genealogical analysis for large sample sizes. PLoS computational biology, 12 (5), e1004842. 10.1371/ journal.pcbi.1004842

Kelleher, J., Thornton, K. R., Ashander, J., & Ralph, P. L. (2018). Efficient pedigree recording for fast population genetics simulation. PLoS computational biology, 14 (11), e1006581. 10.1371/journal.pcbi.1006581

Kern, A. D., & Hahn, M. W. (2018). The neutral theory in light of natural selection. Molecular Biology and Evolution, 35 (6), 1366–1371. 10.1093/molbev/msy092

Kern, A. D., Jones, C. D., & Begun, D. J. (2002). Genomic effects of nucleotide substitutions in drosophila simulans. Genetics, 162 (4), 1753–1761. Retrieved February 9, 2020, from https://www.genetics. org/content/162/4/1753

Kim, B. Y., Huber, C. D., & Lohmueller, K. E. (2017). Inference of the distribution of selection coefficients for new nonsynonymous mutations using large samples. Genetics, 206 (1), 345–361.

Kim, Y., & Stephan, W. (2000). Joint effects of genetic hitchhiking and background selection on neutral variation. Genetics, 155 (3), 1415–1427. Retrieved February 8, 2020, from https://www.ncbi.nlm. nih.gov/pmc/articles/PMC1461159/

Kong, A., Gudbjartsson, D. F., Sainz, J., Jonsdottir, G. M., Gudjonsson, S. A., Richardsson, B., Sigurdardottir, S., Barnard, J., Hallbeck, B., Masson, G., Shlien, A., Palsson, S. T., Frigge, M. L., Thorgeirsson, T. E., Gulcher, J. R., & Stefansson, K. (2002). A high-resolution recombination map of the human genome. Nature Genetics, 31 (3), 241–247. 10.1038/ng917

Kong, A., Thorleifsson, G., Gudbjartsson, D. F., Masson, G., Sigurdsson, A., Jonasdottir, A., Walters, G. B., Jonasdottir, A., Gylfason, A., Kristinsson, K. T., Gudjonsson, S. A., Frigge, M. L., Helgason, A., Thorsteinsdottir, U., & Stefansson, K. (2010). Fine-scale recombination rate differences between sexes, populations and individuals. Nature, 467 (7319), 1099–1103. 10.1038/ nature09525

Kronenberg, Z. N., Fiddes, I. T., Gordon, D., Murali, S., Cantsilieris, S., Meyerson, O. S., Underwood, J. G., Nelson, B. J., Chaisson, M. J. P., Dougherty, M. L., Munson, K. M., Hastie, A. R., Diekhans, M., Hormozdiari, F., Lorusso, N., Hoekzema, K., Qiu, R., Clark, K., Raja, A.,... Eichler, E.E. (2018). High-resolution comparative analysis of great ape genomes. Science, 360 (6393). 10.1126/science.aar6343

Laval, G., Patin, E., Boutillier, P., & Quintana-Murci, L. (2021). Sporadic occurrence of recent selective sweeps from standing variation in humans as revealed by an approximate bayesian computation approach. Genetics, 219 (4), iyab161. 10.1093/genetics/iyab161

Lewontin, R. C. (1974). The genetic basis of evolutionary change (Vol. 560). Columbia University Press New York.

Lohmueller, K. E., Albrechtsen, A., Li, Y., Kim, S. Y., Korneliussen, T., Vinckenbosch, N., Tian, G., Huerta-Sanchez, E., Feder, A. F., Grarup, N., Jørgensen, T., Jiang, T., Witte, D. R., Sandbæk, A., Hellmann, I., Lauritzen, T., Hansen, T., Pedersen, O., Wang, J., & Nielsen, R. (2011). Natural selection affects multiple aspects of genetic variation at putatively neutral sites across the human genome. PLOS Genetics, 7 (10), e1002326. 10.1371/journal.pgen.1002326

Macpherson, J. M., Sella, G., Davis, J. C., & Petrov, D. A. (2007). Genomewide spatial correspondence between nonsynonymous divergence and neutral polymorphism reveals extensive adaptation in drosophila. Genetics, 177 (4), 2083–2099. 10.1534/genetics.107.080226

Maynard Smith, J., & Haigh, J. (1974). The hitch-hiking effect of a favourable gene. Genetical Research, 23 (1), 23–35.

McDonald, J. H., & Kreitman, M. (1991). Adaptive protein evolution at the adh locus in drosophila. Nature, 351 (6328), 652–654. 10.1038/351652a0

Miles, A., Bot, P. I., Rodrigues, M. F., Ralph, P., Harding, N., Pisupati, R., & Rae, S. (2020). Cggh/scikitallel: V1. 3.2. *Zenodo*.

Murphy, D. A., Elyashiv, E., Amster, G., & Sella, G. (2022). Broad-scale variation in human genetic diversity levels is predicted by purifying selection on coding and non-coding elements (M. Nordborg, Ed.). eLife, 11, e76065. 10.7554/eLife.76065

Nachman, M. W., & Crowell, S. L. (2000). Estimate of the mutation rate per nucleotide in humans. Genetics, 156 (1), 297–304. 10.1093/genetics/156.1.297

Nam, K., Munch, K., Mailund, T., Nater, A., Greminger, M. P., Krützen, M., Marqùes-Bonet, T., & Schierup, M. H. (2017). Evidence that the rate of strong selective sweeps increases with population size in the great apes. Proceedings of the National Academy of Sciences, 114 (7), 1613–1618. 10.1073/pnas.1605660114

Nielsen, R., Williamson, S., Kim, Y., Hubisz, M. J., Clark, A. G., & Bustamante, C. (2005). Genomic scans for selective sweeps using SNP data [Company: Cold Spring Harbor Laboratory Press Distributor: Cold Spring Harbor Laboratory Press Institution: Cold Spring Harbor Laboratory Press Label: Cold Spring Harbor Laboratory Press Publisher: Cold Spring Harbor Lab]. Genome Research, 15 (11), 1566–1575. 10.1101/gr.4252305

Orr, H. A. (2003). The distribution of fitness effects among beneficial mutations. Genetics, 163 (4), 1519–1526. Retrieved September 27, 2023, from https://www.ncbi.nlm.nih.gov/pmc/articles/PMC1462510/

Phung, T. N., Huber, C. D., & Lohmueller, K. E. (2016). Determining the effect of natural selection on linked neutral divergence across species. PLOS Genetics, 12 (8), e1006199. 10.1371/journal.pgen.1006199

Pouyet, F., Aeschbacher, S., Thíery, A., & Excoffier, L. (2018). Background selection and biased gene conversion affect more than 95% of the human genome and bias demographic inferences (K. Veeramah, P. J. Wittkopp, & I. Gronau, Eds.). eLife, 7, e36317. 10.7554/eLife.36317

Prado-Martinez, J., Sudmant, P. H., Kidd, J. M., Li, H., Kelley, J. L., Lorente-Galdos, B., Veeramah, K. R., Woerner, A. E., O’Connor, T. D., Santpere, G., Cagan, A., Theunert, C., Casals, F., Laayouni, H., Munch, K., Hobolth, A., Halager, A. E., Malig, M., Hernandez-Rodriguez, J.,... Marques-Bonet, T. (2013). Great ape genetic diversity and population history. Nature, 499 (7459), 471–475. 10.1038/nature12228

Przeworski, M. (2002). The signature of positive selection at randomly chosen loci. Genetics, 160 (3), 1179– 1189. 10.1093/genetics/160.3.1179

Ralph, P., Thornton, K., & Kelleher, J. (2020). Efficiently summarizing relationships in large samples: A general duality between statistics of genealogies and genomes. Genetics, 215 (3), 779–797. 10.1534/genetics.120.303253

Rodrigues, M. F., & Ralph, P. L. (2021). Vignette: Parallelizing SLiM simulations in a phylogenetic tree — PySLiM manual. Retrieved September 24, 2021, from https://tskit.dev/pyslim/docs/latest/vignetteparallelphylo.html

Sattath, S., Elyashiv, E., Kolodny, O., Rinott, Y., & Sella, G. (2011). Pervasive adaptive protein evolution apparent in diversity patterns around amino acid substitutions in drosophila simulans. PLoS genetics, 7 (2), e1001302. 10.1371/journal.pgen.1001302

Scally, A., & Durbin, R. (2012). Revising the human mutation rate: Implications for understanding human evolution [Number: 10 Publisher: Nature Publishing Group]. Nature Reviews Genetics, 13 (10), 745–753. 10.1038/nrg3295

Schrider, D. R. (2020). Background selection does not mimic the patterns of genetic diversity produced by selective sweeps. Genetics, 216 (2), 499–519. 10.1534/genetics.120.303469

Schrider, D. R., & Kern, A. D. (2016). S/HIC: Robust identification of soft and hard sweeps using machine learning. PLOS Genetics, 12 (3), e1005928. 10.1371/journal.pgen.1005928

Schrider, D. R., & Kern, A. D. (2017). Soft sweeps are the dominant mode of adaptation in the human genome. Molecular Biology and Evolution, 34 (8), 1863–1877. 10.1093/molbev/msx154

Siepel, A., Bejerano, G., Pedersen, J. S., Hinrichs, A. S., Hou, M., Rosenbloom, K., Clawson, H., Spieth, J., Hillier, L. W., Richards, S., Weinstock, G. M., Wilson, R. K., Gibbs, R. A., Kent, W. J., Miller, W., & Haussler, D. (2005). Evolutionarily conserved elements in vertebrate, insect, worm, and yeast genomes. Genome Research, 15 (8), 1034–1050. 10.1101/gr.3715005

Simonsen, K. L., Churchill, G. A., & Aquadro, C. F. (1995). Properties of statistical tests of neutrality for DNA polymorphism data. Genetics, 141 (1), 413–429. 10.1093/genetics/141.1.413

Slotte, T. (2014). The impact of linked selection on plant genomic variation. Briefings in Functional Genomics, 13 (4), 268–275. 10.1093/bfgp/elu009

Smith, N., & Eyre-Walker, A. (2002). Adaptive protein evolution in drosophila. Nature, 415 (6875), 1022– 1024. 10.1038/4151022a

Smith, T. C. A., Arndt, P. F., & Eyre-Walker, A. (2018). Large scale variation in the rate of germ-line de novo mutation, base composition, divergence and diversity in humans. PLOS Genetics, 14 (3), e1007254. 10.1371/journal.pgen.1007254

Stankowski, S., Chase, M. A., Fuiten, A. M., Rodrigues, M. F., Ralph, P. L., & Streisfeld, M. A. (2019). Widespread selection and gene flow shape the genomic landscape during a radiation of monkeyflowers. PLOS Biology, 17 (7), e3000391. 10.1371/journal.pbio.3000391

Stevison, L. S., Woerner, A. E., Kidd, J. M., Kelley, J. L., Veeramah, K. R., McManus, K. F., Bustamante, C. D., Hammer, M. F., & Wall, J. D. (2016). The time scale of recombination rate evolution in great apes. Molecular Biology and Evolution, 33 (4), 928–945. 10.1093/molbev/msv331

Torres, R., Szpiech, Z. A., & Hernandez, R. D. (2018). Human demographic history has amplified the effects of background selection across the genome. PLOS Genetics, 14 (6), e1007387. 10.1371/journal.pgen.1007387

Turner, T. L., Hahn, M. W., & Nuzhdin, S. V. (2005). Genomic islands of speciation in anopheles gambiae. PLoS Biology, 3 (9). 10.1371/journal.pbio.0030285

van Doren, B. M., Campagna, L., Helm, B., Illera, J. C., Lovette, I. J., & Liedvogel, M. (2017). Correlated patterns of genetic diversity and differentiation across an avian family. Molecular Ecology, 26 (15), 3982–3997. 10.1111/mec.14083

Wang, J., Street, N. R., Park, E.-J., Liu, J., & Ingvarsson, P. K. (2020). Evidence for widespread selection in shaping the genomic landscape during speciation of populus. Molecular Ecology, 29 (6), 1120–1136. 10.1111/mec.15388

Williamson, R. J., Josephs, E. B., Platts, A. E., Hazzouri, K. M., Haudry, A., Blanchette, M., & Wright, S. I. (2014). Evidence for widespread positive and negative selection in coding and conserved noncoding regions of capsella grandiflora. PLoS Genetics, 10 (9). 10.1371/journal.pgen.1004622

Won, Y.-J., & Hey, J. (2005). Divergence population genetics of chimpanzees. Molecular Biology and Evolution, 22 (2), 297–307. 10.1093/molbev/msi017

Zhen, Y., Huber, C. D., Davies, R. W., & Lohmueller, K. E. (2021). Greater strength of selection and higher proportion of beneficial amino acid changing mutations in humans compared with mice and drosophila melanogaster. Genome Research, 31 (1), 110–120. 10.1101/gr.256636.119

